# Development of LYTACs via incorporating a nucleolin-targeting and lysosome-directed aptamer

**DOI:** 10.1101/2025.08.29.672993

**Authors:** Fang Qiu, Ziting Feng, Hongzhen Chen, Xiaoxuan Zhang, Jianmin Guo, Chunhao Cao, Aiping Lu, Chao Liang

## Abstract

Lysosomal targeting chimeras (LYTACs) represent an emerging class of bifunctional molecules that bridge extracellular target proteins with intrinsic lysosome-targeting receptors (LTRs) on the cell surface, facilitating endocytic internalization and subsequent lysosomal degradation of the targets. However, the therapeutic potential of LYTACs has been limited by the scarcity of suitable intrinsic LTRs. We previously identified an aptamer, SAPT8, that selectively targets nucleolin, a shuttling protein overexpressed on the surface of pathogenic FLSs in rheumatoid arthritis (RA), and induced its lysosomal degradation. In this study, we repurposed SAPT8 as a tumor-targeting and lysosome-directed ligand, leveraging the elevated expression of NCL on tumor cell surfaces. By conjugating SAPT8 with either the c-Met-binding aptamer SL1 or the small molecule inhibitor Tepotinib, we engineered novel LYTACs that demonstrated potent tumor-targeting capability and induced concurrent degradation of both c-Met and NCL, leading to significant antitumor effects. Furthermore, fusion of SAPT8 with VEGFR-2-targeting aptamer Apt02 generated LYTACs that simultaneously degraded VEGFR-2 and NCL, effectively suppressing RA-FLS activity. These results establish SAPT8 as a versatile platform for developing next-generation LYTACs, overcoming current limitations in extracellular protein degradation by circumventing dependence on endogenous LTRs.

**Graphic Abstract:** 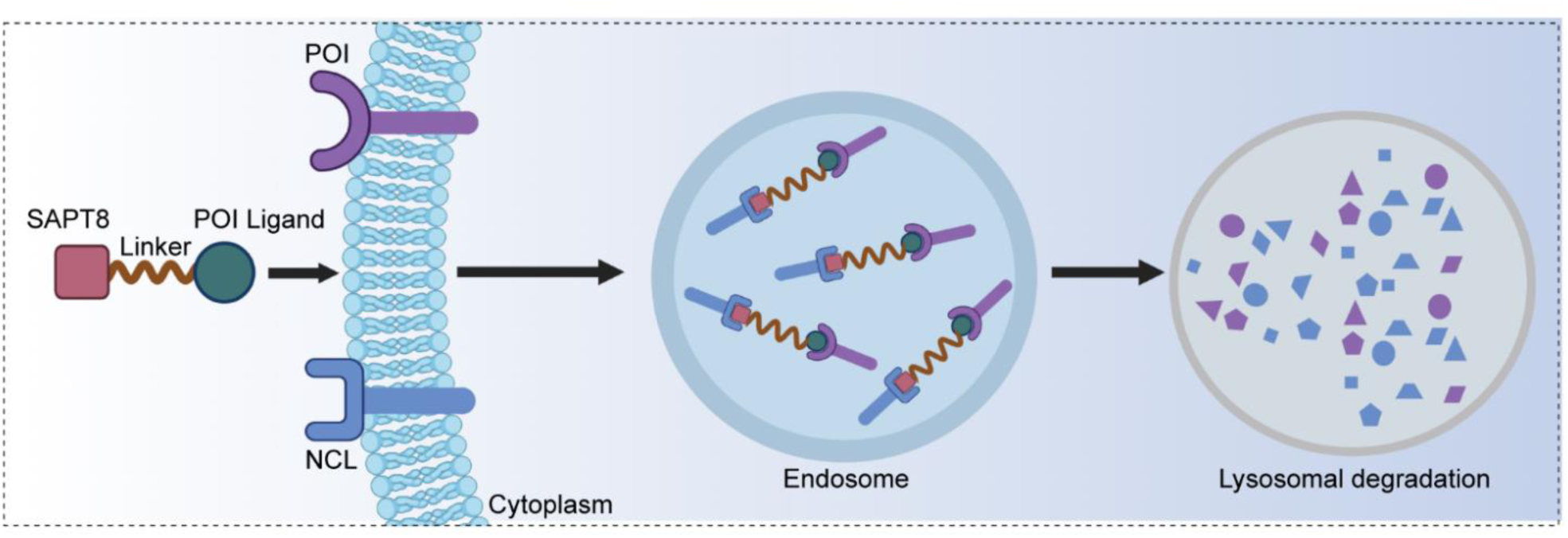

## Introduction

Targeted protein degradation (TPD) has revolutionized therapeutic strategies by enabling the selective elimination of proteins of interest (POIs) that are traditionally deemed “undruggable”[1–3]. This approach harnesses the cellular endogenous proteolytic machinery through chimeric degrader molecules that simultaneously bind both the POIs and degradation effectors[4]. The TPD field commenced with the creation of proteolysis-targeting chimeras (PROTACs) in 2001[5]. These heterobifunctional molecules induce the close proximity of POIs to E3 ligases, facilitating the ubiquitination and subsequent degradation of the POIs by the proteasome[6]. As a dominant modality in TPD, PROTACs have demonstrated significant advancements, with numerous candidates entering clinical trials[7–9]. However, PROTAC technology is inherently constrained to intracellular targets, excluding extracellular proteins from its therapeutic scope[10, 11].

Extracellular proteins constitute nearly half of the human proteome and represent a rich pool of therapeutic targets implicated in various human diseases[12]. The related TPD strategy for selective degradation of these proteins present in the extracellular space has only emerged in 2020 and is termed lysosomal targeting chimeras (LYTACs)[13, 14]. Conventional LYTACs are chimeric molecules that link extracellular POIs and intrinsic lysosome-targeting receptors (LTRs) on the cell surface[15]. The resulting complex is then internalized by endocytosis and trafficked to lysosomes, where the POIs are degraded[16]. Despite their potential, the current LYTAC toolbox remains limited, with the availability of only a few intrinsic LTRs, such as the pioneering cation-independent mannose 6-phosphate receptor (CI-M6PR), the latter employed asialoglycoprotein receptor (ASGPR), and a small number of other recently identified candidates[17–20]. To fully exploit the therapeutic potential of LYTACs, there is an urgent need to discover additional lysosome-directed ligands, thereby expanding the versatility of this emerging degradation platform[19].

Aptamers are oligonucleotides that specifically bind to their targets via folding intricate tertiary structures[21]. Traditionally, they are identified via systematic evolution of ligands by exponential enrichment (SELEX), enabling target recognition and broad application as targeting ligands for the precise delivery of therapeutics into cells[22]. Recently, we developed a novel cell-specific and cytotoxic SELEX (CSCT-SELEX) approach to generate unique aptamers with both cell-targeting capabilities and inherent therapeutic potential, without requiring prior knowledge of cell surface signatures[23]. Using this strategy, we identified a single-stranded DNA aptamer, SAPT8, which selectively targets pathogenic fibroblast-like synoviocytes (FLSs) in rheumatoid arthritis (RA) and induces their apoptosis, offering a new strategy for RA treatment[23]. We revealed that SAPT8 specifically binds to nucleolin (NCL), a shuttling protein abundantly expressed on the surface of pathogenic FLSs and implicated in their abnormal transformation in RA[23]. Mechanistically, SAPT8 acts as a lysosome-directed agent, and upon binding, it induces the internalization and subsequent lysosomal degradation of NCL, thus contributing to its therapeutic effects in RA[23]. Interestingly, beyond its aberrant expression on RA-FLSs, NCL has long been recognized as a surface protein overexpressed on tumor cells and functions as an oncoprotein[24–26]. Based on these findings, we hypothesize that SAPT8 could be repurposed for the development of tumor cell- or FLS-specific LYTACs.

To validate our hypothesis, we investigated the tumor-targeting ability and lysosomal trafficking of SAPT8 through its interaction with cell surface NCL. Furthermore, we engineered LYTACs by conjugating SAPT8 with specific ligands for c-mesenchymal-epithelial transition factor (c-Met). These chimeric molecules demonstrated efficient tumor-targeting capacity and facilitated the lysosomal degradation of both c-Met and NCL, resulting in significant tumor suppression. Additionally, we conjugated SAPT8 to a ligand targeting vascular endothelial growth factor receptor-2 (VEGFR-2), creating LYTACs that induced lysosome-mediated simultaneous degradation of both VEGFR-2 and NCL, leading to functional inhibition of RA-FLSs. Collectively, our findings highlighted the versatility of the SAPT8-based LYTAC platform for the knockdown of extracellular POIs.

## RESULTS

### NCL-dependent tumor specificity and lysosomal trafficking of SAPT8

We evaluated the tumor specificity of SAPT8 *in vitro* by incubating human osteosarcoma U-2 OS cells with SAPT8. SAPT8 exhibited significantly higher binding affinity compared to a negative control sequence (NC) (**Fig. 1A**). This binding was both time- and concentration-dependent (**Fig. 1B** and **1C**), with an equilibrium dissociation constant (*K_d_*) of 116.7 ± 34.56 nM (**Fig. 1D**). A biotin-streptavidin pull-down assay further confirmed that SAPT8, but not the NC, could specifically capture NCL in U-2 OS cells (**Fig. 1E**). When NCL expression was silenced in U-2 OS cells, SAPT8 binding was markedly decreased (**Fig. 1F**), suggesting that NCL was the cell-surface target of SAPT8. Additionally, we observed robust internalization of SAPT8 into U-2 OS cells, where it co-localized with a lysosome tracker (**Fig. 1G**). NCL silencing significantly reduced both internalization and lysosomal localization of SAPT8 (**Fig. 1H**), demonstrating that the cellular uptake and lysosomal trafficking of SAPT8 were mediated by NCL. These findings collectively establish SAPT8 as a promising tumor-specific and lysosome-directed aptamer.

**Fig. 1.**
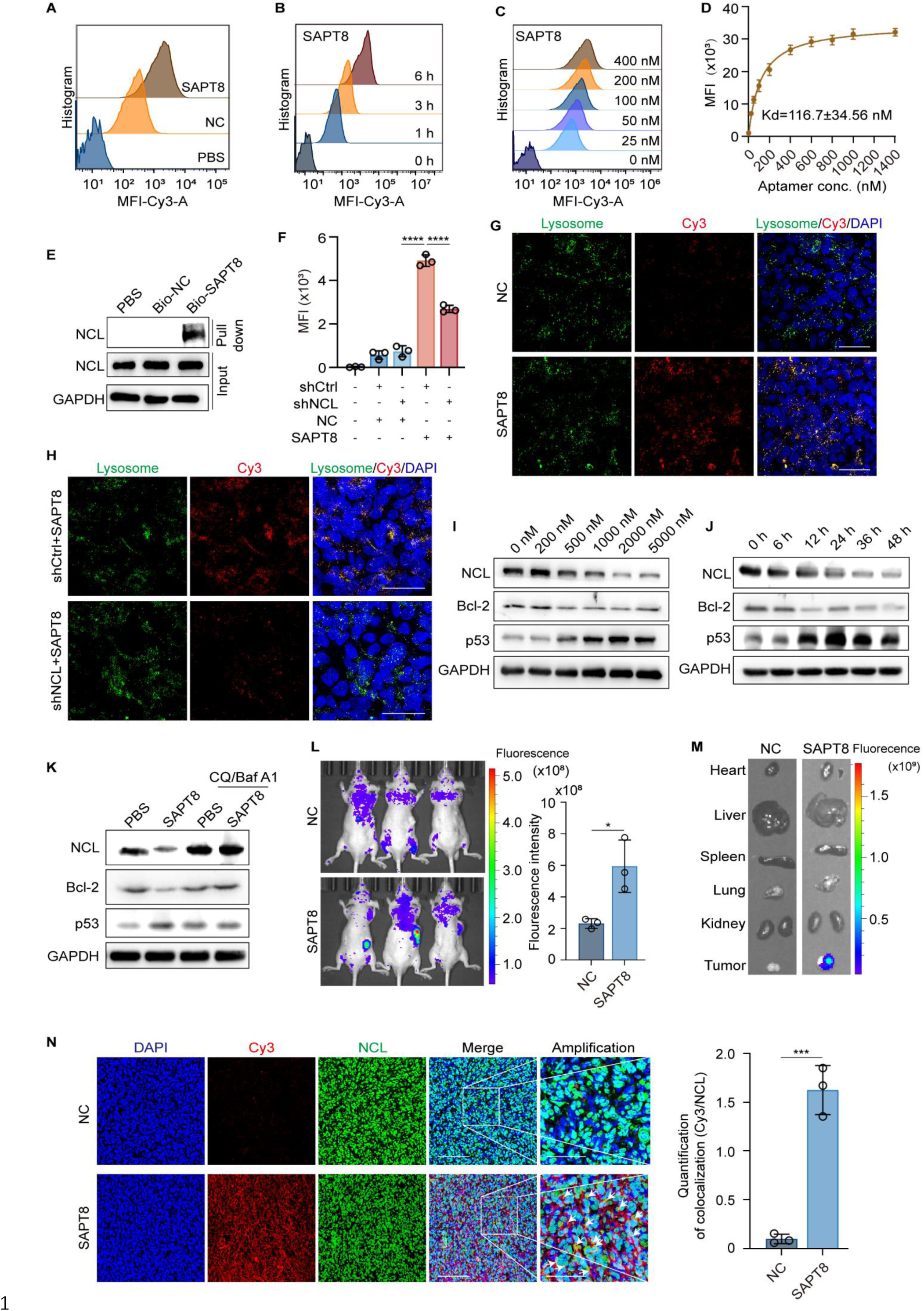
Tumor targeting and lysosomal localization of SAPT8. (**A**) Binding assay of 200 nM Cy3-labeled negative control (NC) or SAPT8 with U-2 OS cells after incubation for 3 h. (**B**) Binding assay of 200 nM Cy3-labeled SAPT8 with U-2 OS cells after incubation for different time points (0, 1, 3, and 6 h). (**C**) Binding assay of Cy3-labeled SAPT8 with U-2 OS cells after incubation for 3 h at different concentrations (0, 25, 50, 100, 200, and 400 nM). (**D**) The equilibrium dissociation constant (*K_d_*) of Cy3-labeled SAPT8 against U-2 OS cells. Cells were incubated with increasing concentrations (conc.) of SAPT8 for 3 h, and mean fluorescence intensity (MFI) was determined. *K_d_* value was calculated using nonlinear fitting analysis. (**E**) Pull-down assay for the interaction between biotin-labelled NC (Bio-NC) or SAPT8 (Bio-SAPT8) and NCL in U-2 OS cells after incubation for 8 h. (**F**) MFI of U-2 OS cells transfected with shRNA targeting NCL (shNCL) or negative control shRNA (shCtrl) after incubation with 200 nM NC and SAPT8 for 3 h. (**G**) Colocalization of 200 nM Cy3-labeled NC or SAPT8 (red) with LysoTracker (green) in U-2 OS cells after incubation for 6 h. Scale bars: 50 μm. (**H**) Colocalization of 200 nM Cy3-labeled NC or SAPT8 (red) with LysoTracker (green) in U-2 OS cells transfected with shNCL or shCtrl after incubation for 6 h. Scale bars: 50 μm. (**I**) Protein expression of NCL, Bcl-2, and p53 in U-2 OS cells treated with SAPT8 at different concentrations (0, 200, 500, 1000, 2000, and 5000 nM) for 24 h. (**J**) Protein expression of NCL, Bcl-2, and p53 in U-2 OS cells treated with 1000 nM SAPT8 for different time points (0, 6, 12, 24, 36, and 48 h). (**K**) Protein expression of NCL, Bcl-2, and p53 in U-2 OS cells treated with 1000 nM SAPT8 for 24 h in the presence or absence of lysosomal inhibitors CQ and Baf A1. (**L**) *In vivo* fluorescence imaging of nude mice bearing U-2 OS xenografts after tail-vein injection of 5 nmol Cy3-labeled NC or SAPT8 for 15 h. Left, representative biophotonic images. Right, quantification of the whole-body fluorescence. n = 3 mice per group. (**M**) *Ex vivo* fluorescence of Cy3-labeled NC or SAPT8 in major organs (heart, liver, spleen, lung, and kidney) and tumor tissues. (**N**) Colocalization of Cy3-labeled NC or SAPT8 with NCL in tumor tissues. Left, representative images. Right, quantification of colocalization. Scale bars: 100 µm. Each experiment was repeated three times. Data were represented as mean ± SD, and statistical significance was calculated by the one-way ANOVA with a post hoc test (F) or the two-tailed Student’s *t*-test (L and N). *\*p* < 0.05, *\*\*\*p* < 0.001*\*\*\*\*p* < 0.0001.

NCL is known to interact with the pro-apoptotic p53 mRNA at the 5′ untranslated region (5′-UTR) and prevent its translation, while also binding to the 3′-UTR of the anti-apoptotic B-cell lymphoma 2 (Bcl-2) mRNA to enhance its stability and protect it from nuclease degradation[23, 27, 28]. Through these mechanisms, NCL facilitates tumor cell evasion of apoptosis[27, 28]. We examined the levels of NCL and its downstream targets, Bcl-2 and p53, in U-2 OS cells following SAPT8 treatment. Our results demonstrated that SAPT8 reduced the expression of NCL, resulting in decreased Bcl-2 level and elevated p53 expression (**Fig. 1I** and **1J**). To explore whether these effects were lysosome-dependent, we treated U-2 OS cells with SAPT8 in the presence or absence of the lysosomal inhibitors Bafilomycin A1 (Baf A1) and chloroquine (CQ)[29]. Baf A1/CQ treatment reversed the SAPT8-induced degradation of NCL, as well as prevented the associated downregulation of Bcl-2 and upregulation of p53 (**Fig. 1K**). These findings indicate that SAPT8 specifically promotes lysosomal degradation of NCL, thereby modulating the expression of p53 and Bcl-2 in tumor cells.

### *In vitro* antitumor activity of SAPT8 via engaging NCL

Given that SAPT8 modulates the NCL-p53/Bcl-2 signaling axis, we evaluated its antitumor effects *in vitro*. Colony formation and EdU assays revealed that SAPT8 inhibited the proliferation of U-2 OS cells in a concentration-dependent manner (**Fig. S1A** and **S1B**). Terminal deoxynucleotidyl transferase dUTP nick end labeling (TUNEL) assay showed that SAPT8 treatment induced apoptosis in U-2 OS cells (**Fig. S1C**). However, upon silencing NCL expression, SAPT8 failed to inhibit U-2 OS cell growth or induce apoptosis (**Fig. S1D** and **S1E**), indicating that its antitumor activity is NCL-dependent. Moreover, SAPT8 was able to bind to human cervical carcinoma HeLa cells (**Fig. S2A**). In these cells, SAPT8 downregulated the expression of NCL and Bcl-2, while upregulating p53 level (**Fig. S2B**). Consistently, SAPT8 treatment inhibited proliferation and enhanced apoptosis of HeLa cells (**Fig. S2C-S2E**). Similarly, SAPT8 demonstrated high binding ability with human colorectal HCT116 cells, reduced NCL and Bcl-2 expression, increased p53 level, and inhibited proliferation and induced apoptosis of these cells (**Fig. S2F**-**S2J**). To evaluate the specificity of SAPT8, we investigated its effects on human normal epithelial MCF 10A cells *in vitro*. SAPT8 failed to bind to MCF 10A cells (**Fig. S3A**), and increasing concentrations of SAPT8 did not alter NCL, p53, or Bcl-2 levels (**Fig. S3B**), nor did it affect the proliferation of these cells (**Fig. S3C** and **S3D**). These findings indicate that SAPT8 selectively exerts therapeutic effects on tumor cells while sparing normal cells.

### *In vivo* tumor distribution and NCL colocalization of SAPT8

To evaluate the *in vivo* tumor-targeting ability of SAPT8, we established U-2 OS xenograft mouse models and administered a single intravenous injection of Cy3-labeled NC or SAPT8 for 15 h. *In vivo* whole-body imaging demonstrated robust and specific accumulation of SAPT8 at tumor sites, with fluorescence intensity significantly higher than that observed in the NC-treated group, which only displayed background-level signals (**Fig. 1L**). Subsequent *ex vivo* imaging further confirmed that SAPT8 preferentially accumulated in tumors compared to NC (**Fig. 1M**). Next, we examined whether SAPT8 colocalized with NCL within tumors *in vivo*. Confocal imaging showed strong aggregation of fluorescence signals within tumor tissues from SAPT8-injected mice, contrasting with barely detectable signals in NC-treated tumors (**Fig. 1N**). Importantly, SAPT8 was found to colocalize with NCL throughout the tumor tissue (**Fig. 1N**). These data suggest that SAPT8 possesses potent tumor-targeting ability *in vivo*, mediated by its specific interaction with NCL.

### Construction of SAPT8-SL1 LYTACs targeting c-Met

Building upon our identification of SAPT8 as a tumor-targeting and lysosome-directed aptamer, we sought to investigate its potential application in the development of LYTACs. c-Met is frequently overexpressed on the surface of tumor cells and plays a critical role in tumorigenesis[30, 31]. Before conjugating SAPT8 with ligands specific to c-Met, we conducted co-immunoprecipitation (co-IP) experiments, demonstrating that NCL did not physically interact with c-Met in either U-2 OS cells or HeLa cells (**Fig. S4A** and **S4B**). Consistently, the biotin-streptavidin pull-down assay showed that SAPT8 specifically captured NCL, but not c-Met, in both cells (**Fig. S4C** and **S4D**). To construct LYTAC degraders targeting c-Met, we engineered chimeric molecules by non-covalently linking the 5’ end of SAPT8 to either the 5’ or 3’ terminus of the c-Met-specific aptamer SL1[32] using a double-stranded DNA linker composed of six adenine-thymine (A-T) base pairs, resulting in two constructs, SAPT8-SL1-1 and SAPT8-SL1-2 (**Fig. 2A**). Successful assembly of these dual-aptamer chimeras was confirmed by non-denaturing polyacrylamide gel electrophoresis (PAGE), which revealed migration patterns consistent with their expected molecular weights (**Fig. S4E**). We anticipated that the SAPT8-SL1 chimeras would efficiently target NCL and c-Met on the surface of cancer cells and promote their co-transport to lysosomes for degradation, thereby inhibiting tumor progression (**Fig. 2B**).

**Fig. 2.**
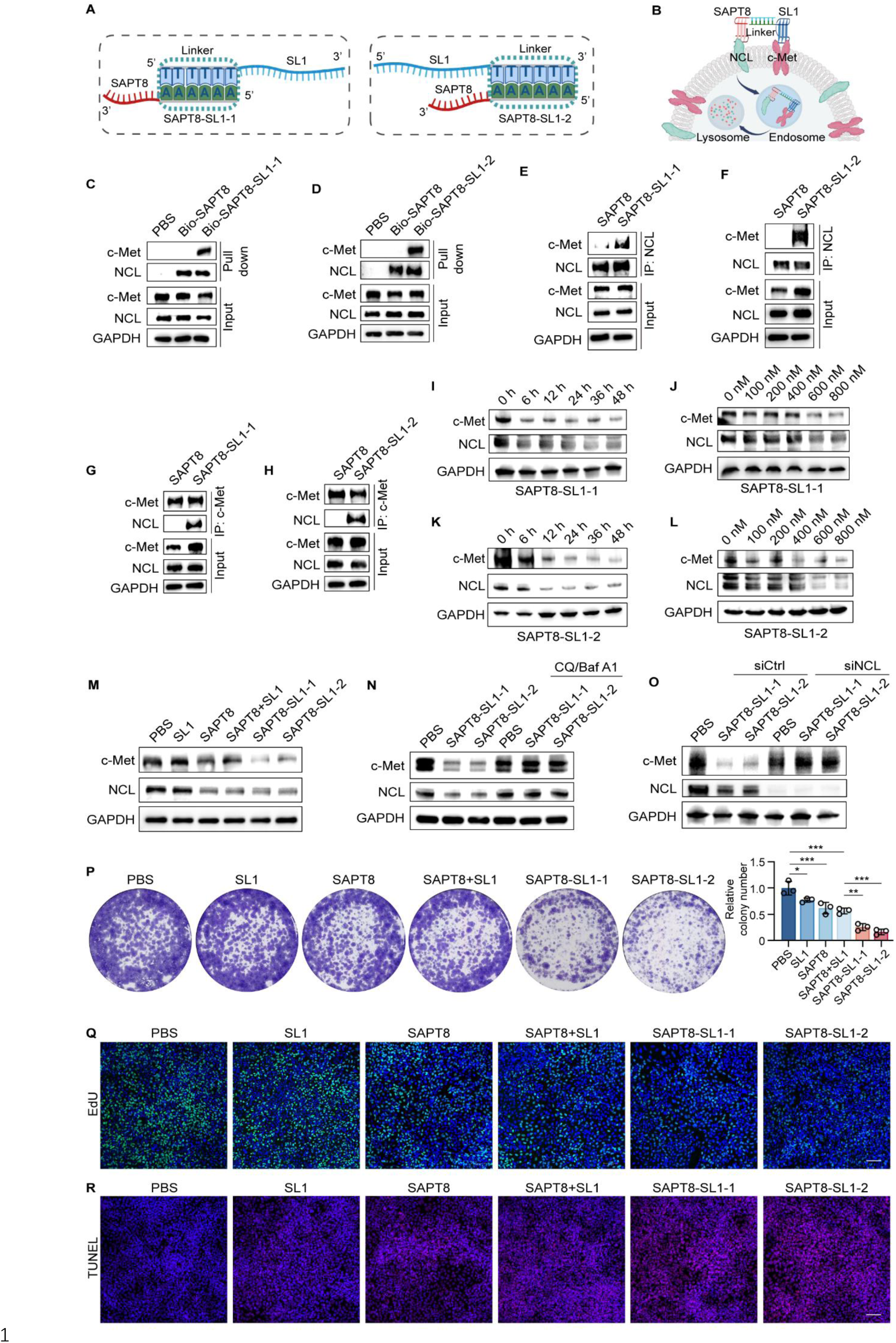
Degradation of c-Met and antitumor activity of SAPT8-SL1 LYTACs *in vitro*. (**A**) Schematic diagram showing the conjugation of SAPT8 with the 5’ or 3’ terminal of c-Met aptamer SL1 via a double-stranded DNA linker composed of six A-T base pairs, resulting in the generation of SAPT8-SL1-1 and SAPT8-SL1-2. (**B**) Schematic diagram illustrating the proposed mode of action of SAPT8-SL1 LYTACs. Briefly, SAPT8-SL1 LYTACs would selectively recognize tumor cells and then induce the internalization and lysosomal degradation of c-Met and NCL. (**C** and **D**) Pull-down assay of c-Met and NCL in U-2 OS cells after treatment with 1 μM biotin (Bio)-labeled SAPT8, SAPT8-SL1-1 (**C**), or SAPT8-SL1-2 (**D**) for 8 h. (**E** and **F**) Co-IP of c-Met and NCL in U-2 OS cells treated with 1 μM SAPT8, SAPT8-SL1-1 (**E**), or SAPT8-SL1-2 (**F**) for 8 h using an anti-NCL antibody. (**G** and **H**) Co-IP of c-Met and NCL in U-2 OS cells after treatment with 1 μM SAPT8, SAPT8-SL1-1 (**G**), or SAPT8-SL1-2 (**H**) for 8 h using an anti-c-Met antibody. (**I**) Protein expression of c-Met and NCL in U-2 OS cells after treatment with 600 nM SAPT8-SL1-1 for different time points (0, 6, 12, 24, 36, and 48 h). (**J**) Protein expression of c-Met and NCL in U-2 OS cells after treatment with SAPT8-SL1-1 at different concentrations (0, 100, 200, 400, 600, and 800 nM) for 12 h. (**K**) Protein expression of c-Met and NCL in U-2 OS cells after treatment with 600 nM SAPT8-SL1-2 for different time points (0, 6, 12, 24, 36, and 48 h). (**L**) Protein expression of c-Met and NCL in U-2 OS cells after treatment with SAPT8-SL1-2 at different concentrations (0, 100, 200, 400, 600, and 800 nM) for 12 h. (**M**) Protein expression of c-Met and NCL in U-2 OS cells after treatment with SL1, SAPT8, SAPT8+SL1, SAPT8-SL1-1, or SAPT8-SL1-2 at a concentration of 600 nM for 12 h. (**N**) Protein expression of c-Met and NCL in U-2 OS cells after treatment with SAPT8-SL1-1 or SAPT8-SL1-2 at a concentration of 600 nM for 12 h, with or without the presence of lysosomal inhibitors CQ and Baf A1. (**O**) Protein expression of c-Met and NCL in siCtrl- or siNCL-transfected U-2 OS cells after treatment with SAPT8-SL1-1 or SAPT8-SL1-2 at a concentration of 600 nM for 12 h. (**P**) Colony formation of U-2 OS cells after daily treatment with SL1, SAPT8, SAPT8+SL1, SAPT8-SL1-1, or SAPT8-SL1-2 at a concentration of 600 nM for 12 days. (**Q**) EdU assay for proliferation of U-2 OS cells after daily treatment with SL1, SAPT8, SAPT8+SL1, SAPT8-SL1-1, or SAPT8-SL1-2 at a concentration of 600 nM for 5 days. Nuclei were stained with DAPI. Scale bars: 100 μm. (**R**) TUNEL assay for apoptosis of U-2 OS cells after daily treatment with SL1, SAPT8, SAPT8+SL1, SAPT8-SL1-1, or SAPT8-SL1-2 at a concentration of 600 nM for 5 days. Nuclei were stained with DAPI. Scale bars: 100 µm. Each experiment was repeated three times. Data were represented as mean ± SD, and statistical significance was calculated by the one-way ANOVA with a post hoc test (P). *\*p* < 0.05, *\*\*p* < 0.01, *\*\*\*p* < 0.001.

### Lysosomal degradation of c-Met by SAPT8-SL1 LYTACs *in vitro*

We incubated U-2 OS cells with biotin-labeled SAPT8-SL1-1 or SAPT8-SL1-2, demonstrating that both chimeras effectively captured c-Met and NCL (**Fig. 2C** and **2D**). Furthermore, c-Met was co-precipitated with NCL (**Fig. 2E** and **2F**), and reciprocal co-precipitation of NCL with c-Met was observed in the presence of SAPT8-SL1-1 or SAPT8-SL1-2 (**Fig. 2G** and **2H**). To assess whether SAPT8-SL1-1 and SAPT8-SL1-2 could induce lysosomal degradation of c-Met, U-2 OS cells were treated with increasing incubation times and concentrations of these chimeras. The results showed a concomitant decrease in both c-Met and NCL protein levels in a time- and concentration-dependent manner **(Fig. 2I-2L)**. In contrast, treatment with SL1 alone, SAPT8 alone, or a physical mixture of SAPT8 and SL1 did not result in c-Met degradation **(Fig. 2M)**. We investigated whether the degradation of c-Met and NCL by SAPT8-SL1-1 and SAPT8-SL1-2 was indeed lysosome-dependent. U-2 OS cells were treated with these chimeras in the presence or absence of lysosomal inhibitors Chloroquine (CQ) and Bafilomycin A1 (Baf A1)[29]. Notably, CQ and Baf A1 inhibited the SAPT8-SL1-1- and SAPT8-SL1-2-mediated degradation of both c-Met and NCL **(Fig. 2N)**. Moreover, silencing NCL in U-2 OS cells impeded the ability of both chimeras to induce c-Met degradation **(Fig. 2O)**, indicating that SAPT8-SL1-mediated c-Met degradation is NCL-dependent. Since NCL is a shuttling protein that continually traffics between the nucleus, cytoplasm, and cell surface of tumor cells[33, 34], we evaluated NCL levels across different subcellular compartments. Interestingly, after incubation with SAPT8-SL1-2, membrane-associated NCL in U-2 OS cells showed a slight initial decrease followed by recovery, whereas the whole-cell NCL level was markedly reduced (**Fig. S5**). This observation suggests that intracellular pools of NCL can rapidly replenish membrane NCL following its internalization by LYTACs. In addition to U-2 OS cells, we demonstrated that SAPT8-SL1-1 or SAPT8-SL1-2 also induced concentration- and time-dependent degradation of c-Met and NCL in HeLa cells (**Fig. S6A-Fig. S6D**).

### Antitumor effects of SAPT8-SL1 LYTACs *in vitro*

We determined the *in vitro* antitumor potential of SAPT8-SL1 LYTACs targeting c-Met. U-2 OS cells were treated with PBS, SL1, SAPT8, a physical combination of SAPT8 and SL1 (SAPT8 + SL1), SAPT8-SL1-1, or SAPT8-SL1-2, followed by colony formation, EdU, and TUNEL assays. Compared to the PBS control, both SAPT8-SL1-1 and SAPT8-SL1-2 significantly inhibited the proliferation and induced apoptosis of U-2 OS cells (**Fig. 2P-2R**). Notably, SL1 alone, SAPT8 alone, and SAPT8 + SL1 also exhibited tumor inhibitory effects relative to PBS (**Fig. 2P-2R**), consistent with our above findings regarding the antitumor effects of SAPT8 and the reported antagonistic effect of SL1 on tumor growth[35]. However, the antitumor efficacy of SAPT8-SL1-1 and SAPT8-SL1-2 on U-2 OS cells was markedly superior to that of SL1 alone, SAPT8 alone, or their combination (**Fig. 2P-2R**). These results indicate that SAPT8-SL1 LYTACs, capable of simultaneously mediating degradation of both c-Met and NCL, offer enhanced therapeutic potency over strategies targeting NCL alone, c-Met alone, or both in combination. In addition, we incubated HeLa cells with PBS, SL1, SAPT8, SAPT8 + SL1, SAPT8-SL1-1, or SAPT8-SL1-2. HeLa cells exposed to SAPT8-SL1-1 or SAPT8-SL1-2 also demonstrated a pronounced reduction in proliferation and a significant increase in apoptosis compared to those treated with PBS, SL1, SAPT8, or the SAPT8 and SL1 combination (**Fig. S6E**-**S6G**).

### Tumor-targeting property of the SAPT8-SL1 LYTACs *in vivo*

To evaluate the tumor-targeting properties of SAPT8-SL1-1 and SAPT8-SL1-2 *in vivo*, U-2 OS-xenograft nude mice were intravenously injected with a single dose of Cy5-labeled SL1, SAPT8-SL1-1, or SAPT8-SL1-2 for 15 h. *In vivo* whole-body imaging revealed that fluorescence intensities at the tumor sites were significantly stronger for SAPT8-SL1-1 and SAPT8-SL1-2 than for SL1 alone (**Fig. 3A**). *Ex vivo* fluorescence imaging further demonstrated that both chimeras accumulated more efficiently in tumors compared to SL1 (**Fig. 3B**). Next, we examined whether SAPT8-SL1-1 and SAPT8-SL1-2 colocalized with c-Met or NCL within U-2 OS tumor tissues *in vivo*. Confocal imaging showed prominent aggregation of fluorescence signals in tumors from mice injected with SAPT8-SL1-1 or SAPT8-SL1-2, whereas minimal signals were detected in tumors from SL1-treated mice (**Fig. 3C** and **3D**). Importantly, both chimeras colocalized with c-Met and NCL in tumor tissues (**Fig. 3C** and **3D**).

**Fig. 3.**
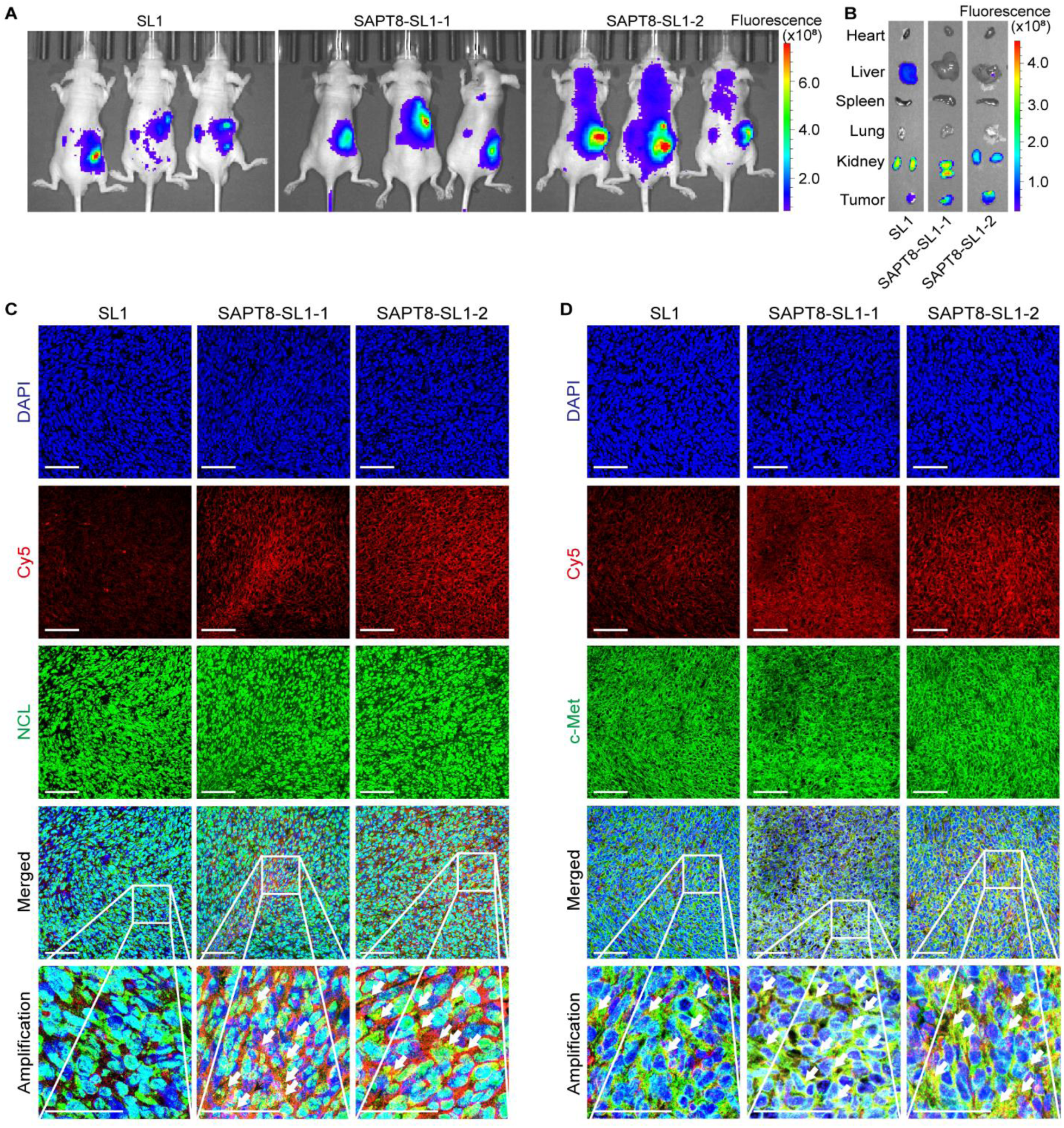
Tissue distribution of SAPT8-SL1 LYTACs *in vivo*. (**A**) *In vivo* whole-body fluorescence imaging of BALB/c nude mice bearing U-2 OS xenografts after tail-vein injection of 5 nmol Cy5-labeled SL1, SAPT8-SL1-1, or SAPT8-SL1-2 for 15 h. (**B**) *Ex vivo* fluorescence of Cy5-labeled SL1, SAPT8-SL1-1, or SAPT8-SL1-2 in major organs (heart, liver, spleen, lung, and kidney) and tumor tissues. (**C** and **D**) Immunofluorescence staining showing *in vivo* colocalization of Cy5-labeled SL1, SAPT8-SL1-1, or SAPT8-SL1-2 with NCL (**C**) or c-Met (**D**) in tumor tissues. Scale bars: 100 µm. n = 3 mice per group.

### Antitumor potency of SAPT8-SL1 LYTACs in an orthotopic xenograft model

We established an orthotopic U-2 OS xenograft mouse model to evaluate the therapeutic efficacy and safety of SAPT8-SL1-1 and SAPT8-SL1-2 *in vivo*. Tumor-bearing mice were intravenously administered PBS, SL1, SAPT8, SAPT8 + SL1, SAPT8-SL1-1, or SAPT8-SL1-2 every two days over a 10-day period (**Fig. 4A**). Both SAPT8-SL1-1 and SAPT8-SL1-2 significantly suppressed orthotopic tumor growth in the tibial bone compared with PBS treatment (**Fig. 4B** and **4C**). Micro-computed tomography (μCT) showed reduced tumor-associated bone destruction in mice treated with SAPT8-SL1-1 or SAPT8-SL1-2 (**Fig. 4D** and **4E**). Hematoxylin and eosin (H&E) staining indicated a notable increase in apoptotic cells within tumors from mice treated with these chimeras (**Fig. 4F**). Levels of NCL and c-Met in tumor tissues were significantly decreased in the SAPT8-SL1-1 and SAPT8-SL1-2 groups compared to the PBS group (**Fig. 4G** and **4H**). Treatment with SAPT8-SL1 chimeras also resulted in fewer Ki-67-positive proliferative cells and more TUNEL-positive apoptotic cells in tumors (**Fig. 4I** and **4J**). Although SL1, SAPT8, and SAPT8 + SL1 individually exhibited antitumor effects, their efficacy was less pronounced than that observed with SAPT8-SL1-1 or SAPT8-SL1-2 (**Fig. 4B-4J**). Neither SAPT8-SL1-1 nor SAPT8-SL1-2 induced significant changes in the body weight among treated mice (**Fig. S7A**). Serum biochemical analyses indicated no alteration in liver or kidney function parameters, including uric acid (UA), creatinine (CR), alanine aminotransferase (ALT), alkaline phosphatase (ALP), blood urea nitrogen (BUN), or total protein (TP) following treatment with the chimeras (**Fig. S7B**-**S7G**). Histological evaluation confirmed the absence of major organ damage in mice receiving SAPT8-SL1 chimeras (**Fig. S7H**).

**Fig. 4.**
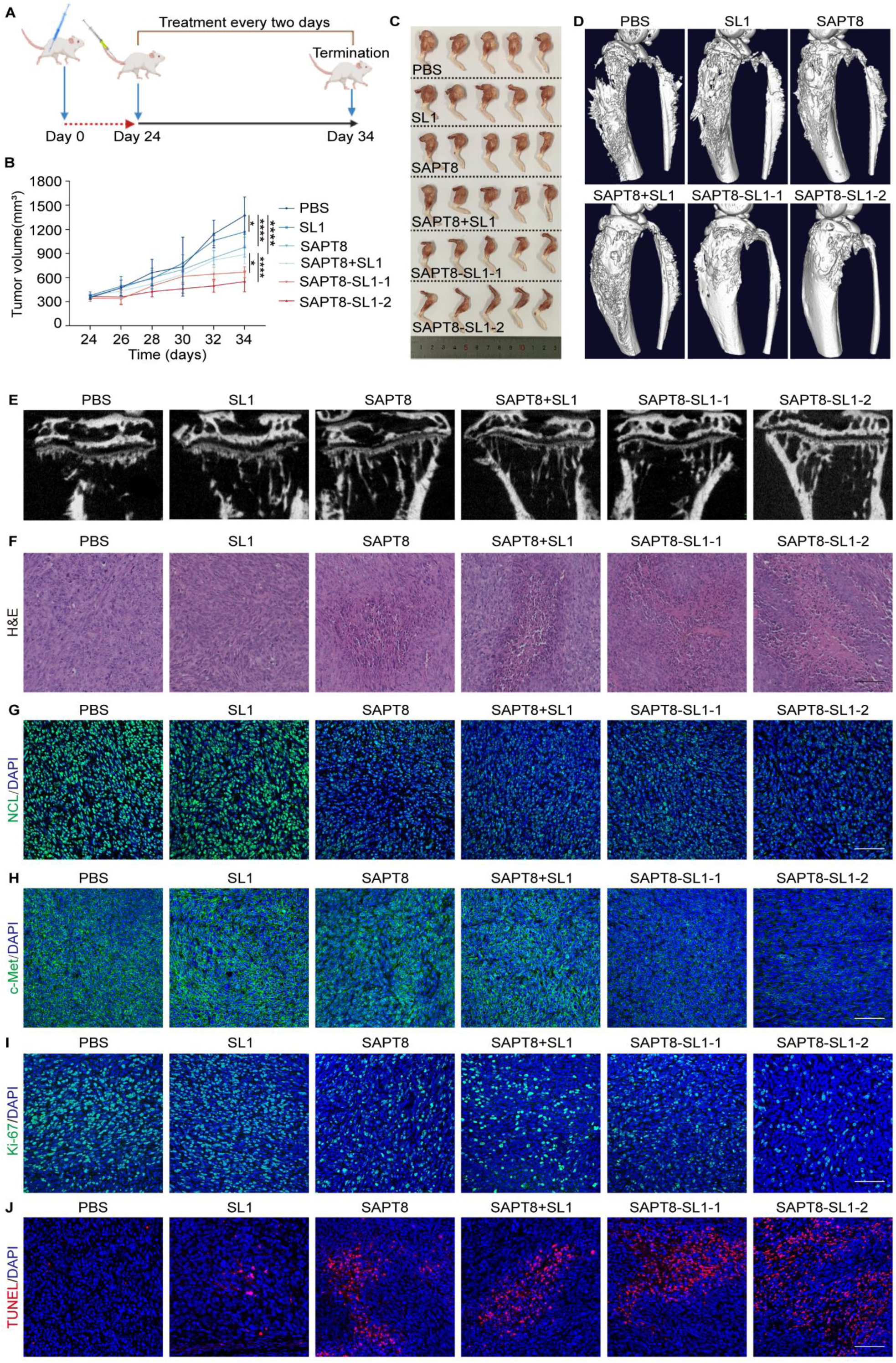
Antitumor efficacy of SAPT8-SL1 LYTACs in an orthotopic xenograft mouse model. (**A**) Schematic of experimental design for evaluating SAPT8-SL1 efficacy in an orthotopic xenograft mouse model. Briefly, SCID mice were inoculated with 6×10^5^ U-2 OS cells via intra-bone marrow injections. After 24 days, mice bearing orthotopic U-2 OS xenografts were intravenously injected with SL1, SAPT8, SAPT8+SL1, SAPT8-SL1-1, or SAPT8-SL1-2 for 10 days at a dose of 6 μmol/kg every two days. (**B**) Tumor volume in SCID mice from each treatment group. (**C**) Images of orthotopic xenografts from SCID mice. (**D**) Representative μCT images showing the proximal tibia of tumor-bearing mice in each treatment group. (**E**) Tumor-associated bone destruction in each treatment group. (**F**) H&E staining of the orthotopic tumor sections. Scale bars: 100 μm. (**G-I**) Immunofluorescence staining showing expression of NCL (**G**), c-Met (**H**), and Ki-67 (**I**) in tumor sections. (**J**) TUNEL assay for detecting apoptotic cells in the tumor sections. Scale bars: 50 μm, n = 5 for each group. Data were represented as mean ± SD, and statistical significance was calculated by the two-way ANOVA with a post hoc test (B). *\*p* < 0.05, *\*\*\*\*p* < 0.0001.

### Antitumor activity of SAPT8-SL1 LYTACs in a subcutaneous xenograft model

To further validate the therapeutic efficacy and safety profile of SAPT8-SL1-1 and SAPT8-SL1-2, we employed a subcutaneous U-2 OS xenograft mouse model. Mice bearing U-2 OS xenografts received tail-vein injection of PBS, SL1, SAPT8, SAPT8 + SL1, SAPT8-SL1-1, or SAPT8-SL1-2 every two days over 12-day period (**Fig. S8A**). The average tumor volumes in mice treated with SAPT8-SL1-1 or SAPT8-SL1-2 were significantly smaller than those in the PBS group (**Fig. S8B** and **S8C**). While SL1, SAPT8, and SAPT8 + SL1 also resulted in reduced tumor growth compared to PBS, SAPT8-SL1-1 and SAPT8-SL1-2 exhibited superior efficacy (**Fig. S8B** and **S8C**). H&E staining revealed a marked increase in apoptotic cells within tumors from mice treated with the SAPT8-SL1 chimeras compared to other groups (**Fig. S8D**). Both SAPT8-SL1-1 and SAPT8-SL1-2 effectively reduced levels of NCL and c-Met in tumors (**Fig. S8E** and **S8F**) and decreased the number of Ki-67-positive proliferative cells while increasing TUNEL-positive apoptotic cells (**Fig. S8G** and **S8H**). Neither SAPT8-SL1-1 nor SAPT8-SL1-2 induced significant changes in body weight among the treated mice (**Fig. S9A**). Liver and kidney function parameters (UA, CR, ALT, ALP, BUN, and TP) remained within normal ranges following treatment with SAPT8-SL1-1 or SAPT8-SL1-2 (**Fig. S9A**-**S9G**). Histopathological examination of major organs confirmed the absence of treatment-related toxicity caused by SAPT8-SL1 chimeras (**Fig. S9H**).

### Synthesis of SAPT8-Tep LYTACs

To demonstrate the versatility of the SAPT8-based LYTAC platform, we conjugated SAPT8 to the small molecule c-Met inhibitor Tepotinib (Tep)[36] via covalent linkers, including alkyl linkers of varying lengths and a polyethylene glycol (PEG) linker. This approach resulted in the creation of three aptamer-small molecule hybrid LYTAC chimeras: SAPT8-Tep-1, SAPT8-Tep-2, and SAPT8-Tep-3 (**Fig. 5A** and **S10A-S10C**). Detailed synthetic routes were provided in the **Supplementary Methods**. The structures and purities of the key intermediates and all three SAPT8-Tep chimeras were confirmed using ^1^H-NMR, ultra-performance liquid chromatography-mass spectrometry (UPLC-MS), liquid chromatography-mass spectrometry (LC-MS), and high-performance liquid chromatography (HPLC) (**Supplementary Methods**). We hypothesized that these SAPT8-Tep chimeras would enable dual targeting of NCL and c-Met on the surfaces of cancer cells and promote the lysosomal degradation of c-Met together with NCL, thereby offering a promising antitumor approach (**Fig. 5B**).

**Fig. 5.**
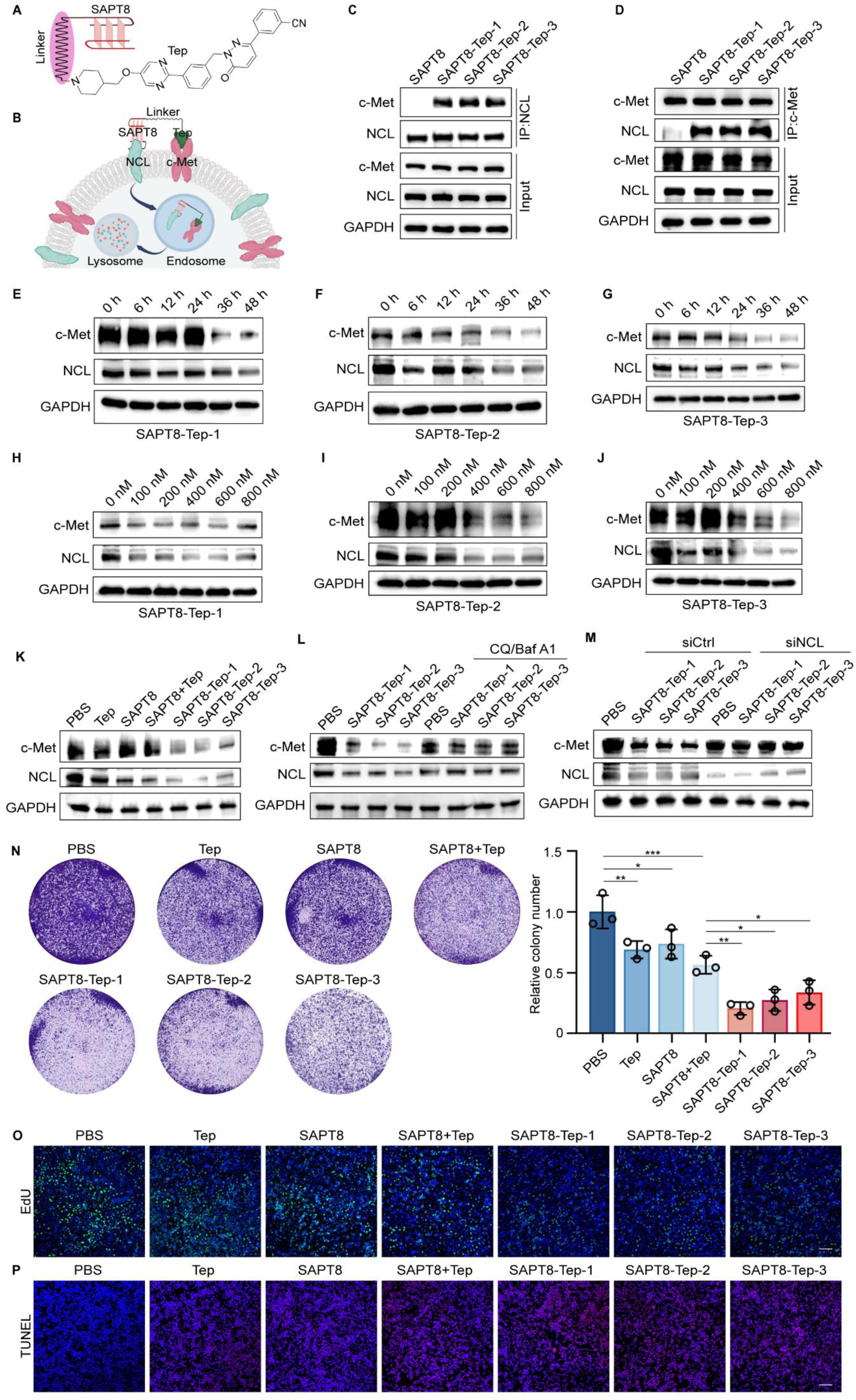
Degradation of c-Met and antitumor activity of SAPT8-Tep LYTACs *in vitro*. (**A**) Schematic diagram showing the conjugation of SAPT8 with c-Met small molecule inhibitor Tep. (**B**) Schematic diagram illustrating the proposed mode of action of SAPT8-Tep LYTACs. Briefly, SAPT8-Tep LYTACs would selectively recognize tumor cells and then induce the internalization and lysosomal degradation of c-Met and NCL. (**C** and **D**) Co-IP of NCL and c-Met in HeLa cells treated with 1 μM SAPT8, SAPT8-Tep-1, SAPT8-Tep-2, or SAPT8-Tep-3 for 8 h using an anti-NCL (**C**) or an anti-c-Met antibody (**D**). (**E-G**) Protein expression of c-Met and NCL in HeLa cells after treatment with SAPT8-Tep-1 (**E**), SAPT8-Tep-2 (**F**), or SAPT8-Tep-3 (**G**) at a concentration of 600 nM for different time points (0, 6, 12, 24, 36, and 48 h). (**H-J**) Protein expression of c-Met and NCL in HeLa cells after treatment with SAPT8-Tep-1 (**H**), SAPT8-Tep-2 (**I**), or SAPT8-Tep-3 (**J**) at different concentrations (0, 100, 200, 400, 600, and 800 nM) for 36 h. (**K**) Protein expression of c-Met and NCL in HeLa cells after treatment with Tep, SAPT8, SAPT8+Tep, SAPT8-Tep-1, SAPT8-Tep-2, or SAPT8-Tep-3 at a concentration of 600 nM for 36 h. (**L**) Protein expression of c-Met and NCL in HeLa cells after treatment with SAPT8-Tep-1, SAPT8-Tep-2, or SAPT8-Tep-3 at a concentration of 600 nM for 36 h, with or without the presence of lysosomal inhibitors CQ and Baf A1. (**M**) Protein expression of c-Met and NCL in siCtrl- or siNCL-transfected HeLa cells after treatment with SAPT8-Tep-1, SAPT8-Tep-2, and SAPT8-Tep-3 at a concentration of 600 nM for 36 h. (**N**) Colony formation of HeLa cells after treatment with Tep, SAPT8, SAPT8+Tep, SAPT8-Tep-1, SAPT8-Tep-2, and SAPT8-Tep-3 at a concentration of 600 nM for 7 days. (**O**) EdU assay for proliferation of HeLa cells after daily treatment with Tep, SAPT8, SAPT8+Tep, SAPT8-Tep-1, SAPT8-Tep-2, or SAPT8-Tep-3 at a concentration of 600 nM for 5 days. Nuclei were stained with DAPI. Scale bars: 100 μm. (**P**) TUNEL assay for apoptosis of HeLa cells after daily treatment with Tep, SAPT8, SAPT8+Tep, SAPT8-Tep-1, SAPT8-Tep-2, and SAPT8-Tep-3 at a concentration of 600 nM for 5 days. Nuclei were stained with DAPI. Scale bars = 100 µm. Each experiment was repeated three times. Data were represented as mean ± SD, and statistical significance was calculated by the one-way ANOVA with a post hoc test (N). *\*p* < 0.05, *\*\*p* < 0.01, *\*\*\*p* < 0.001.

### Lysosomal degradation of c-Met and antitumor activity of SAPT8-Tep LYTACs *in vitro*

Upon incubating HeLa cells with SAPT8-Tep-1, SAPT8-Tep-2, and SAPT8-Tep-3, we showed co-precipitation of c-Met with NCL, and vice versa (**Fig. 5C** and **5D**). To assess their degradation ability, we treated HeLa cells with each chimera and found that SAPT8-Tep-1, SAPT8-Tep-2, and SAPT8-Tep-3 significantly decreased the levels of both proteins in a time- and concentration-dependent manner **(Fig. 5E**–**Fig. 5J)**. Comparatively, HeLa cells treated with PBS, Tep, SAPT8, or SAPT8 + Tep did not exhibit c-Met degradation, whereas all three SAPT8-Tep chimeras facilitated simultaneous degradation of both c-Met and NCL **(Fig. 5K)**. Moreover, the reduction of c-Met induced by SAPT8-Tep chimeras was effectively blocked by lysosomal inhibitors CQ and Baf A1 **(Fig. 5L)**, highlighting the lysosomal dependency of this process. NCL silencing abolished SAPT8-Tep-mediated c-Met degradation **(Fig. 5M)**. When compared with SAPT8-SL1 LYTACs, the SAPT8-Tep chimeras exhibited superior c-Met degradation efficiency (**Fig. S11**). *In vitro* antitumor evaluation using colony formation and EdU assays revealed that SAPT8-Tep-1, SAPT8-Tep-2, and SAPT8-Tep-3 markedly inhibited HeLa cell proliferation, with their antiproliferative impacts surpassing those of Tep, SAPT8, or their combination (**Fig. 5N** and **Fig. 5O**). Consistently, TUNEL assays confirmed that SAPT8-Tep chimeras induced a significantly higher level of apoptosis compared to other treatments (**Fig. 5P**).

In addition to HeLa cells, we also demonstrated time- and concentration-dependent degradation of c-Met and NCL induced by SAPT8-Tep-1, SAPT8-Tep-2, or SAPT8-Tep-3 in human tumorigenic lung epithelial A549 cells (**Fig. S12A-S12F**). A549 cells were treated with PBS, Tep, SAPT8, SAPT8 + Tep, SAPT8-Tep-1, SAPT8-Tep-2, or SAPT8-Tep-3. The chimeras induced simultaneous degradation of both c-Met and NCL, whereas SAPT8 alone, Tep alone, or their combination exhibited no c-Met degradation activity **(Fig. S12G)**. To assess their antitumor potential, colony formation and EdU assays showed that SAPT8-Tep-1, SAPT8-Tep-2, and SAPT8-Tep-3 more effectively inhibited the proliferation of A549 cells than SAPT8 alone, Tep alone, or their combination (**Fig. S12H** and **S12I**). The TUNEL assay demonstrated that these chimeras induced more pronounced apoptosis in A549 cells (**Fig. S12J**). Furthermore, the SAPT8-Tep chimeras also mediated degradation of c-Met and NCL in U-2 OS cells (**Fig. S13A-S13C**).

### Antitumor activity of SAPT8-Tep LYTACs in a subcutaneous xenograft model

We selected SAPT8-Tep-1 for evaluating the therapeutic efficacy and safety of SAPT8-Tep LYTACs *in vivo* using a HeLa-xenografted mouse model. Mice bearing tumor xenografts received intravenous treatments with PBS, Tep, SAPT8, SAPT8 + Tep, or SAPT8-Tep-1 every two days over a 12-day period (**Fig. 6A**). Our findings revealed that the average tumor volume in SAPT8-Tep-1-treated mice was smaller than those in the PBS, Tep, SAPT8, or SAPT8 + Tep groups (**Fig. 6B-6C**). H&E staining revealed a substantial increase in apoptotic cells within the tumors of SAPT8-Tep-1-treated mice when compared to other treatments (**Fig. 6D**). Tumor levels of NCL and c-Met were considerably lowered in the SAPT8-Tep-1 group (**Fig. 6E** and **6F**). SAPT8-Tep-1 treatment also resulted in a decrease in Ki-67-positive proliferative cells and an increase in TUNEL-positive apoptotic cells within tumors (**Fig. 6G** and **6H**). Although SL1, SAPT8, and SAPT8 + SL1 also contributed to reduced tumor growth compared to the PBS group, SAPT8-SL1-1 and SAPT8-SL1-2 demonstrated superior efficacy (**Fig. 6B** and **6H**). Notably, SAPT8-Tep-1 did not cause significant changes in body weight among tumor-bearing mice (**Fig. S14A**). Serum biochemical analyses showed no significant alterations in liver or kidney function parameters (UA, CR, ALT, ALP, BUN, and TP) in the SAPT8-Tep-1 group (**Fig. S14B**-**S14G**). Histological evaluations verified that SAPT8-Tep-1 did not induce major organ damage (**Fig. S14H**).

**Fig. 6.**
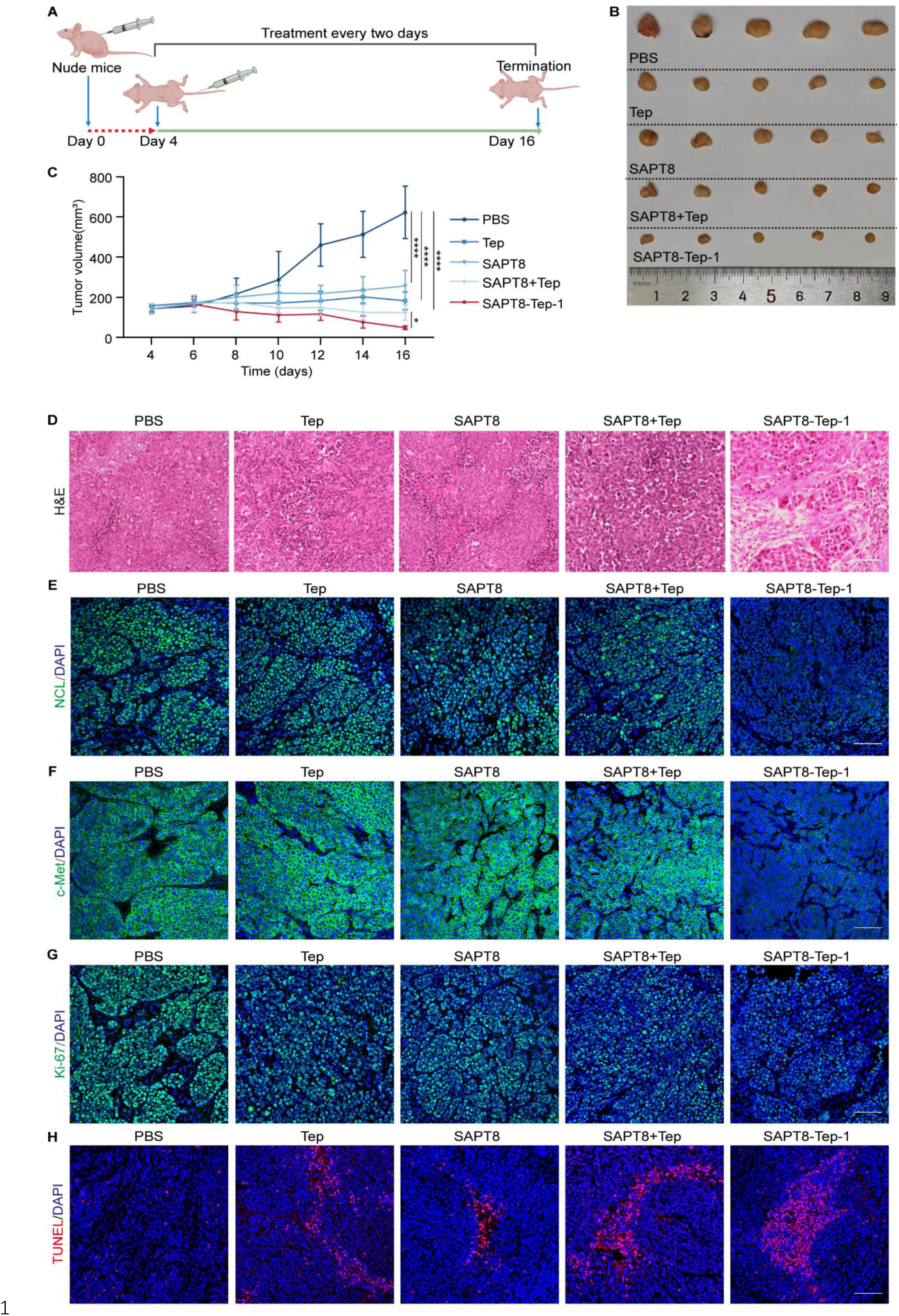
Antitumor efficacy of SAPT8-Tep LYTACs in a subcutaneous xenograft mouse model. (**A**) Schematic of the experimental design for evaluating the antitumor efficacy of SAPT8-Tep-1 in a subcutaneous xenograft mouse model. Briefly, BALB/c nude mice were subcutaneously injected with 5×10^6^ HeLa cells. After 4 days, mice bearing xenograft tumors were intravenously treated with Tep, SAPT8, SAPT8+Tep, and SAPT8-Tep-1 at a dose of 6 μmol/kg every two days for 12 days. (**B**) Images of the xenografted tumors from each treatment group. (**C**) Tumor volume of the nude mice in each treatment group. (**D**) H&E staining of tumor sections from each treatment group. Scale bars: 100 μm. (**E**-**G**) Immunofluorescence staining of tumor sections for NCL (**E**), c-Met (**F**), and Ki-67 (**G**) expression in each treatment group. (**H**) TUNEL assay detecting apoptotic cells in tumor sections from each treatment group. Scale bars: 100 μm, n = 5 for each group. Data were represented as mean ± SD, and statistical significance was calculated by the two-way ANOVA (C) with a post hoc test. *\*p* < 0.05, *\*\*\*\*p* < 0.0001.

### Construction of SAPT8-Apt02 LYTACs targeting VEGFR-2

Since SAPT8 was initially screened to target pathogenic FLSs in RA via its interaction with NCL and trigger lysosomal degradation of NCL[23], we attempted to develop LYTACs for inhibiting the pathogenic transformation of these RA-FLSs. Given that RA-FLSs overexpress VEGFR-2, which triggers their survival and migration[37–39], we designated it as the POI for LYTAC development. We first confirmed that NCL does not physically interact with VEGFR-2 in RA-FLSs via co-IP assays (**Fig. S15A**). Furthermore, we demonstrated that SAPT8 specifically captured NCL, but not VEGFR-2, by the biotin-streptavidin pull-down assay (**Fig. S15B**). We engineered chimeric molecules by covalently conjugating SAPT8 to either the 5’ or 3’ terminus of the VEGFR-2-specific aptamer Apt02[40] using a single-stranded DNA linker composed of six T bases, yielding SAPT8-Apt02-1 and SAPT8-Apt02-2 (**Fig. S15C**). We hypothesized that these bifunctional chimeras would shuttle both VEGFR-2 and NCL to lysosomes for degradation, thereby inhibiting FLS pathogenicity in RA (**Fig. S15D**). Successful construction of both chimeras was confirmed by mass spectrometric analysis (**Fig. S15E-S15F**).

### Lysosomal degradation of VEGFR-2 and antitumor effects of SAPT8-Apt02 LYTACs *in vitro*

To evaluate SAPT8-Apt02 LYTAC effects on VEGFR-2 expression, RA-FLSs were incubated with biotin-labeled SAPT8-Apt02-1 or SAPT8-Apt02-2. Both constructs effectively captured VEGFR-2 and NCL (**Fig. S15G**). VEGFR-2 was co-precipitated with NCL and vice versa in the presence of SAPT8-Apt02-1 or SAPT8-Apt02-2 (**Fig. S15H** and **S15I**). Degradation kinetics analysis revealed that treatment with increasing concentrations or incubation times of SAPT8-Apt02-1 and SAPT8-Apt02-2 induced synchronous, dose- and time-dependent reduction of both VEGFR-2 and NCL **(Fig. S15J-S15M)**. This degradation was specific to the chimeric constructs, as neither Apt02 alone, SAPT8 alone, nor a SAPT8+SL1 physical mixture affected VEGFR-2 level **(Fig. S15N)**. Lysosomal inhibitors CQ and Baf A1 were able to block SAPT8-Apt02-1- and SAPT8-Apt02-2-mediated degradation of VEGFR-2 and NCL **(Fig. S15O)**. Moreover, silencing NCL in RA-FLSs impeded the ability of SAPT8-Apt02-1 and SAPT8-Apt02-2 to induce VEGFR-2 degradation **(Fig. S15P)**. We also analyzed the phenotypes of RA-FLSs after treatment with SAPT8-Apt02 LYTACs for VEGFR-2. RA-FLSs were treated with PBS, Apt02, SAPT8, SAPT8 + Apt02, SAPT8-Apt02-1, or SAPT8-Apt02-2 and then subjected to TUNEL and migration assays. Consistent with our previous findings, SAPT8 alone attenuated the aggressive phenotype of RA-FLSs when compared to PBS[23]. However, SAPT8-Apt02-1 and SAPT8-Apt02-2 induced significantly greater levels of apoptosis and more effectively inhibited the migration of RA-FLSs (**Fig. S15Q-S15R**).

## DISCUSSION

To date, LYTAC and PROTAC have established themselves as the leading technologies in the PTD field[41, 42]. Although both utilize a similar bifunctional molecular architecture, they are not functionally redundant: PROTACs facilitate the proteosome-dependent degradation of intracellular POIs, while LYTACs are tailored to eliminate extracellular POIs via the lysosomal pathway[4]. This complementarity allows for comprehensive coverage of the whole proteome as druggable targets[43]. However, clinical translation faces analogous challenges for both platforms. A major bottleneck for PROTACs is the limited repertoire of available E3 ligases and their cognate ligands, despite over 600 E3 ligases being encoded in the human genome[44]. Similarly, LYTAC development is hampered by the scarcity of LTRs and their corresponding ligands[17–20]. The current LYTAC platform has evolved through successive generations, with first-generation designs relying on the CI-M6PR and second-generation designs employing the ASGPR. Recent efforts have expanded the LYTAC toolbox to include other LTR candidates, such as transferrin receptor (TfR), glucose transporters (GLUTs), CXC chemokine receptor 7 (CXCR7), glucagon-like peptide 1 receptor (GLP-1R), folate receptor (FR), sortilin, and integrins[17–20]. However, most of these LTRs are intrinsic, posing two major limitations. First, their constitutive expression across diseased and normal tissues increases the risk of on-target, off-tissue effects, often necessitating targeted delivery systems to minimize toxicity[43, 45]. Second, certain LTRs, such as ASGPR and IGF2R, can be occupied by their endogenous ligands, further reducing their efficacy in mediating POI degradation[12].

In this study, we successfully repurposed the aptamer SAPT8, initially screened through our CSCT-SELEX strategy for targeting RA-FLSs[23], for specific tumor cell binding. This repurposing was enabled by the shared overexpression of NCL, the molecular target of SAPT8, on the surfaces of both cell types[23, 46]. Notably, the NCL overexpression on pathogenic FLSs further elucidates their acquisition of tumor-like characteristics during RA progression, including hyperproliferation, apoptosis resistance, and enhanced invasive and migratory capacities[23, 47]. Through *in vitro* and *in vivo* validation, we confirmed the tumor-targeting specificity of SAPT8. Mechanistic studies revealed that SAPT8 binding to surface NCL on tumor cells exhibited lysosomal trafficking ability, mirroring its previously observed behavior in RA-FLSs[23]. Leveraging this property, we developed a SAPT8-based LYTAC platform that effectively degraded c-Met on tumor cells and VEGFR-2 on RA-FLSs. A key advantage of this platform is its exceptional selectivity: as normal cells lack surface NCL expression[48], SAPT8-based LYTACs did not degrade c-Met in these cells and exhibited favorable safety profiles. This inherent specificity effectively eliminates concerns about on-target, off-tissue toxicity of SAPT8-based LYTACs. Moreover, as NCL is not an intrinsic LTR, potential competition from endogenous ligands is negligible, ensuring unimpeded efficacy of the SAPT8-based LYTAC platform.

The intrinsic LTRs are presumed to undergo clathrin- or caveolae-mediated endocytosis followed by routing through early and late endosomes to lysosomes[49–52]. The acidic environment within these compartments theoretically facilitates the dissociation of the LTR from its cargo, allowing them to recycle back to the cell membrane[12]. However, evidence regarding universal recycling versus degradation of internalized LTRs is conflicting. Indeed, recycling efficiency varies significantly among different LTRs, with shuttling receptors like TfR and sortilin exhibiting higher recycling rates compared to IGF2R and ASGPR[12, 42, 53]. Further mechanistic studies are essential to clarify the precise fate of the internalized LTRs, even though most studies assume complete recycling without assessing the levels of LTRs post-LYTAC treatment[19, 54]. In contrast to these conventional approaches, our study demonstrated that NCL-mediated internalization of SAPT8-based LYTACs resulted in the simultaneous degradation of NCL and POIs (c-Met and VEGFR-2). This likely occurs because NCL is not an intrinsic LTR and thus lacks a recycling mechanism. Although this raised initial concerns about potential depletion of surface NCL and consequent reduction in LYTAC efficacy, we observed a remarkable compensatory phenomenon: despite decreased total cellular NCL levels, membrane NCL expression was dynamically maintained throughout SAPT8-based LYTAC treatment. This compensation is possible because NCL constantly shuttles between the nucleus, cytoplasm, and cell surface of tumor cells and RA-FLSs[23, 55], thus maintaining sufficient surface expression for LYTAC engagement. We therefore propose the following self-renewing cycle: (1) SAPT8-based LYTACs facilitate the elimination of surface NCL along with the POI, (2) Intracellular NCL pools rapidly replenish surface expression, (3) Extracellular LYTACs bind the newly translocated NCL, initiating subsequent rounds of POI and NCL degradation. This cycle persists until the LYTACs are fully utilized, sustaining therapeutic efficacy.

Up to now, our identified SAPT8 is not the only aptamer to be explored for targeting NCL. AS1411, another NCL-binding aptamer, has undergone extensive investigation for decades and advanced to phase I/II clinical trials as an antitumor agent, although it did not achieve clinical success[56–58]. This raises the question of whether AS1411 could similarly serve as a lysosome-directed ligand for LYTACs. However, mechanistic studies reveal critical differences: AS1411 primarily inhibits NCL phosphorylation and fails to undergo lysosomal trafficking for inducing NCL degradation after cellular internalization[58, 59]. This is attributed to its predominant uptake via macropinocytosis, which allows it to escape the endosome and access the cytoplasm due to the leaky nature of macropinosomes[60]. In contrast, SAPT8 employs both clathrin- and caveolae-mediated endocytosis, which are known to direct cargo to lysosomes[23]. This fundamental distinction in cellular processing routes explains the suitability of SAPT8 for LYTAC design. Intriguingly, we have suggested an alternative application of AS1411 in TPD strategies. Given lysosomal escape, cytoplasmic access, as well as the established intracellular interaction between NCL and the E3 ligase murine double minute 2 (MDM2)[61], we propose that AS1411 could serve as an MDM2 recruiter via employing NCL as a molecular bridge[62, 63]. This insight enabled our development of AS1411-based, NCL-bridged PROTACs capable of inducing proteasomal degradation of intracellular POIs[32, 64, 65]. Critically, the molecular basis for the distinct endocytic pathways utilized by AS1411 and SAPT8 remains unclear and warrants further investigation. Understanding these differences could optimize NCL aptamer-based strategies for degrading both intracellular (via PROTACs) and extracellular (via LYTACs) POIs.

In this work, we developed two distinct LYTAC architectures for c-Met degradation: dual-aptamer SAPT8-SL1 chimeras and aptamer-small-molecule SAPT8-Tep hybrids. Comparative analysis revealed that the SAPT8-Tep LYTACs demonstrated superior c-Met degradation efficacy compared to SAPT8-SL1 chimeras. Several factors may contribute to this performance difference. First, SAPT8-SL1 LYTACs were entirely oligonucleotide-based macromolecules, consisting of two aptamers connected by a nucleic acid linker, resulting in significantly larger structures. In contrast, SAPT8-Tep hybrids are smaller constructs, formed by covalently linking the SAPT8 aptamer to a small molecule Tep. The larger size of SAPT8-SL1 LYTACs may impede cellular entry, thereby reducing effectiveness. Second, the SAPT8-SL1 design relies on the non-covalent coupling of the two aptamers via complementary A-T base pairing. This arrangement likely offers lower stability compared to the covalent bonds utilized in SAPT8-Tep hybrids. Third, Linker length and composition are crucial determinants of LYTAC activity[13]. The dual-aptamer LYTACs employed a single nucleic acid linker, whereas the SAPT8-Tep hybrids incorporated three distinct linkers: two alkyl linkers of different lengths and a PEG linker. This enhanced linker flexibility in the hybrid design may contribute to its improved performance[66]. Future research should explore a broader range of linker configurations to optimize the therapeutic efficacy of both LYTACs.

Collectively, we developed a novel SAPT8-based LYTAC platform that achieves dual functionality: cell-type-specific targeting (tumor cells or FLSs) via NCL recognition and lysosomal degradation of extracellular POIs. This proof-of-concept study expands the therapeutic landscape of TPD for extracellular POIs. By leveraging SAPT8-mediated NCL engagement, we establish a compelling alternative to conventional LYTAC strategies reliant on intrinsic LTRs.

## MATERIALS AND METHODS

### Cell culture

The following cell lines were obtained from the American Type Culture Collection (ATCC, USA): human osteosarcoma U-2 OS (HTB-96), human cervical carcinoma HeLa (CRM-CCL-2), human non-small cell lung cancer A549 (CCL-185), human colorectal HCT116 (CCL-247), and human normal epithelial MCF 10A (CRL-10317). RA-FLSs were isolated from synovial tissues of RA patients[67]. Cells were cultured in Dulbecco’s Modified Eagle’s Medium (DMEM), McCoy’s 5A Medium, or RPMI-1640 medium (Corning), supplemented with 10% fetal bovine serum (FBS) and 1% penicillin-streptomycin. All cell lines tested negative for mycoplasma contamination and were maintained at 37°C in a humidified atmosphere with 5% CO_2_ and 95% humidity.

### Construction of LYTACs

The DNA oligonucleotides, including NC, SAPT8, SL1, and Apt02, were synthesized by Sangon Biotech Co., Ltd (Shanghai, China). The SAPT8-SL1 LYTAC degraders were assembled as follows: the 5’-terminus of SAPT8 was ligated to the sense strand of double-stranded DNA linker (6 A), while the 3’- or 5’-terminus of SL1 was conjugated to the corresponding antisense strands of the DNA linker (6 T). The resulting sequences were heated at 95°C for 5 min, followed by gradual cooling at 37°C for 30 min, and then mixed at a molar ratio of 1:1 at 4°C overnight. The sequences used were as follows: SAPT8: 5’-CCGGCTGCTGCGGC ATCGGGCTAGATATCGAT-3’[23]. NC: 5’-GGAATTCCCGGTGCGCCGATCGCCGGATATAACTT-3’[23]. SL1: 5’-ATCAGGCTGGATGGTAGCTCGGTCGGGGTGGGTGGGTTGGCAAGTCTGAT-3’[32]. Double-stranded DNA linker (6 A-T base pairs): 5’-AAAAAA-3’/5’-TTTTTT-3’[68]. For assembly of SAPT8-Apt02 LYTAC degraders, SAPT8 was linked to the 3’ or 5’ end of Apt02 via a single-stranded DNA linker consisting of six T bases. Both SAPT8-Apt02-1 and SAPT8-Apt02-2 were synthesized by Sangon Biotech Co., Ltd. (Shanghai, China). Apt02: 5′-GCTGATAGGATGGGTTGTAGGTCTAGGGGGGGGCC-3′[69].

### Binding assay by flow cytometry

U-2 OS cells (6×10^4^ per well), HeLa cells (5×10^4^ per well), HCT116 cells (5×10^4^ per well), or MCF 10A (6×10^4^ per well) were seeded in 6-well plates and cultured for 24 h. After the treatments in each study, these cells were digested using 0.25% trypsin-EDTA (Thermo Fisher, USA) and washed with PBS. Fluorescence signals were detected by a flow cytometer (BD, FACSCanto SORP, USA), and data were analyzed using FlowJo software. To measure the equilibrium dissociation constant (*K_d_*), U-2 OS cells were treated with increasing concentrations of Cy3-labeled SAPT8 aptamer. The *K_d_* value for the interaction between the SAPT8 aptamer and U-2 OS cells was determined by fitting the fluorescence intensity data with Prism GraphPad 9 software using the equation *Y* = B_max_ *X*/(K_d_+*X*), where *X* represents the aptamer concentration, *Y* is MFI at concentration *X*, and B_max_ is maximum MFI[70].

### Pull-down assays

After different treatments, cells were lysed with IP lysis buffer (pH 7.4, 0.025 M Tris, 0.15 M NaCl, 0.001 M EDTA, 1% NP-40, and 5% Glycerol; Thermo, USA) supplemented with a proteinase inhibitor cocktail (Sigma, USA). Supernatants were incubated with Streptavidin Sepharose High Performance Beads (GE Healthcare, UK) at 4 °C overnight. After washing the beads five times, the sepharose beads were resuspended in 1×SDS-PAGE protein loading buffer and heated at 100 °C for 10 min to elute bound proteins. Protein samples of the pull-down assay and input were analyzed by SDS–PAGE and western blotting[71].

### Sub-cellular distribution of SAPT8

U-2 OS cells (2×10^4^ per well) were seeded in confocal plates and treated with 200 nM Cy3-labeled NC or SAPT8 for 7 h. Cells were then stained with Lyso-Tracker Red dyes for 30 min, washed five times with PBS, fixed with 4% paraformaldehyde for 20 min, and counterstained with DAPI (Beyotime, China) for 10 min to label nuclei. Images were acquired using a confocal fluorescence microscope (Zeiss, LSM980, DE)[72].

### Co-IP assay

Co-IP assay was performed as previously described[73]. Briefly, cells subjected to different treatments were lysed with IP lysis buffer (pH 7.4, 0.025 M Tris, 0.15 M NaCl, 0.001 M EDTA, 1% NP-40, and 5% Glycerol; Thermo, USA) supplemented with a proteinase inhibitor cocktail (NCE, China). Cell lysates were incubated with the corresponding primary antibody (anti-NCL, 14574, CST; anti-c-Met, 8198, CST; anti-VEGFR-2, ab134191) overnight at 4 °C, followed by coupling to Protein A/G Magnetic Beads (Thermo Fisher Scientific, USA) at room temperature for 1.5 h. After washing the beads five times using a DynaMag™-2 Magnet (Thermo Fisher Scientific, USA), the immunoprecipitated complexes were resuspended in 1×SDS-PAGE protein loading buffer, heated at 100 °C for 10 min to elute bound proteins, and analyzed by SDS-PAGE and Western blotting.

### Western blotting

Western blotting was performed as previously described[74]. Briefly, following treatments, cells were lysed with lysis buffer (50 mM Tris-HCl (pH 7.4), 150 mM NaCl, 1% Triton X-100, 2 mM EDTA, and 10% glycerol) containing a protease inhibitor cocktail (Sigma, USA). Total protein concentrations were quantified using a bicinchoninic acid (BCA) assay (Thermo Fisher Scientific, USA). Proteins from each sample were separated by SDS-PAGE and transferred to polyvinylidene fluoride (PVDF) membrane (Millipore, MA, USA) via a transfer apparatus (Bio-Rad, Trans-Blot Turbo™, USA). After blocking, membranes were incubated with primary antibodies (anti-NCL, 14574, CST; anti-c-Met, 8198, CST; anti-p53, 2524, CST; anti-Bcl-2, 12789-1-AP, Proteintech; anti-GAPDH, AC002, ABclonal) overnight at 4°C. Following three washes with Tris-buffered Saline containing 0.1% Tween20 (TBST, T1081, Solarbio, CHN), membranes were incubated with HRP-conjugated secondary antibodies for 1 h at room temperature. Immunodetection was performed using an enhanced chemiluminescence ECL kit (ABclonal, CHN) and visualized with a chemiluminescence imaging system (Tanon, Multi5200, CHN).

### Gene silencing by siRNA

Cells were seeded in 6-well plates and allowed to adhere for 24 h until reaching 70-80% confluency. Following the manufacturer’s protocol, cells were transfected with 20 nM siRNA using Lipofectamine RANiMAX Reagent (Thermo Fisher, USA). siRNA sequences were as follows: siNCL (sense strand, 5’-GGAUGACGACGACG ACGAAGATT-3’; antisense strand, 5’-UCUUCGUCGUCGUCGUCAUCCTT-3’); and siCtrl (sense strand, 5’-UUCUCCGAACGUG UCACGUTT-3’; antisense strand, 5’-ACGUGACACGU UCGGAGAATT-3’)[75].

### Lentivirus production and transduction

Lentivirus production and transduction were performed as previously described[76]. Briefly, a mixture of lentiviral plasmids, psPAX2, and pMD2.G at a weight ratio of 5: 2: 3 was co-transfected into 293T cells using Effectene transfection reagent (Qiagen, Germany). The culture medium was replenished at 12 h post-transfection. Viral supernatant was collected at 48 h and 72 h. For transduction, U-2 OS cells (8×10^4^ per well) were seeded in 6-well plates and allowed to adhere for 24 h. Cells were then incubated with lentivirus supernatant containing 8 μg/mL polybrene (Sigma-Aldrich, USA). The culture medium was changed at 36 h post-transfection, and antibiotic selection was initiated at 72 h post-transfection using 2 μg/mL puromycin (Beyotime, CHN). The targeted gene expression in U-2 OS cells was measured 3 days after initiating antibiotic selection. The information on plasmids used in a prior study[23].

### Colony formation assay

Cells were seeded in 6-well plates and allowed to adhere for 24 h. After different treatments in each study, the cells were fixed in a 4% paraformaldehyde solution (Dingguo Biotechnology, China) for 20 min and stained with crystal violet (Beyotime, CHN) for 30 min at room temperature. Colony images were captured using a digital camera[77].

### EdU assay

Cells and tumor sections were processed according to the BeyoClick™ EdU Cell Proliferation Kit protocol (Beyotime, CHN). Briefly, after different treatments for 5 days, cells were incubated with 10 µM EdU reagent (500 μL/well) for 2 h, washed three times with PBS, fixed with 4% paraformaldehyde for 10 min, permeabilized with 0.3% Triton X-100 for another 10 min, and incubated with click-reaction reagent in the dark for 30 min. For tumor sections, deparaffinized sections were incubated with 10 µM EdU reagent (500 μL/section) for 2 h, followed by click-reaction reagent in the dark for 30 min. All samples were counterstained with DAPI and imaged using a confocal microscope (Zeiss LSM980, Germany)[78].

### TUNEL assay

Cells and tumor sections were processed using the One Step TUNEL Apoptosis Assay Kit (Beyotime, CHN). Briefly, after fixation with 4% paraformaldehyde for 10 min, samples were incubated with Cy3-labeled dUTP and terminal deoxynucleotidyl transferase enzyme in the dark for 1 h. After washing three times, DAPI was used to counterstain the nucleus. The result of staining was determined with a confocal fluorescence microscope (Zeiss, LSM980, DE)[79].

### Transwell migration assay

To examine cell migration, RA-FLSs (7.5 × 10³ cells per well) were seeded directly into the upper chamber of 24-well Transwell inserts (pore size 8.0 μm; Corning, NY) in 200 μL of serum-free medium. The lower chamber received 600 μL of medium containing 20% FBS. After different treatments, non-migrated cells on the upper surface of the membrane were removed using cotton swabs. Cells that had migrated onto the lower surface of the membrane were fixed with 4% formalin for 20 min and stained with crystal violet for 30 min at room temperature[80]. Subsequently, the inserts were washed with PBS and air-dried. Migrated cells were photographed using an inverted microscope (NanoZoomer S60; Hamamatsu, Japan) and quantified by counting with ImageJ software[80].

### Animals

Female and male BALB/c nude mice and male SCID mice (6-week-old) were purchased from the GemPharmatech Co., Ltd (Guangdong, China) and housed under standard conditions (22 °C, 12 h light/dark cycle) within the Laboratory Animal House at the Southern University of Science and Technology. All *in vivo* studies have gained ethics approval from the Institutional Animal Care and Use Committee (IACUC) of the Southern University of Science and Technology.

### *In vivo* fluorescence imaging

Cy3-labeled NC or SAPT8, and Cy5-labeled SL1, SAPT8-SL1-1, or SAPT8-SL1-2 were intravenously injected into tumor-bearing BALB/c nude mice. Whole-body fluorescence was captured using an IVIS® Spectrum system (PerkinElmer, USA). Post-imaging, tumors and major organs (heart, lung, liver, spleen, and kidney) were excised for ex vivo imaging. Fluorescence intensity was quantified with Living Image v4.4 software (Caliper Life Sciences, USA)[81].

### Orthotopic xenograft model

SCID mice received intra-bone marrow injections of 6×10^5^ U-2 OS cells (20 μL volume) using a 27 G needle. Mice were treated via tail vein every 2 days, starting 24 days post-inoculation. Tumor volume and body weight were monitored until sacrifice on day 34[82].

### Subcutaneous xenograft model

BALB/c nude mice were subcutaneously implanted with 1×10^7^ U-2 OS cells or 5×10^6^ HeLa cells suspended in Matrigel (Solarbio, CHN). Treatments began after 4 days post-implantation and continued every 2 days. Tumors were excised and weighed on day 16[83].

### μCT analysis

These joints were fixed in 10% formalin (Solarbio, CHN), decalcified in 70% ethanol, and scanned using a Skyscan scanner 1276 high-resolution µCT scanner (Bruker, Germany) with a voxel size of 10 μm, at 60 kVp/100 µAmp. 3D reconstructions were generated with NRecon and CTvox software (Bruker, Germany)[84].

### Histological analysis

Tissue sections underwent deparaffinization in xylene and were progressively rehydrated through a graded series of ethanol to water. A series of sections (5 µm thick) was then stained with H&E (Solarbio, CHN) using standard protocols according to the manufacturer’s instructions. The specimens were independently viewed and analyzed by two pathologists using an Aperio GT 450 scanner (Leica, DE)[81, 84, 85].

### Serum biochemical assays

Blood samples were collected from mice after treatment. These serum samples were collected via centrifugation of blood samples in 12000×g at 4 °C for 1 h. Serum biochemical parameters, including uric acid (UA), creatinine (CR), alanine transaminase (ALT), alkaline phosphatase (ALP), blood urea nitrogen (BUN), and total protein (TP), were analyzed using a MS-480 Automatic Biochemistry Analyzer (Medicalsystem Biotechnology, CHN)[86].

### Immunofluorescence staining

Tissue sections were subjected to deparaffinization using xylene, rehydration, and antigen retrieval. These sections were then permeabilized with 0.2% Triton X-100 for 10 min, blocked with 5% BSA in PBS for 1 h, and incubated with primary antibodies anti-NCL (14574, CST), anti-c-Met (8198, CST), anti-p53 (2524, CST), anti-Bcl-2 (12789-1-AP, Proteintech), and Ki-67 (9129, CST) at 4°C overnight. After washing three times using PBS, these sections were incubated with anti-mouse Alexa Fluor 488 (ab150113, Abcam) or anti-rabbit Alexa Fluor 488 (ab150077, Abcam) for 1 h at room temperature. Finally, these sections were counterstained with DAPI, and images were acquired using a confocal fluorescence microscope (Zeiss, LSM980, DE)[84].

### Statistical analysis

All numerical data are expressed as the mean ± SD. Statistical differences among multiple independent groups were analyzed by the one-way ANOVA with a post-hoc test. Statistical differences between two independent groups were determined by the two-tailed Student’s t-test. Statistical differences between the groups that were defined by two categorical factors were analyzed by the two-way ANOVA with a post hoc test. All statistical analyses were performed with GraphPad Prism 9 software. *p* < 0.05 was considered statistically significant. We chose representative images based on the average level of the data for each group. For the *in vivo* experiments, the sample size was pre-determined by a power calculation. The mice were grouped randomly and blindly to researchers. The mice in poor body condition before the experiments were excluded.

## Author Contributions

Fang Qiu: Methodology, Investigation, Formal analysis, Data curation, Visualization, Writing - original draft, Writing - review& editing. Ziting Feng: Conceptualization, Methodology, Writing-review & editing. Hongzhen Chen: Methodology, Data curation. Xiaoxuan Zhang: Methodology, Investigation. Jianmin Guo: Methodology, Investigation. Duoli Xie: Methodology, Investigation. Siran Yue: Methodology, Investigation. Chunhao Cao: Methodology, Investigation. Aiping Lu: Supervision, Funding acquisition. Chao Liang: Conceptualization, Methodology, Validation, Writing - review &editing, Supervision, Project administration, Funding acquisition.

## Acknowledgments

This work is supported by the National Key R&D Program of China (2024YFC3506200 and 2024YFC3506205 to C.L.), the National Natural Science Foundation Council of China (82472394 and 82172386 to C.L.), the Guangdong Basic and Applied Basic Research Foundation (2022A1515012164 to C.L.), the Shenzhen Science and Technology Program (JCYJ20210324104201005 and SGDX20240115112400001 to C.L.), the Hong Kong General Research Fund (12102722 and 12106424 to A.L.), the Hong Kong RGC Theme-based Research Scheme (T12-201/20-R to A.L.) and the Shenzhen LingGene Biotech Co., Ltd.

---

**Fig. S1.**
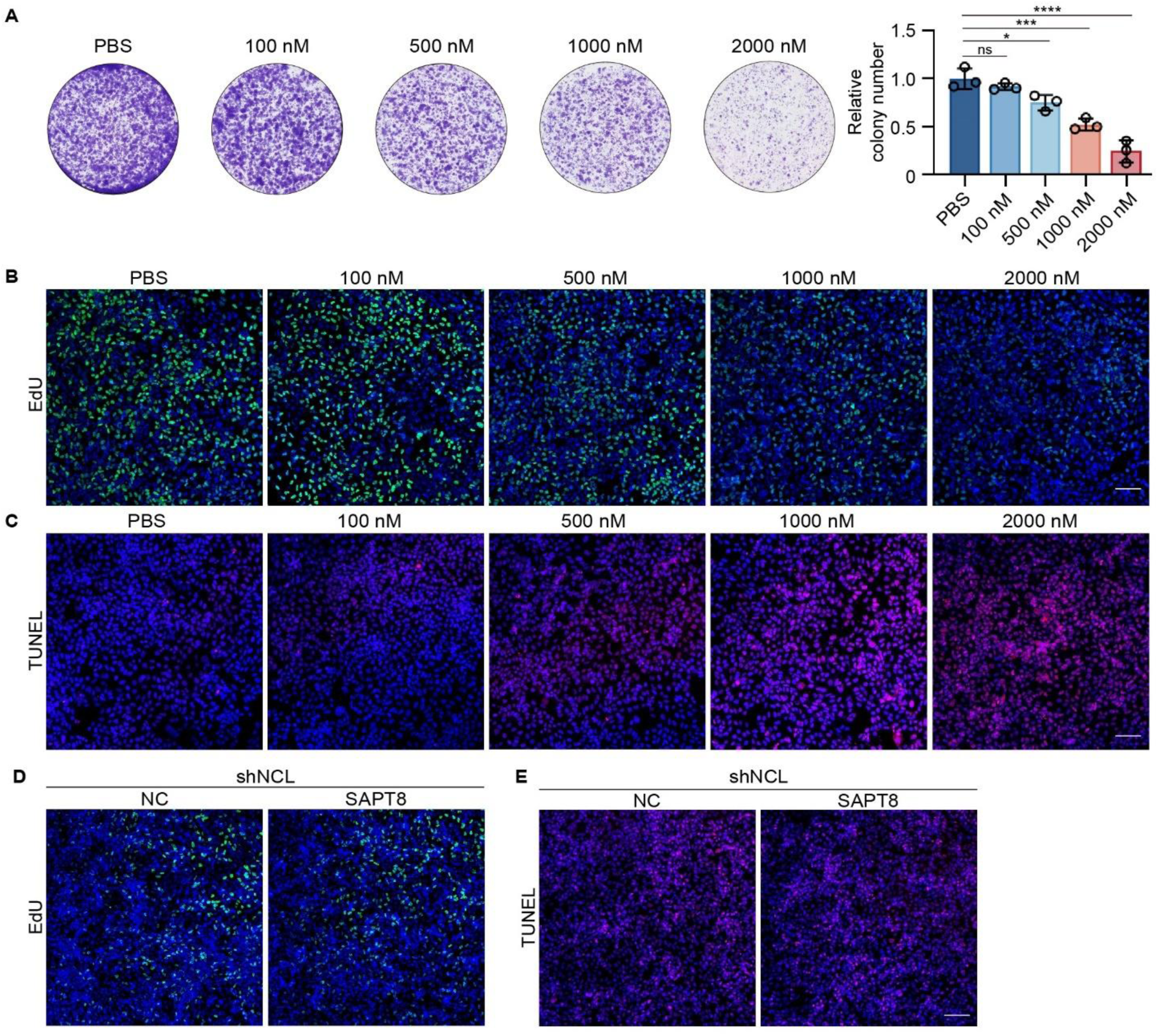
Antitumor effects of SAPT8 on U-2 OS cells *in vitro*. (**A**) Colony formation of U-2 OS cells treated daily with SAPT8 at different concentrations (0, 100, 500, 1000, and 2000 nM) for 10 days. (**B**) EdU assay for proliferation of U-2 OS cells treated daily with SAPT8 at different concentrations (0, 100, 500, 1000, and 2000 nM) for 6 days. (**C**) TUNEL assay for apoptosis of U-2 OS cells treated daily with SAPT8 at different concentrations (0, 100, 500, 1000, and 2000 nM) for 6 days. (**D**) Proliferation of NCL-silenced U-2 OS cells after incubation with 1000 nM NC and SAPT8 for 5 days, as measured by EdU staining. (**E**) Apoptosis of NCL-silenced U-2 OS cells after incubation with 1000 nM NC and SAPT8 for 5 days, as measured by TUNEL staining. Nuclei were stained with DAPI. Scale bars: 100 µm. Each experiment was repeated three times. Data were represented as mean ± SD, and statistical significance was calculated by the one-way ANOVA with a post hoc test (A). ns, no significance, *\*p* < 0.05, *\*\*\*p* < 0.001, *\*\*\*\*p* < 0.0001.

**Fig. S2.**
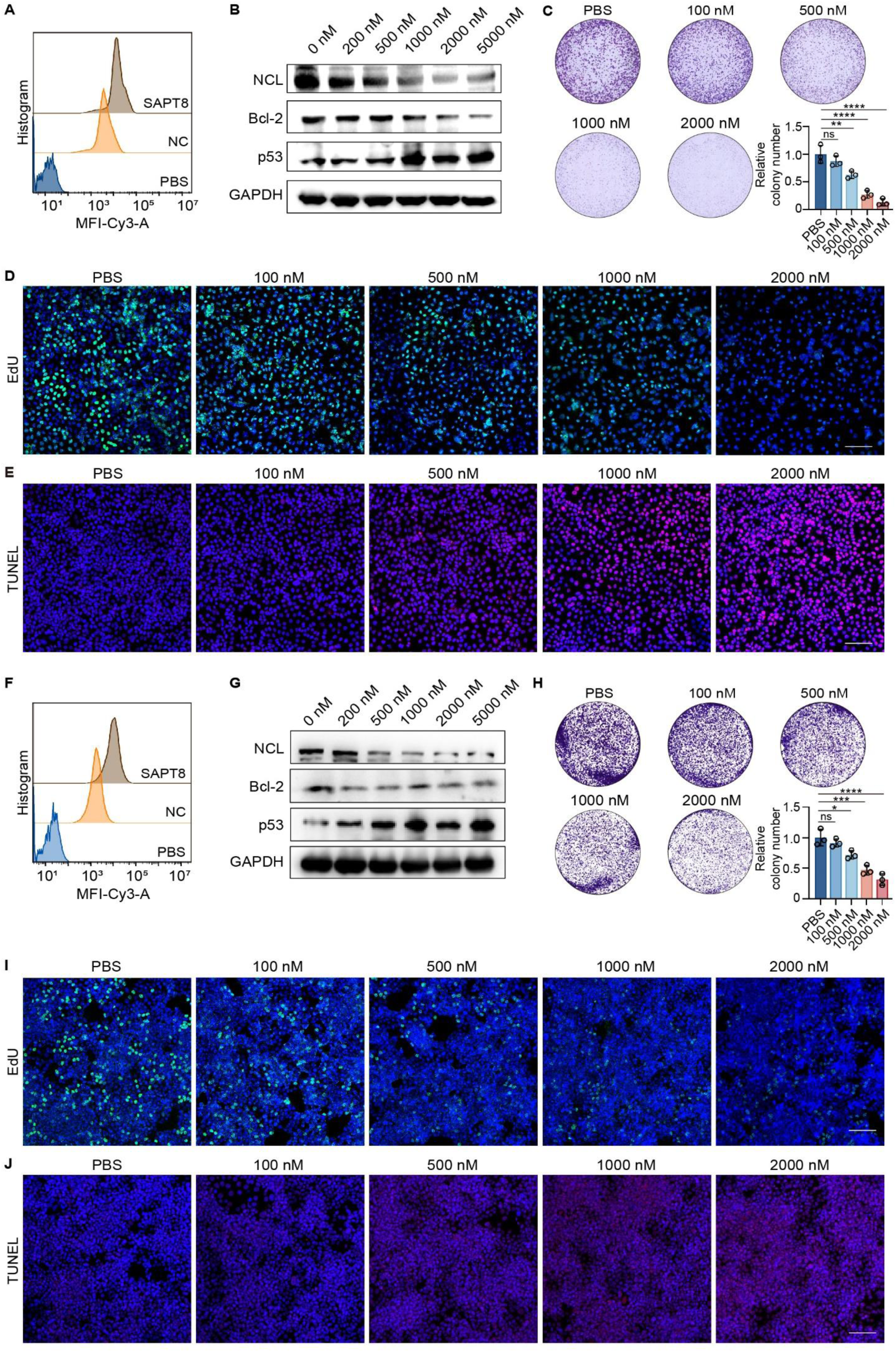
Tumor targeting and antitumor effects of SAPT8 on HeLa and HCT116 cells *in vitro*. (**A**) Binding assay of 200 nM Cy3-labeled NC or SAPT8 with HeLa cells after incubation for 3 h. (**B**) Protein expression of NCL, Bcl-2, and p53 in HeLa cells treated with SAPT8 at different concentrations (0, 200, 500, 1000, 2000, and 5000 nM) for 24 h. (**C**) Colony formation of HeLa cells after daily treatment with SAPT8 at different concentrations (0, 100, 500, 1000, and 2000 nM) for 8 days. (**D**) EdU assay for proliferation of HeLa cells after daily treatment with SAPT8 at different concentrations (0, 100, 500, 1000, and 2000 nM) for 5 days. Nuclei were stained with DAPI. Scale bars: 100 μm. (**E**) TUNEL assay for apoptosis of HeLa cells after daily treatment with SAPT8 at different concentrations (0, 100, 500, 1000, and 2000 nM) for 5 days. Scale bars: 100 µm. (**F**) Binding assay of 200 nM Cy3-labeled NC or SAPT8 with HCT116 cells after incubation for 3 h. (**G**) Protein expression of NCL, Bcl-2, and p53 in HCT116 cells after treatment with SAPT8 at different concentrations (0, 200, 500, 1000, 2000, and 5000 nM) for 24 h. (**H**) Colony formation of HCT116 cells after daily treatment with SAPT8 at different concentrations (0, 100, 500, 1000, and 2000 nM) for 6 days. (**I**) EdU assay for proliferation of HCT116 cells after daily treatment with SAPT8 at different concentrations (0, 100, 500, 1000, and 2000 nM) for 5 days. Nuclei were stained with DAPI. Scale bars: 100 μm. (**J**) TUNEL assay for apoptosis of HCT116 cells after daily treatment with SAPT8 at different concentrations (0, 100, 500, 1000, and 2000 nM) for 5 days. Nuclei were stained with DAPI. Scale bars: 100 µm. Each experiment was repeated three times. Data were represented as mean ± SD, and statistical significance was calculated by the one-way ANOVA with a post hoc test (C and H). ns, no significance, *\*p* < 0.05, *\*\*p* < 0.01, *\*\*\*p* < 0.001, *\*\*\*\*p* < 0.0001.

**Fig. S3.**
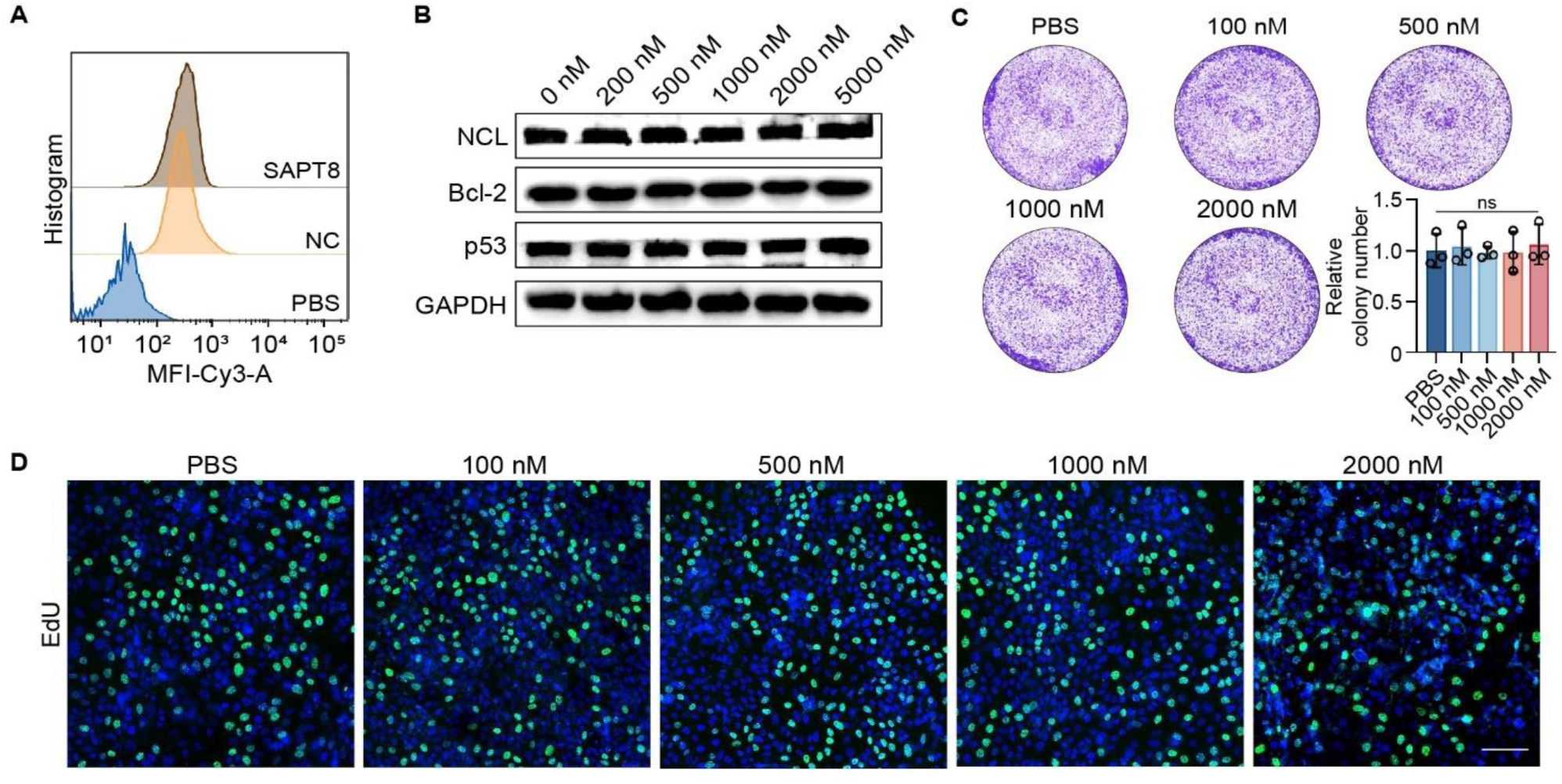
Effects of SAPT8 on normal MCF 10A cells *in vitro*. (**A**) Binding assay of 200 nM Cy3-labeled NC or SAPT8 with MCF 10A cells after incubation for 3 h. (**B**) Protein expression of NCL, Bcl-2, and p53 in MCF 10A cells after treatment with SAPT8 at different concentrations (0, 200, 500, 1000, 2000, and 5000 nM) for 24 h. (**C**) Colony formation of MCF 10A cells after daily treatment with SAPT8 at different concentrations (0, 100, 500, 1000, and 2000 nM) for 7 days. (**D**) EdU assay for proliferation of MCF 10A cells after daily treatment with SAPT8 at different concentrations (0, 100, 500, 1000, and 2000 nM) for 5 days. Nuclei were stained with DAPI. Scale bars: 100 μm. Each experiment was repeated three times. Data were represented as mean ± SD, and statistical significance was calculated by the one-way ANOVA with a post hoc test (C). ns, no significance.

**Fig. S4.**
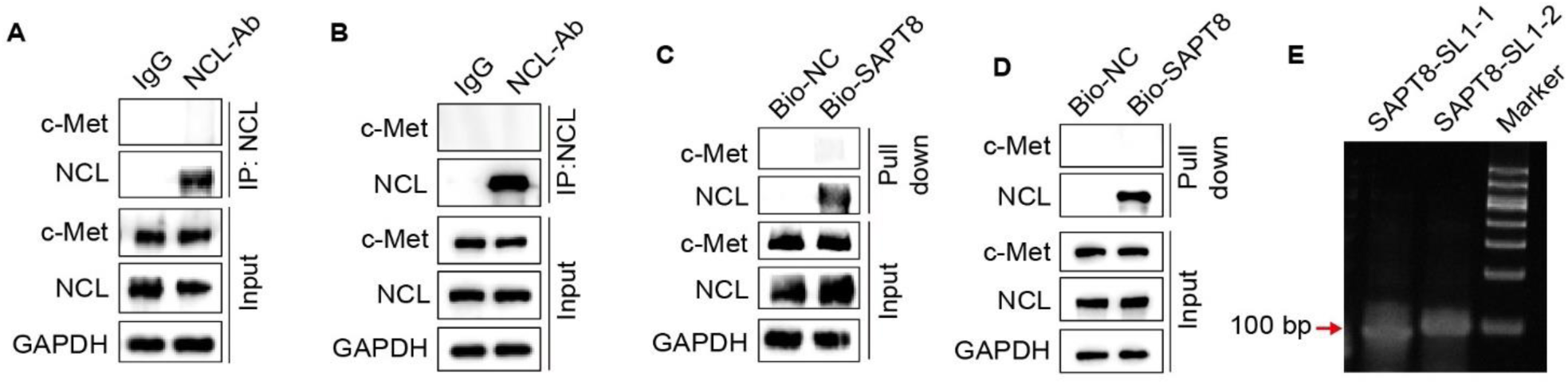
No interaction between NCL and c-Met *in vitro*. (**A**) Co-IP of c-Met and NCL in U-2 OS cells using an anti-NCL antibody. (**B**) Co-IP of c-Met and NCL in HeLa cells using an anti-NCL antibody. (**C**) Pull-down assay of c-Met and NCL in U-2 OS cells treated with 1 μM biotin (Bio)-labelled NC or SAPT8 for 8 h. (**D**) Pull-down assay of c-Met and NCL in HeLa cells treated with 1 μM Bio-labeled NC or SAPT8 for 8 h. (**E**) Nondenaturing polyacrylamide gel electrophoresis (PAGE) of SAPT8-SL1-1 and SAPT8-SL1-2. Each experiment was repeated three times.

**Fig. S5.**
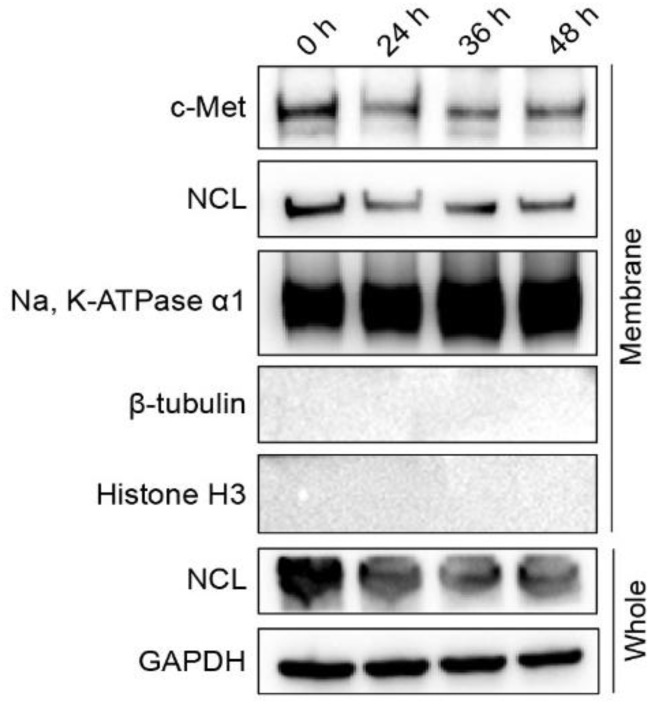
NCL levels across different subcellular compartments *in vitro*. Protein expression of NCL in different subcellular compartments of U-2 OS cells was detected after treatment with 800 nM SAPT8-SL1-2 for different time points (0, 24, 36, and 48 h). Each experiment was repeated three times.

**Fig. S6.**
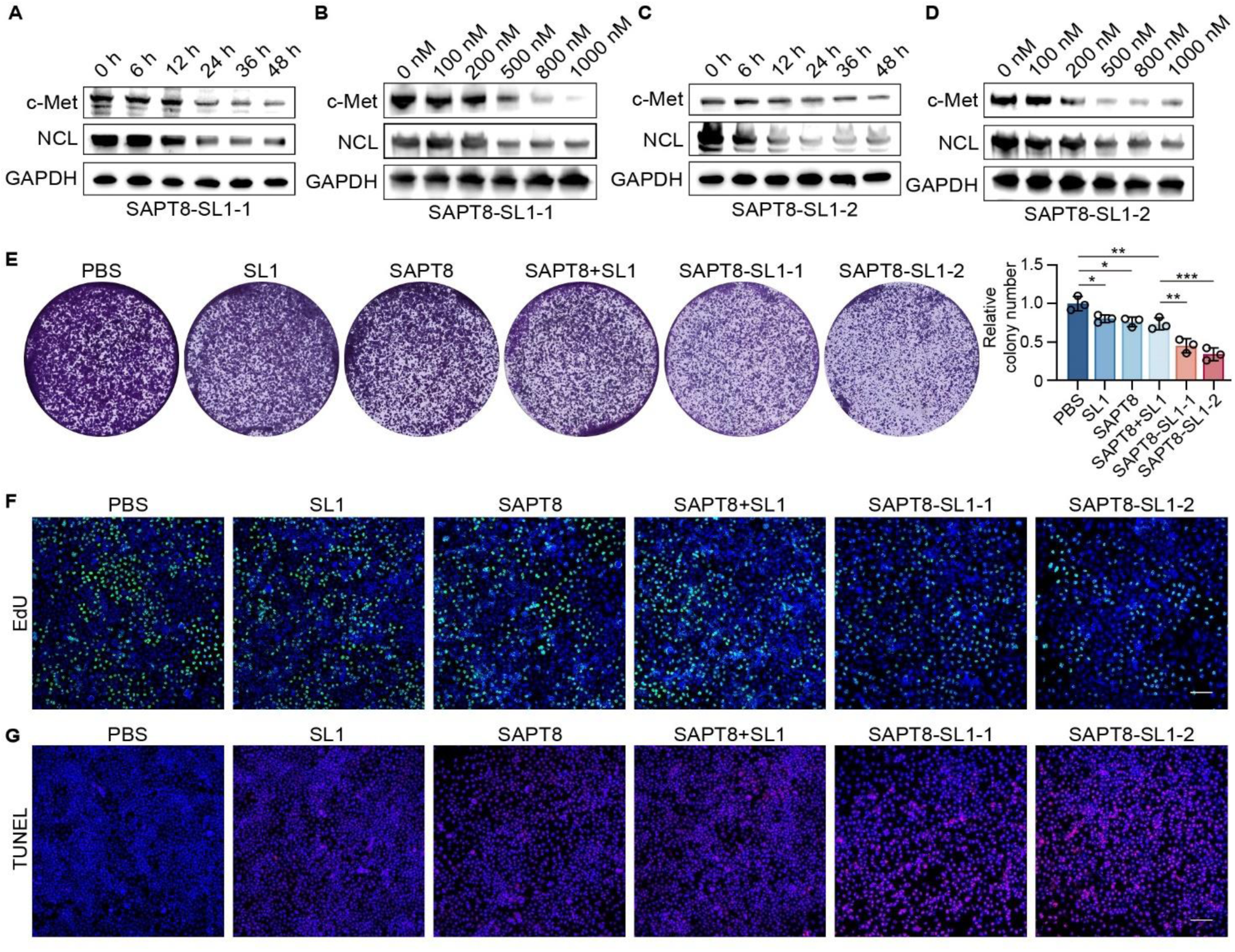
Degradation of c-Met and antitumor activity of SAPT8-SL1 LYTACs in HeLa cells *in vitro*. (**A**) Protein expression of c-Met and NCL in HeLa cells after treatment with 500 nM SAPT8-SL1-1 for different time points (0, 6, 12, 24, 36, and 48 h). (**B**) Protein expression of c-Met and NCL in HeLa cells treated with SAPT8-SL1-1 at different concentrations (0, 100, 200, 500, 800, and 1000 nM) for 12 h. (**C**) Protein expression of c-Met and NCL in HeLa cells treated with 500 nM SAPT8-SL1-2 for different time points (0, 6, 12, 24, 36, and 48 h). (**D**) Protein expression of c-Met and NCL in HeLa cells treated with SAPT8-SL1-2 at different concentrations (0, 100, 200, 500, 800, and 1000 nM) for 12 h. (**E**) Colony formation of HeLa cells after daily treatment with SL1, SAPT8, SAPT8+SL1, SAPT8-SL1-1, or SAPT8-SL1-2 at a concentration of 500 nM for 7 days. (**F**) EdU assay for proliferation of HeLa cells treated daily with SL1, SAPT8, SAPT8+SL1, SAPT8-SL1-1, or SAPT8-SL1-2 at a concentration of 500 nM for 5 days. Nuclei were stained with DAPI. Scale bars: 100 μm. (**G**) TUNEL assay for apoptosis of HeLa cells treated daily with SL1, SAPT8, SAPT8+SL1, SAPT8-SL1-1, or SAPT8-SL1-2 at a concentration of 500 nM for 5 days. Nuclei were stained with DAPI. Scale bars: 100 µm. Each experiment was repeated three times. Data were represented as mean ± SD, and statistical significance was calculated by the one-way ANOVA with a post hoc test (E). *\*p* < 0.05, *\*\*p* < 0.01, *\*\*\*p* < 0.001.

**Fig. S7.**
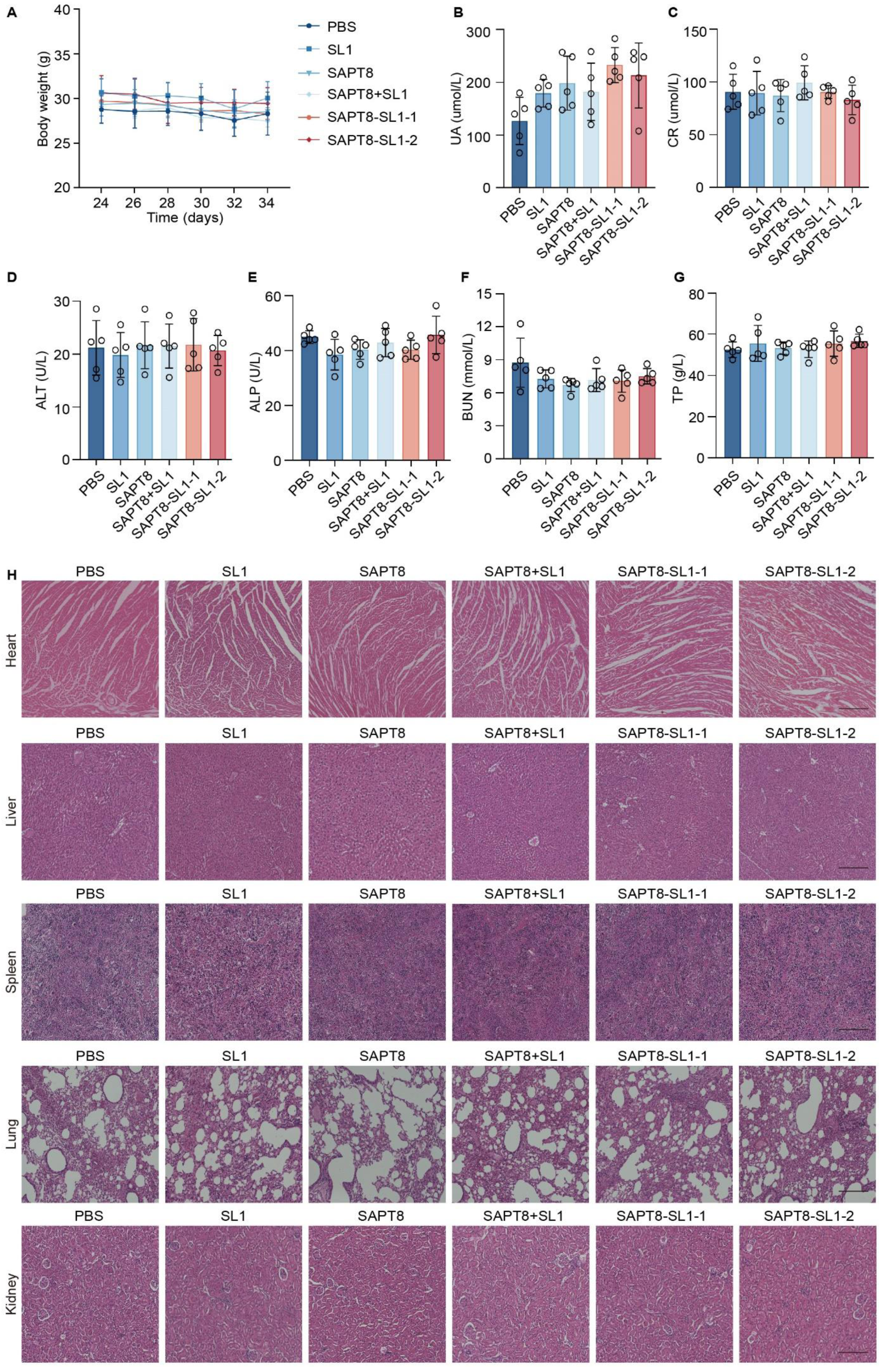
Safety of SAPT8-SL1 LYTACs in an orthotopic xenograft mouse model. (**A**) Changes in body weight of SCID mice bearing orthotopic xenografts in each treatment group. (**B**-**G**) Levels of serum uric acid (UA) (**B**), creatinine (CR) (**C**), alanine transaminase (ALT) (**D**), alkaline phosphatase (ALP) (**E**), blood urea nitrogen (BUN) (**F**), and total protein (TP) (**G**) in mice bearing orthotopic xenografts after different treatments, as determined by the automated hematology analyzer. (**H**) Histological assessments of major organs, including heart, liver, spleen, lung, and kidney, from mice bearing orthotopic xenografts in each treatment group using H&E staining. Scale bars: 100 μm. n = 5 for each group. Data were represented as mean ± SD, and statistical significance was calculated by the two-way ANOVA with a post hoc test (A) or one-way ANOVA with a post hoc test (B-G).

**Fig. S8.**
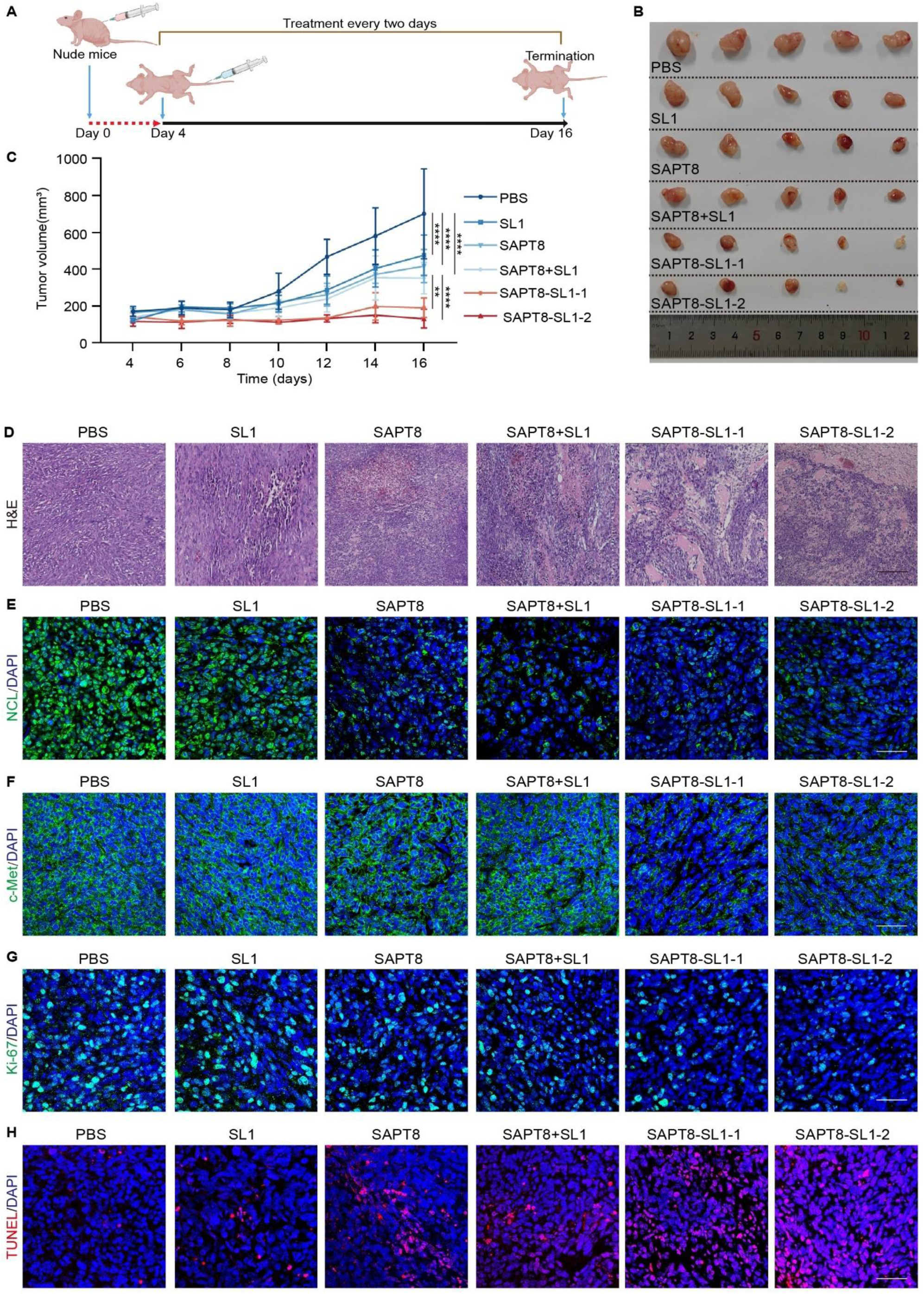
Antitumor efficacy of SAPT8-SL1 LYTACs in a subcutaneous xenograft mouse model. (**A**) Illustration of the experimental design for determining the antitumor efficacy of SAPT8-SL1 in a subcutaneous xenograft mouse model. Briefly, BALB/c nude mice were subcutaneously injected with 1×10^7^ U-2 OS cells. After 4 days, the mice bearing U-2 OS xenografts were intravenously treated with SL1, SAPT8, SAPT8+SL1, SAPT8-SL1-1, and SAPT8-SL1-2 for 12 days at a dose of 6 μmol/kg every two days. (**B**) Images of xenograft tumors from each treatment group. (**C**) Tumor volume in nude mice from each treatment group. (**D**) H&E staining of tumor sections from each treatment group. Scale bars: 100 μm. (**E**-**G**) Immunofluorescence staining of tumor sections for detecting the levels of NCL (**E**), c-Met (**F**), or Ki-67 (**G**) in each treatment group. (**H**) TUNEL assay for detecting apoptotic cells in tumor sections from each treatment group. Scale bars: 50 μm, n = 5 for each group. Data were represented as mean ± SD, and statistical significance was calculated by the two-way ANOVA (C) with a post hoc test. *\*\*p* < 0.01, *\*\*\*\*p* < 0.0001.

**Fig. S9.**
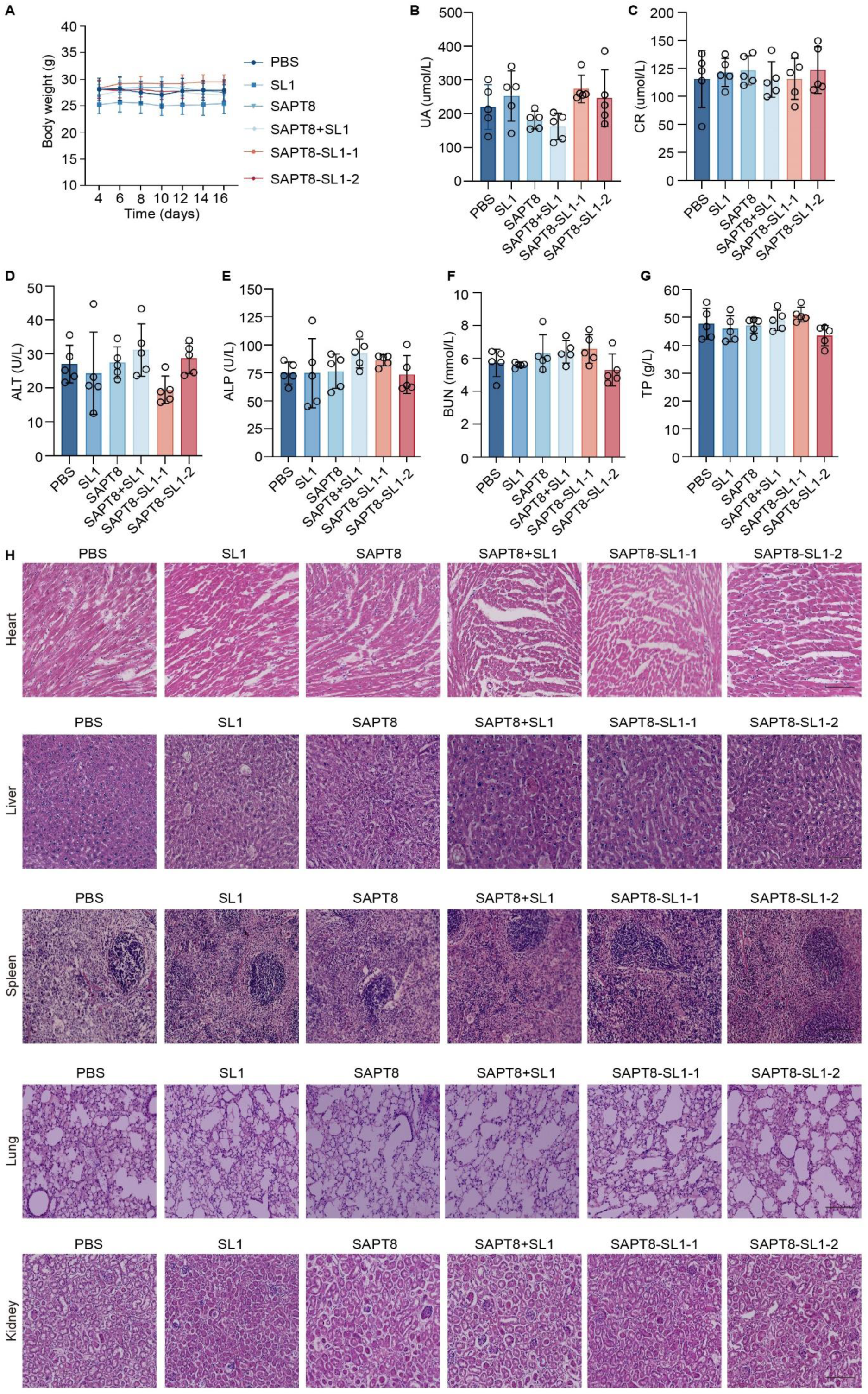
Safety of SAPT8-SL1 LYTACs in a subcutaneous xenograft mouse model. (**A**) Changes in body weight of BALB/c nude mice bearing xenograft tumors in each treatment group. (**B**-**G**) Levels of serum UA (**B**), CR (**C**), ALT (**D**), ALP (**E**), BUN (**F**), and TP (**G**) in nude mice bearing subcutaneous xenograft tumors after different treatments, as determined by the automated hematology analyzer. (**H**) Histological assessments of major organs, including heart, liver, spleen, lung, and kidney, from nude mice in each treatment group using H&E staining. Scale bars: 100 μm. n = 5 for each group. Data were represented as mean ± SD, and statistical significance was calculated by the two-way ANOVA with a post hoc test (A) or one-way ANOVA with a post hoc test (B-G).

**Fig. S10.**
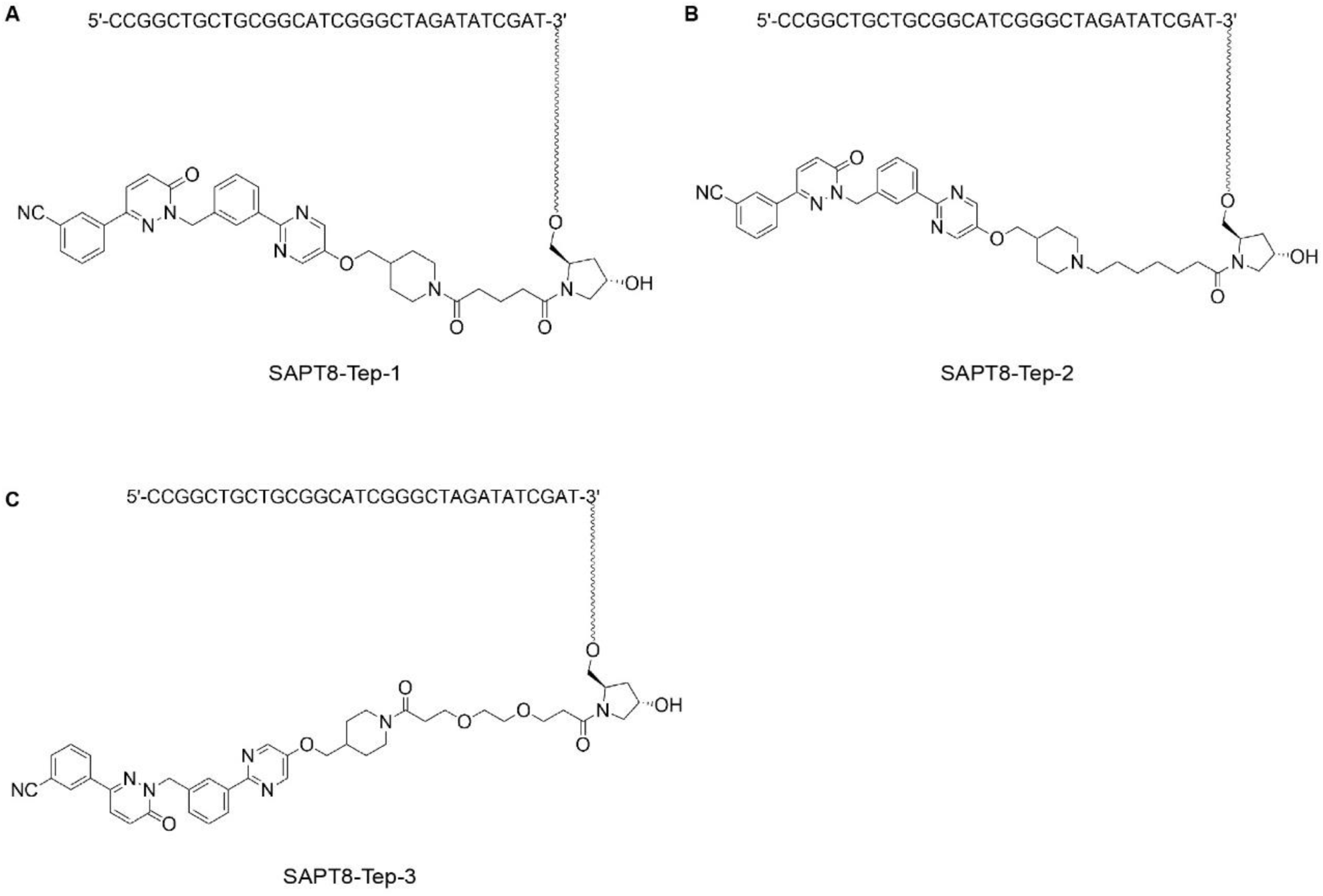
Construction of SAPT8-Tep LYTACs. Briefly, SAPT8 was conjugated with Tep via alkyl chains with different lengths or a PEG linker, resulting in the generation of SAPT8-Tep-1(**A**), SAPT8-Tep-2 (**B**), and SAPT8-Tep-3 (**C**).

**Fig. S11.**
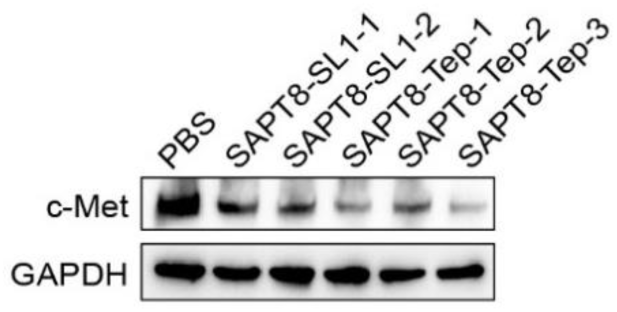
Comparison of SAPT8-SL1 LYTACs and SAPT8-Tep LYTACs *in vitro*. The level of c-Met in U-2 OS cells was determined after treatment with SAPT8-SL1-1, SAPT8-SL1-2, SAPT8-Tep-1, SAPT8-Tep-2, or SAPT8-Tep-3 at a concentration of 600 nM for 24 h.

**Fig. S12.**
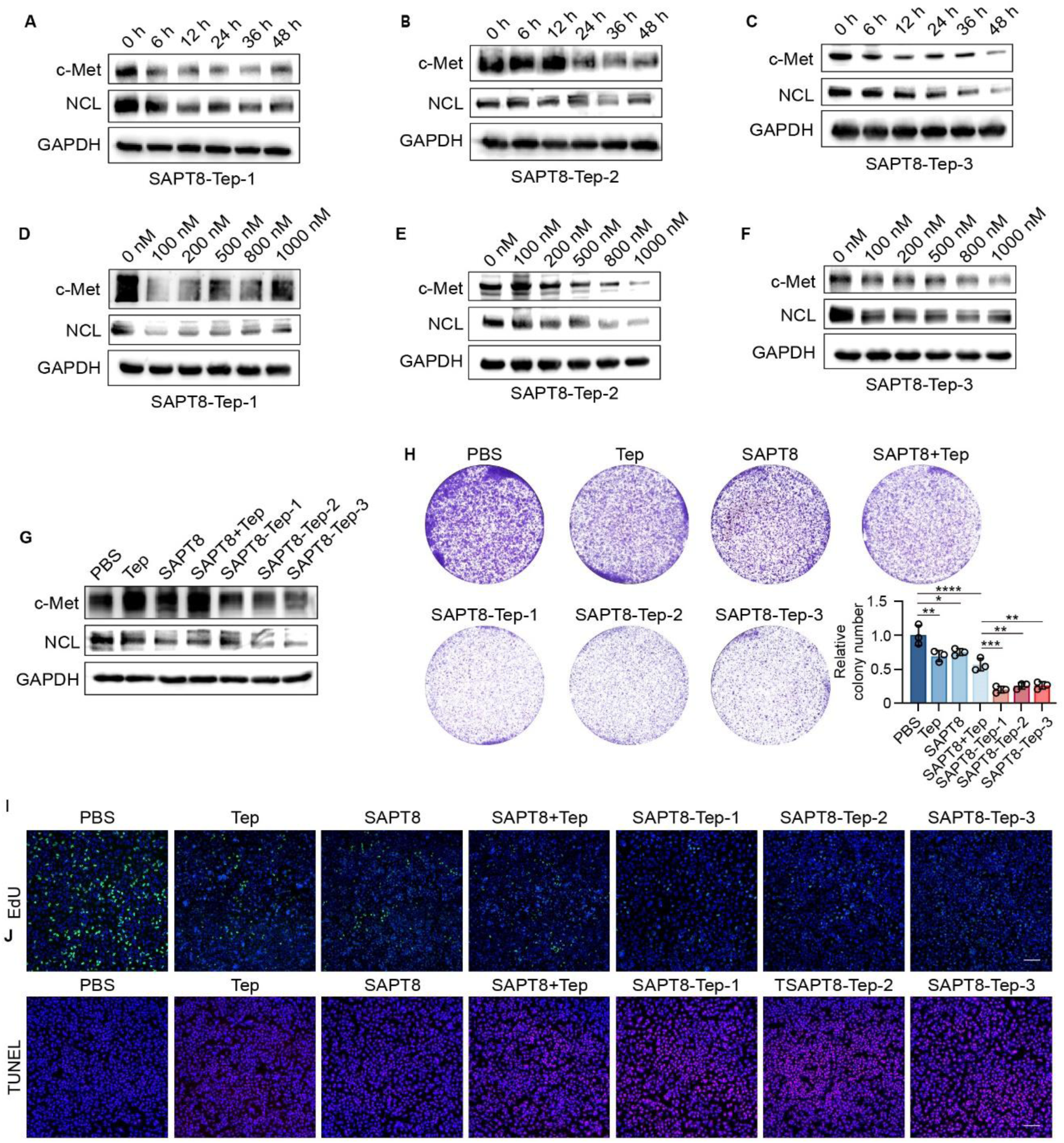
Degradation of c-Met and antitumor activity of SAPT8-Tep LYTACs in A549 cells *in vitro*. (**A-C**) Protein expression of c-Met and NCL in A549 cells after treatment with SAPT8-Tep-1 (**A**), SAPT8-Tep-2 (**B**), or SAPT8-Tep-3 (**C**) at a concentration of 800 nM for different time points (0, 6, 12, 24, 36, and 48 h). (**D-F**) Protein expression of c-Met and NCL in A549 cells after treatment with SAPT8-Tep-1 (**D**), SAPT8-Tep-2 (**E**), or SAPT8-Tep-3 (**F**) at different concentrations (0, 100, 200, 500, 800, and 1000 nM) for 36 h. (**G**) Protein expression of c-Met and NCL in A549 cells after treatment with Tep, SAPT8, SAPT8+Tep, SAPT8-Tep-1, SAPT8-Tep-2, or SAPT8-Tep-3 at a concentration of 800 nM for 36 h. (**H**) Colony formation of A549 cells after daily treatment with Tep, SAPT8, SAPT8+Tep, SAPT8-Tep-1, SAPT8-Tep-2, or SAPT8-Tep-3 at a concentration of 800 nM for 9 days. (**I**) EdU assay for proliferation of A549 cells after daily treatment with Tep, SAPT8, SAPT8+Tep, SAPT8-Tep-1, SAPT8-Tep-2, or SAPT8-Tep-3 at a concentration of 800 nM for 5 days. Nuclei were stained with DAPI. Scale bars: 100 μm. (**J**) TUNEL assay for apoptosis of A549 cells after daily treatment with Tep, SAPT8, SAPT8+Tep, SAPT8-Tep-1, SAPT8-Tep-2, or SAPT8-Tep-3 at a concentration of 800 nM for 5 days. Nuclei were stained with DAPI. Scale bars: 100 µm. Each experiment was repeated three times. Data were represented as mean ± SD, and statistical significance was calculated by the one-way ANOVA with a post hoc test (H). *\*p* < 0.05, *\*\*p* < 0.01, *\*\*\*p* < 0.001, *\*\*\*\*p* < 0.0001.

**Fig. S13.**
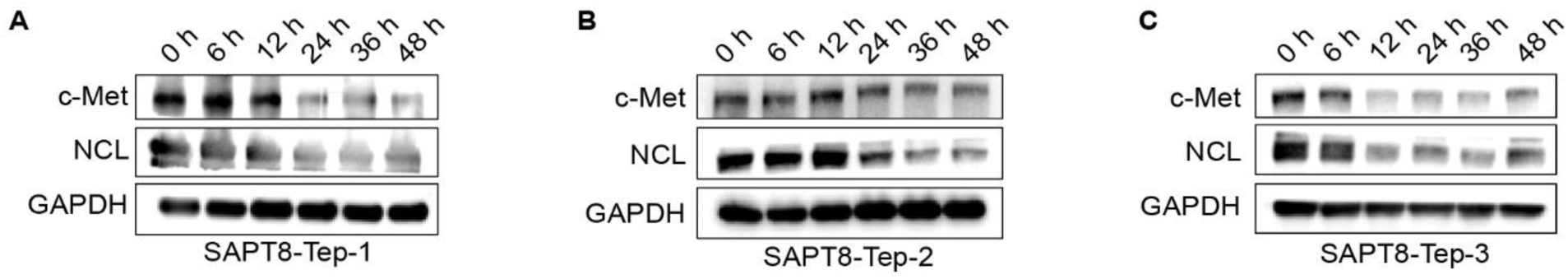
Degradation of c-Met by SAPT8-Tep LYTACs in U-2 OS cells *in vitro*. (**A-C**) Protein expression of c-Met and NCL in U-2 OS cells after treatment with SAPT8-Tep-1 (**A**), SAPT8-Tep-2 (**B**), and SAPT8-Tep-3 (**C**) at a concentration of 800 nM for different time points (0, 6, 12, 24, 36, and 48 h). Each experiment was repeated three times.

**Fig. S14.**
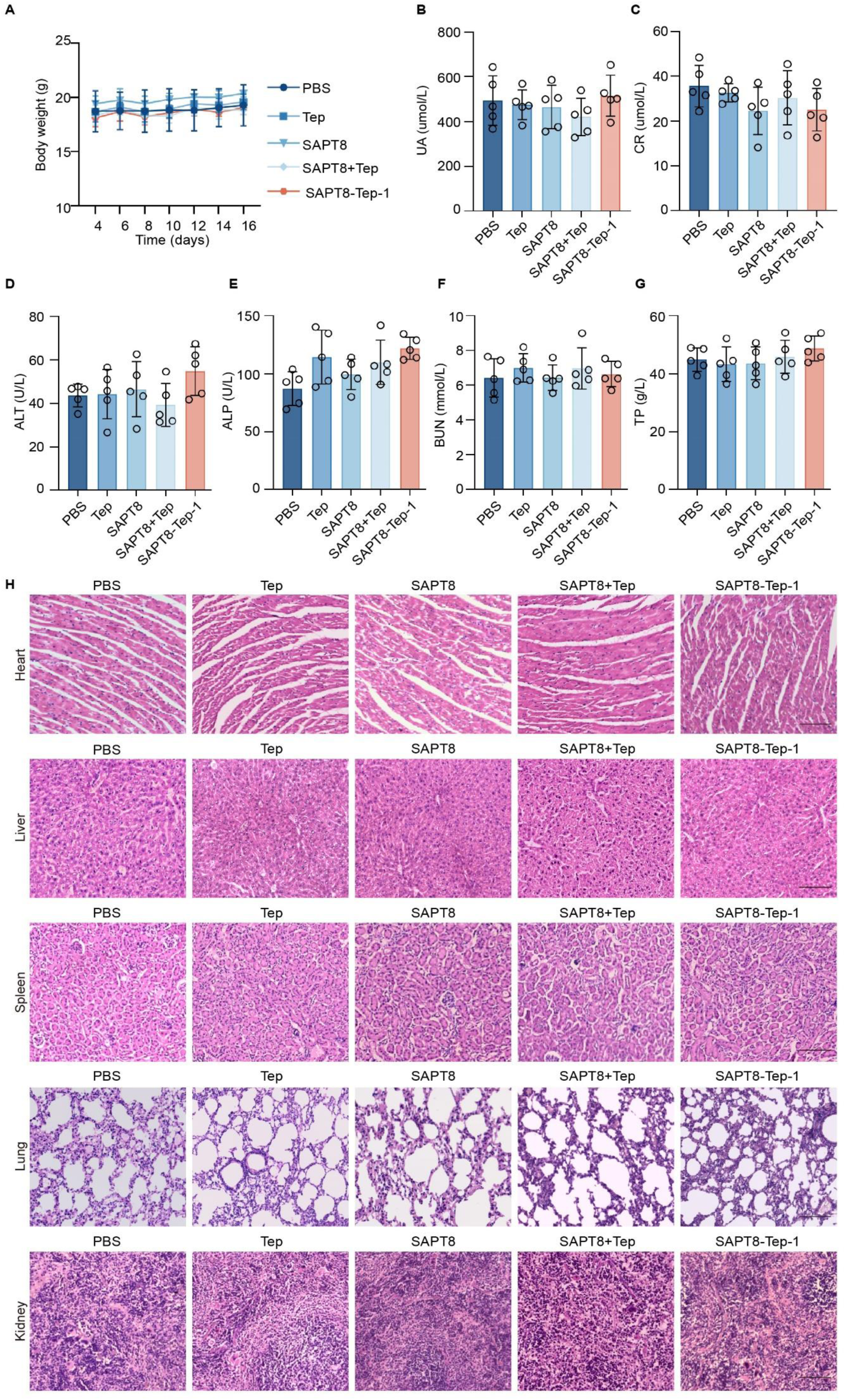
Safety of SAPT8-Tep LYTACs in a subcutaneous xenograft mouse model. (**A**) Changes in body weight of BALB/c nude mice bearing xenograft tumors in each treatment group. (**B**-**G**) Serum levels of UA (**B**), CR (**C**), ALT (**D**), ALP (**E**), BUN (**F**), and TP (**G**) in nude mice bearing subcutaneous xenograft tumors after different treatments, as measured by the automated hematology analyzer. (**H**) Histological assessments of major organs, including heart, liver, spleen, lung, and kidney, from mice in each treatment group using H&E staining. Scale bars: 100 μm. n = 5 for each group. Data were represented as mean ± SD, and statistical significance was calculated by the two-way ANOVA with a post hoc test (A) or one-way ANOVA with a post hoc test (B-G).

**Fig. S15.**
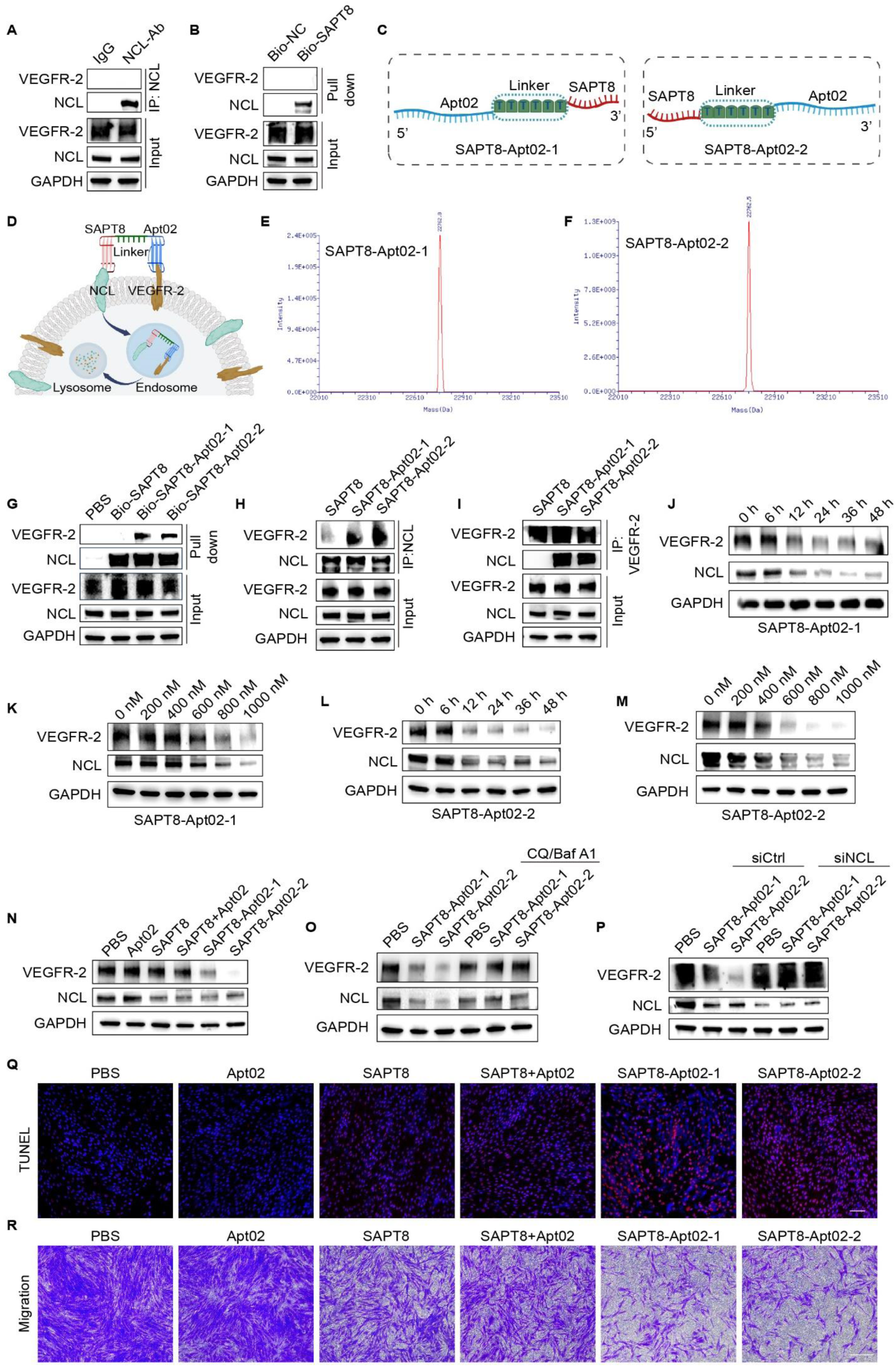
Degradation of VEGFR-2 and antitumor activity of SAPT8-Apt02 LYTACs *in vitro*. (**A**) Co-IP of VEGFR-2 and NCL in RA-FLSs using an anti-NCL antibody. (**B**) Pull-down assay of VEGFR-2 and NCL in RA-FLSs treated with 1 μM Bio-labeled NC or SAPT8 for 8 h. (**C**) Schematic diagram showing the conjugation of SAPT8 with the 5’ or 3’ terminal of VEGFR-2 aptamer Apt02 via a single-stranded DNA linker composed of six T bases, resulting in the generation of SAPT8-Apt02-1 and SAPT8-Apt02-2. (**D**) Schematic diagram illustrating the proposed mode of action of SAPT8-Apt02 LYTACs. Briefly, SAPT8-Apt02 LYTACs would selectively recognize RA-FLSs and then induce the internalization and lysosomal degradation of VEGFR-2 and NCL. (**E-F**) Mass spectrometric identification of SAPT8-Apt02-1 (**E**) and SAPT8-Apt02-2 (**F**). (**G**) Pull-down assay of VEGFR-2 and NCL in RA-FLSs treated with 1 μM biotin-labelled SAPT8, SAPT8-Apt02-1, or SAPT8-Apt02-2 for 8 h. (**H-I**) Co-IP of NCL and VEGFR-2 in RA-FLSs treated with 1 μM SAPT8, SAPT8-Apt02-1, or SAPT8-Apt02-2 for 8 h using an anti-NCL (**H**) or an anti-VEGFR-2 antibody (**I**). (**J**) Protein expression of VEGFR-2 and NCL in RA-FLSs after treatment with SAPT8-Apt02-1 at a concentration of 800 nM for different time points (0, 6, 12, 24, 36, and 48 h). (**K**) Protein expression of VEGFR-2 and NCL in RA-FLSs after treatment with SAPT8-Apt02-1 at different concentrations (0, 200, 400, 600, 800, and 1000 nM) for 24 h. (**L**) Protein expression of VEGFR-2 and NCL in RA-FLSs after treatment with SAPT8-Apt02-2 at a concentration of 800 nM for different time points (0, 6, 12, 24, 36, and 48 h). (**M**) Protein expression of VEGFR-2 and NCL in RA-FLSs after treatment with SAPT8-Apt02-2 at different concentrations (0, 200, 400, 600, 800, and 1000 nM) for 24 h. (**N**) Protein expression of VEGFR-2 and NCL in RA-FLSs after treatment with Apt02, SAPT8, SAPT8+Apt02, SAPT8-Apt02-1, or SAPT8-Apt02-2 at a concentration of 800 nM for 24 h. (**O**) Protein expression of VEGFR-2 and NCL in RA-FLSs after treatment with SAPT8-Apt02-1 or SAPT8-Apt02-2 at a concentration of 800 nM for 24 h, with or without the presence of lysosomal inhibitors CQ and Baf A1. (**P**) Protein expression of VEGFR-2 and NCL in siCtrl- or siNCL-transfected RA-FLSs after treatment with SAPT8-Apt02-1 or SAPT8-Apt02-2 at a concentration of 800 nM for 24 h. (**Q**) TUNEL assay for apoptosis of RA-FLSs after daily treatment with Apt02, SAPT8, SAPT8+Apt02, SAPT8-Apt02-1, or SAPT8-Apt02-2 at a concentration of 800 nM for 5 days. Nuclei were stained with DAPI. Scale bars: 100 μm. (**R**) Migration of RA-FLSs after daily treatment with 800 nM Apt02, SAPT8, SAPT8+Apt02, SAPT8-Apt02-1, or SAPT8-Apt02-2 for 2 days, as determined by transwell assays. Nuclei were stained with DAPI. Scale bars: 200 μm. Each experiment was repeated three times.

## Supplementary Methods

### Scheme 1. Synthetic route for SAPT8-Tep-1

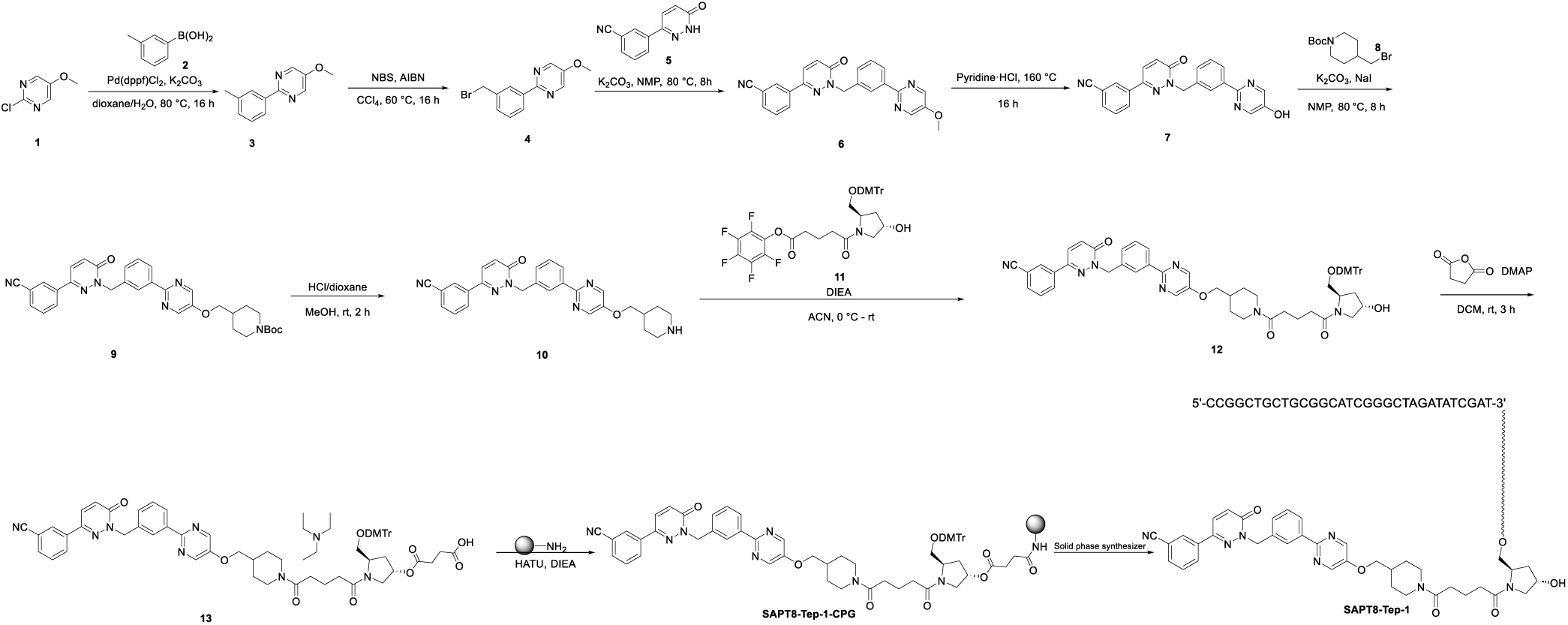

#### (1) Synthesis of compound 3

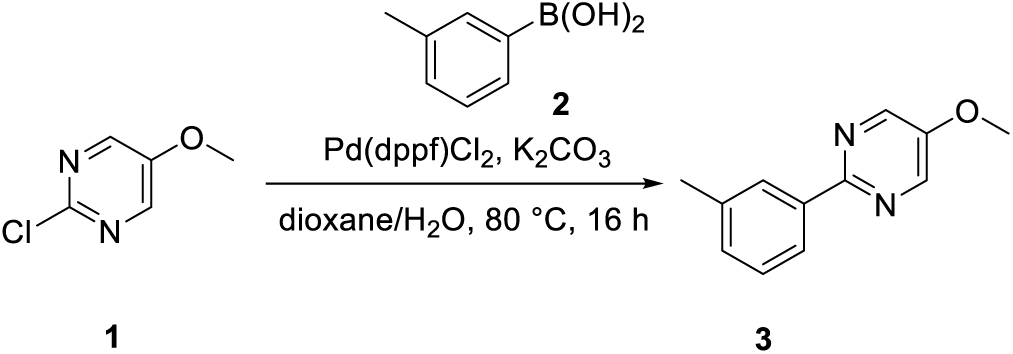

To a solution of **compound 1** (25 g, 172.94 mmol) in dioxane (450 mL) and H2O (150 mL) were added **compound 2** (23.5 g, 172.94 mmol), K2CO3 (47 g, 345.88 mmol) and Pd(dppf)Cl2 (6.2 g, 8.65 mmol). The mixture was stirred at 80 °C under nitrogen atmosphere for 16 hours. LCMS showed the reaction was complete. The mixture was diluted with water (300 mL) and extracted with EtOAc (100 mL x 3). The organic layers were combined and washed with saturated aqueous NaCl (300 mL). The organic phase was dried over anhydrous Na2SO4 and filtered. The filtrate was concentrated in vacuum. The residue was purified by flash column chromatography eluting with EtOAc in PE (0-30%) to afford **compound 3** (32 g, yield: 94%) as a brown solid. LCMS: m/z = 201.2 [M+H] ^+^, Rt = 2.13 min, purity: 96.34% (214 nm).

**^1^HNMR (400 MHz, DMSO-d6)** *δ* 8.64 - 8.62 (m, 2H), 8.20 - 8.05 (m, 2H), 7.40 - 7.36 (m, 1H), 7.29 - 7.27 (m, 1H), 3.96 (s, 3H), 2.40 (s, 3H).

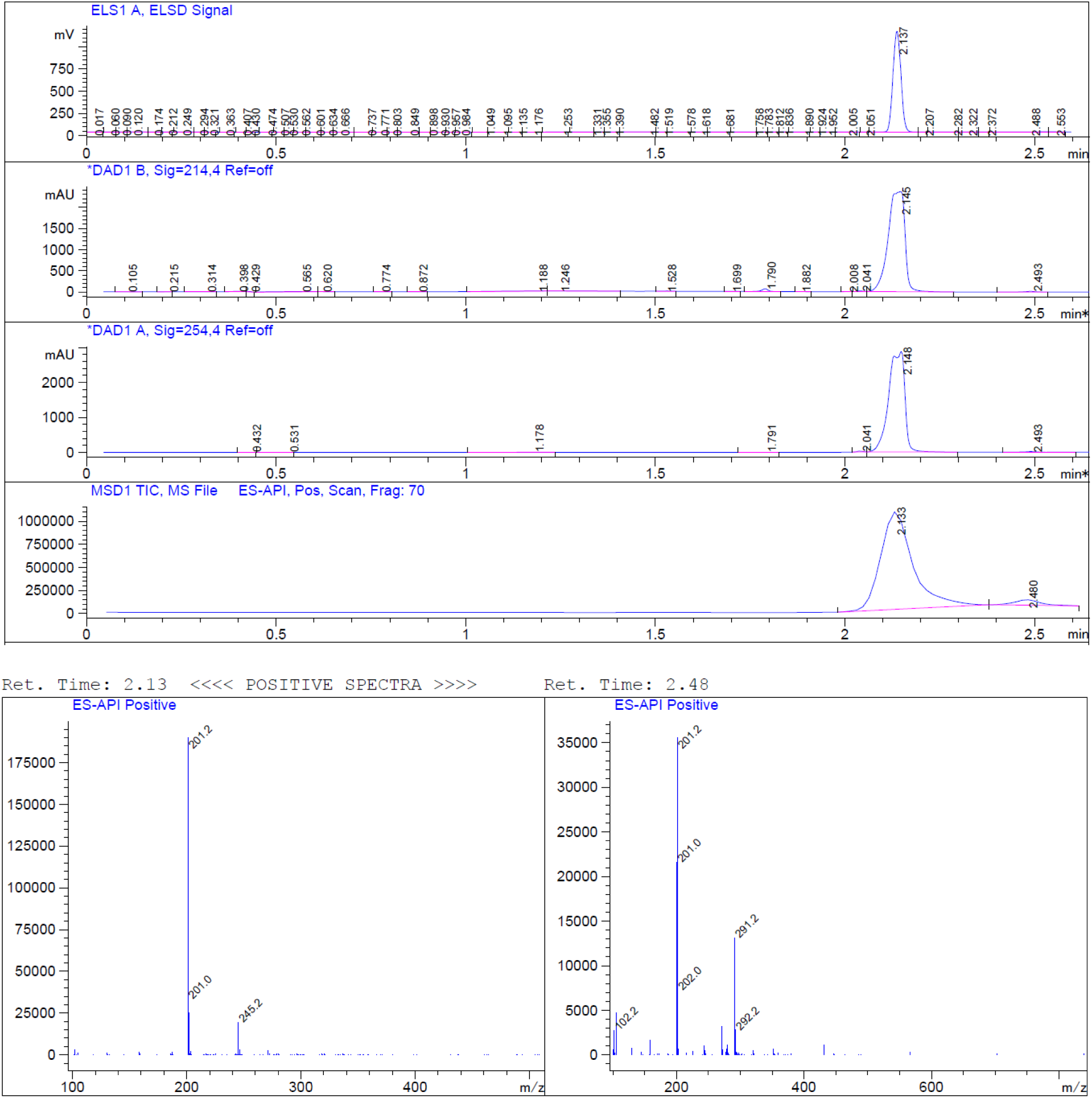

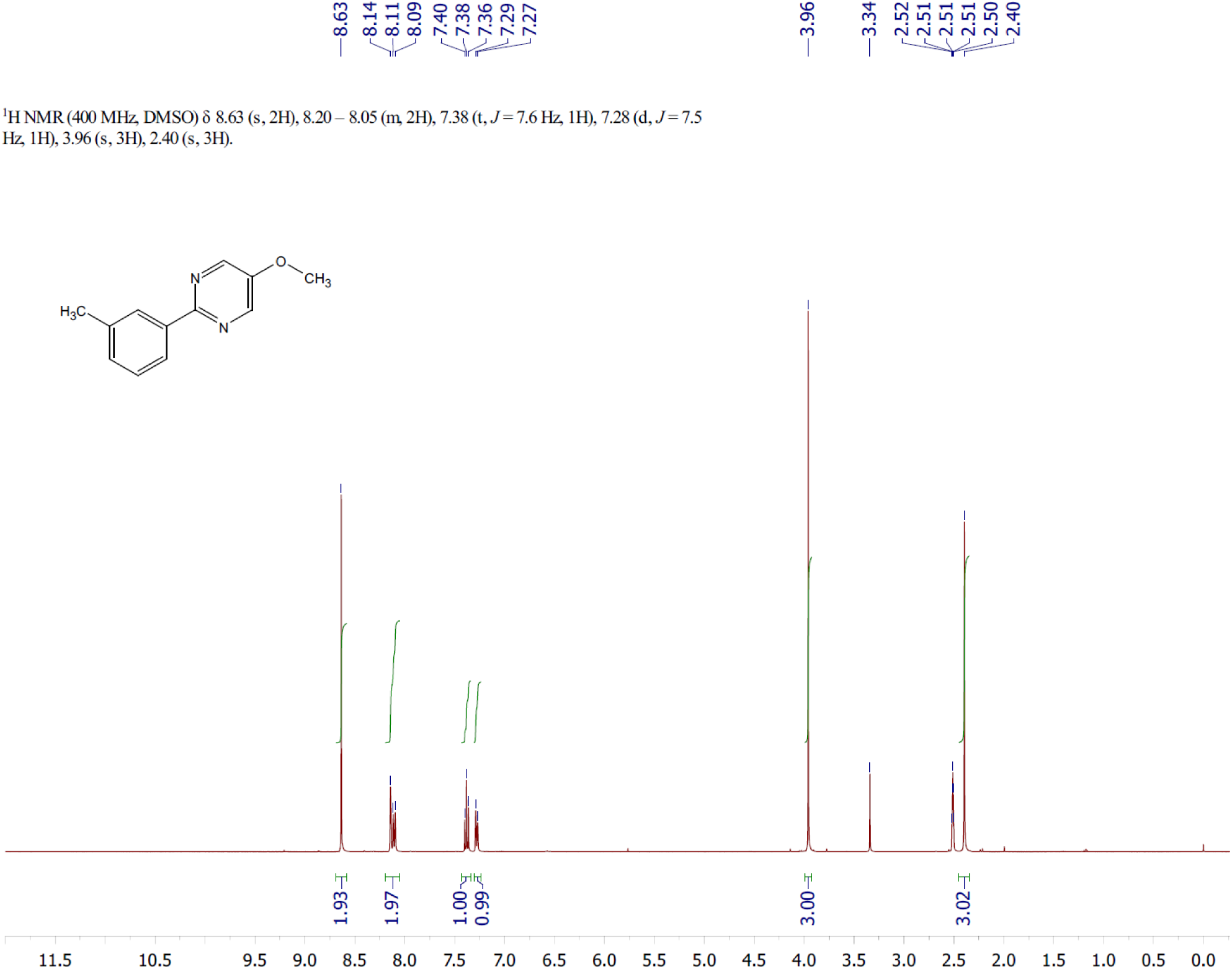

#### (2) Synthesis of compound 4

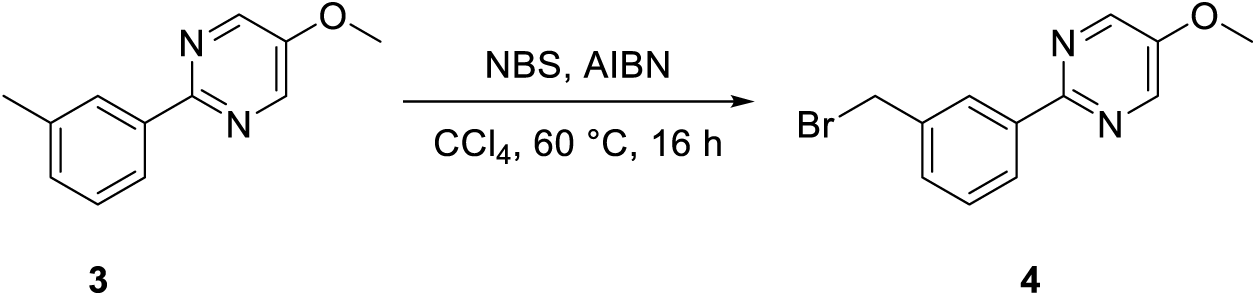

To a solution of **compound 3** (18.7 g, 93.39 mmol) and NBS (16.6 g, 93.39 mmol) in CCl4 (190 mL) was added AIBN (1.5 g, 9.34 mmol). The mixture was stirred at 60 °C under a nitrogen atmosphere for 16 hours. LCMS showed the reaction was complete. The mixture was diluted with water (500 mL) and extracted with EtOAc (150 mL x 3). The organic phase was washed with saturated aqueous NaCl (500 mL), dried over Na2SO4, and filtered. The filtrate was concentrated in vacuo to afford **compound 4** (crude, 30 g) as a brown solid. LCMS: m/z = 279.0 [M+H] ^+^, Rt = 1.657min, purity: 72.41% (214 nm).

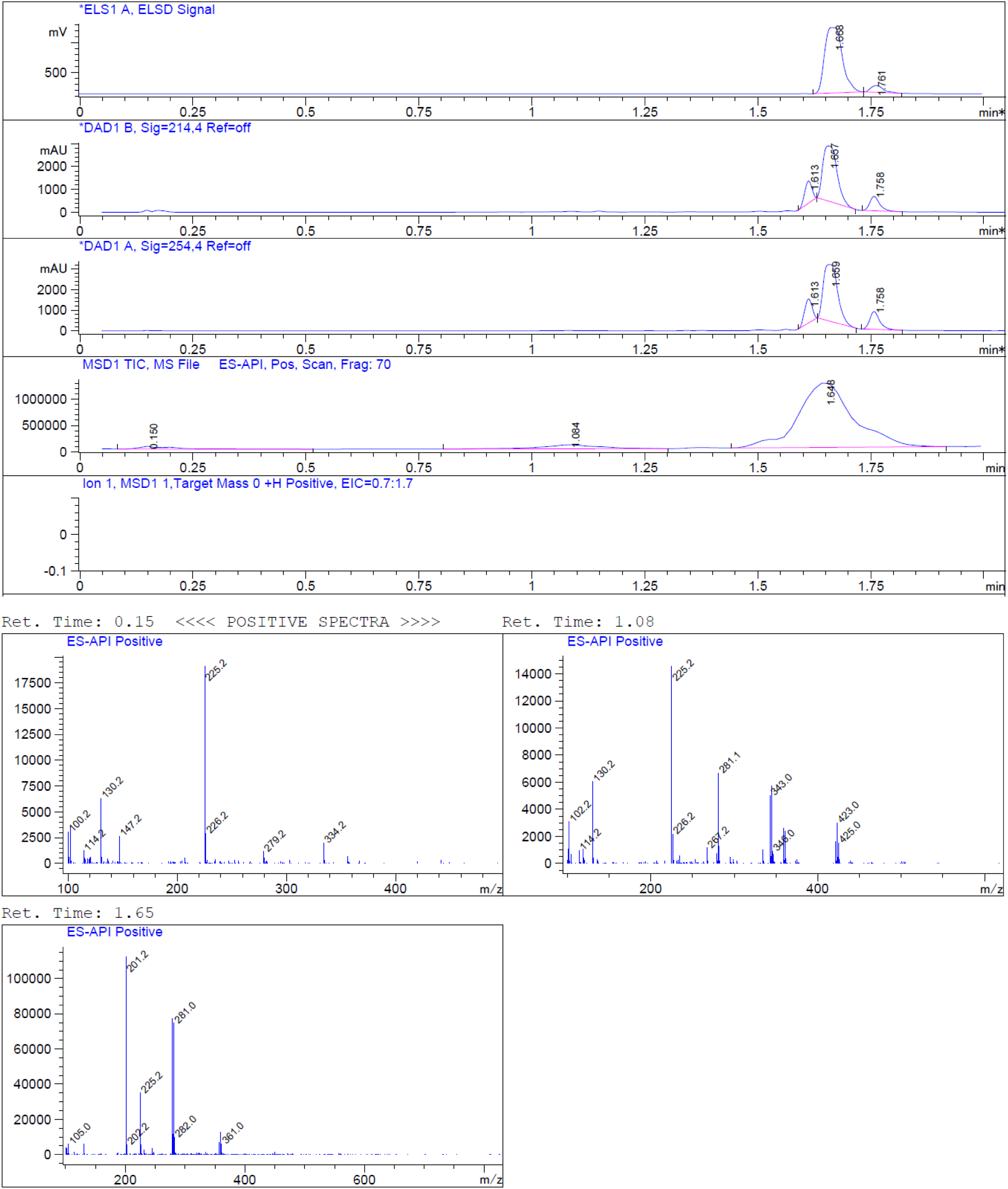

#### (3) Synthesis of compound 6

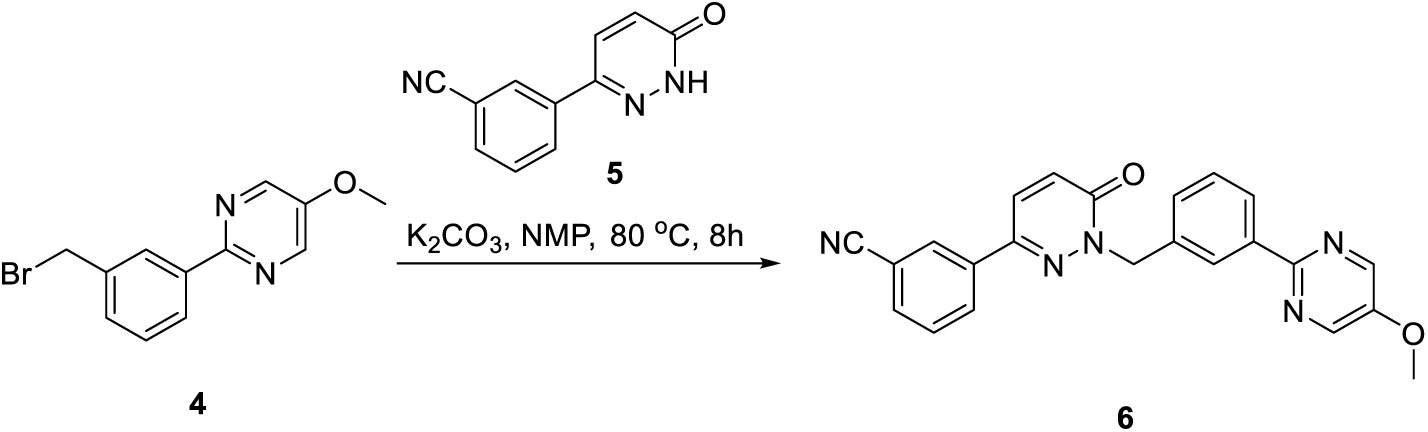

To a stirred solution of **compound 4** (30 g, 107.47 mmol) and K2CO3 (16.3 g, 118.22 mmol) in NMP (250 mL) was added **compound 5** (16.9 g, 85.98 mmol). The mixture was stirred at 80 °C for 16 hours. LCMS showed the reaction was completed. The reaction was quenched by water (1000 mL). The mixture was filtered. The wet filter cake was recrystallized from EtOAc (300 mL) to give **compound 6** as an off-white solid (26 g, crude). LCMS: m/z = 396.0 [M+H] ^+^, Rt = 1.627 min, purity: 88.03% (214 nm)

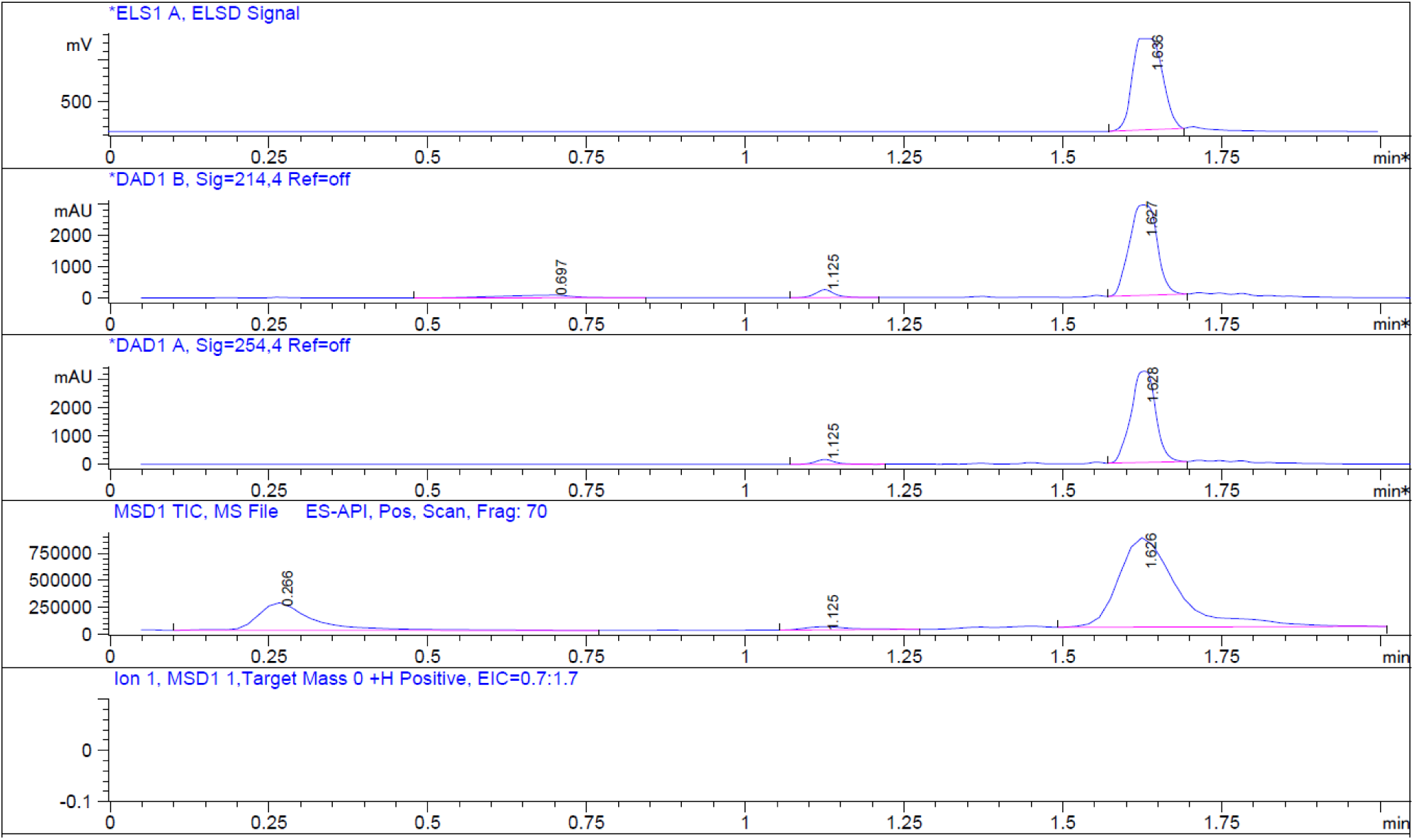

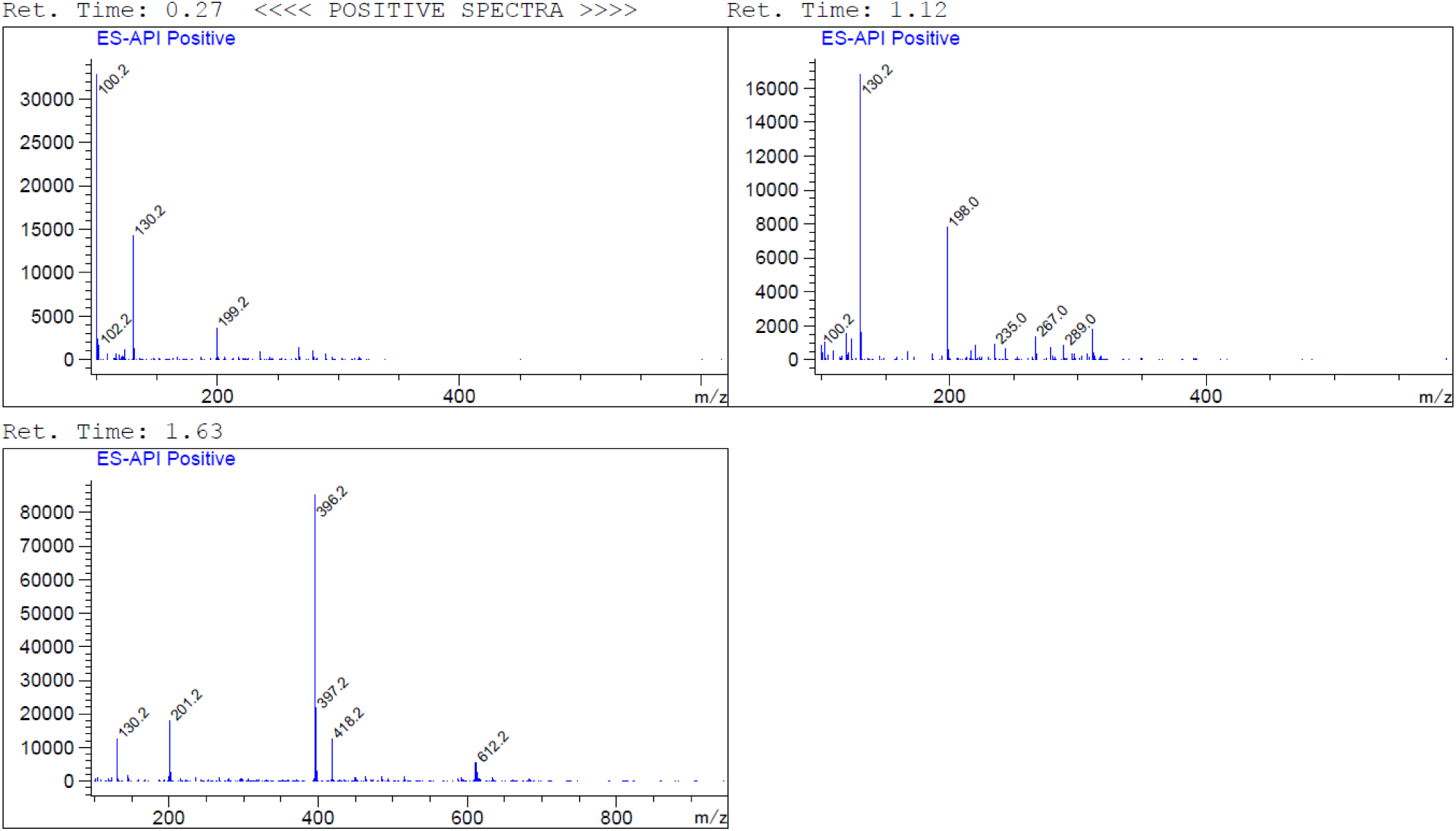

#### (4) Synthesis of compound 7

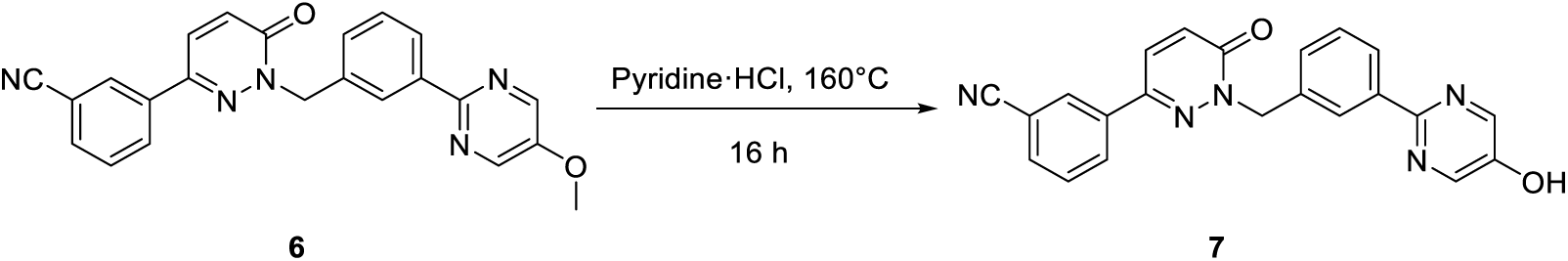

**Compound 6** (26 g, 65.25 mmol) was added pyridine ·HCl. The mixture was stirred at 160 °C for 16 hours. LCMS showed the reaction was completed. The reaction was quenched by water (200 mL). The mixture was filtered. The wet filter cake was recrystallized from EtOAc (200 mL) to give **compound 7** as an off-white solid (31.6 g, crude).

LCMS: m/z = 382.2 [M+H] ^+^, Rt = 1.486 min, purity: 63.38% (214 nm)

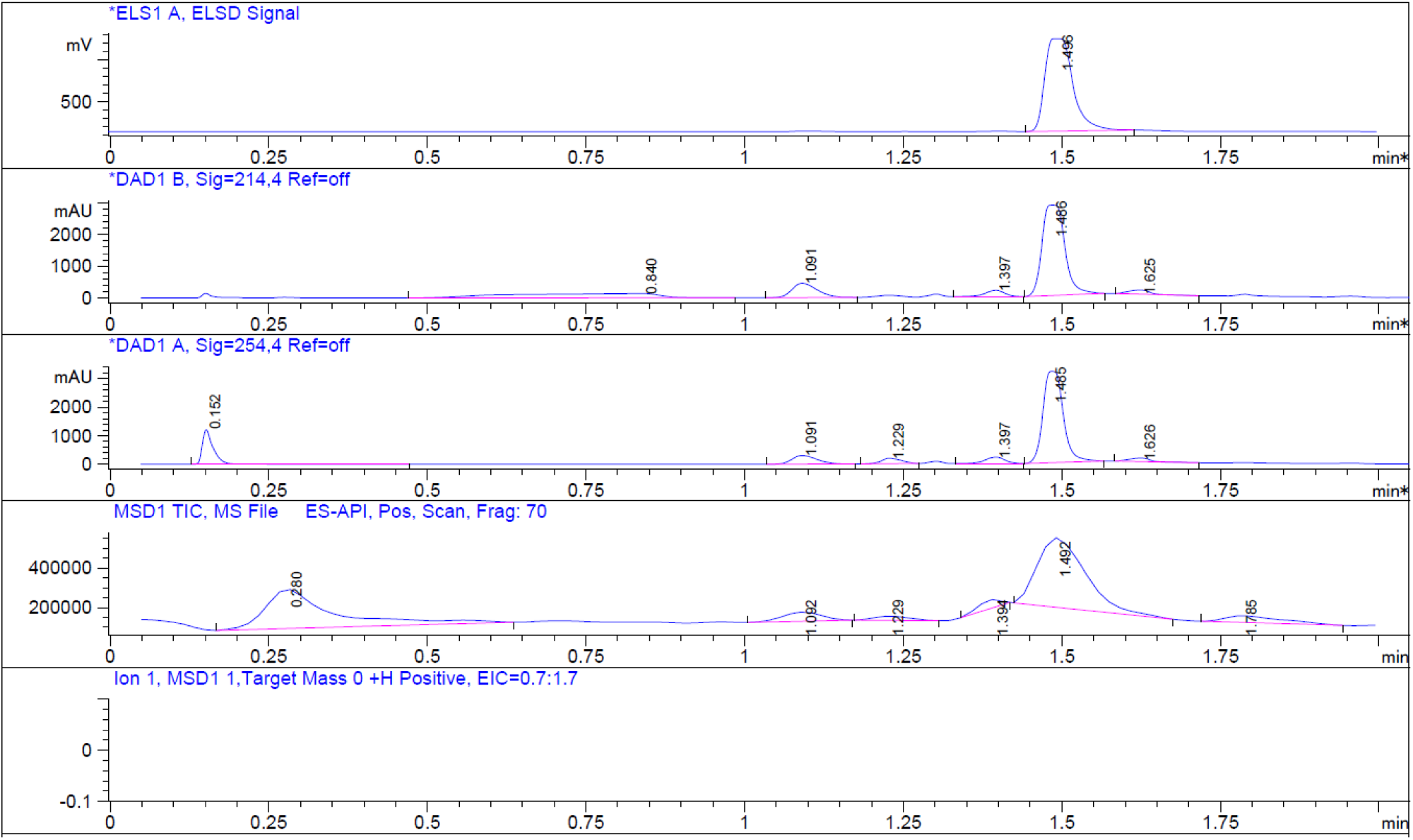

#### (5) Synthesis of compound 9

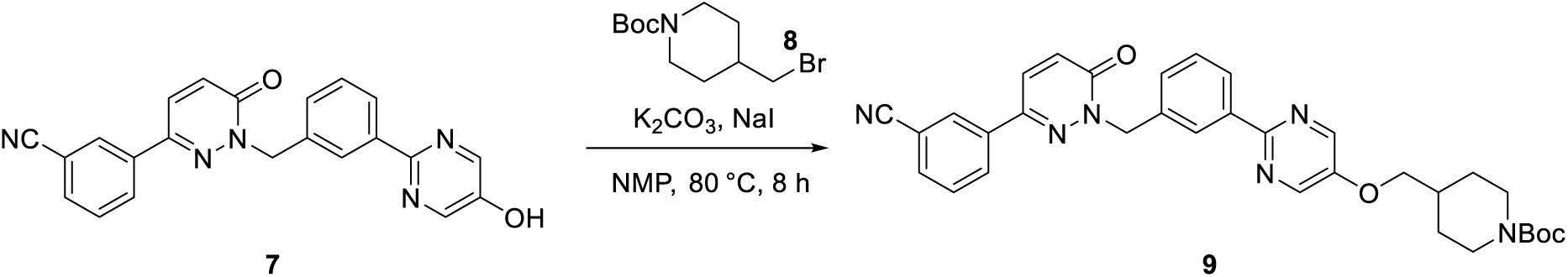

To a suspension of **compound 7** (16.6 g, 43.52 mmol) in NMP (200 mL) were added NaI (0.65 g, 4.35 mmol), K2CO3 (11.75 g, 85.04 mmol), and compound 8 (16.95 g, 60.93 mmol). The reaction was stirred at 80 °C for 8 hours. LCMS showed the reaction was complete. The reaction was quenched by water (800 mL). The mixture was filtered. The wet filter cake was recrystallized from EtOAc (200 mL) to give **compound 9** as an off-white solid (11 g, crude).

LCMS: m/z = 523.2 [M-55] ^+^, Rt = 1.885 min, purity: 91.46% (214nm)

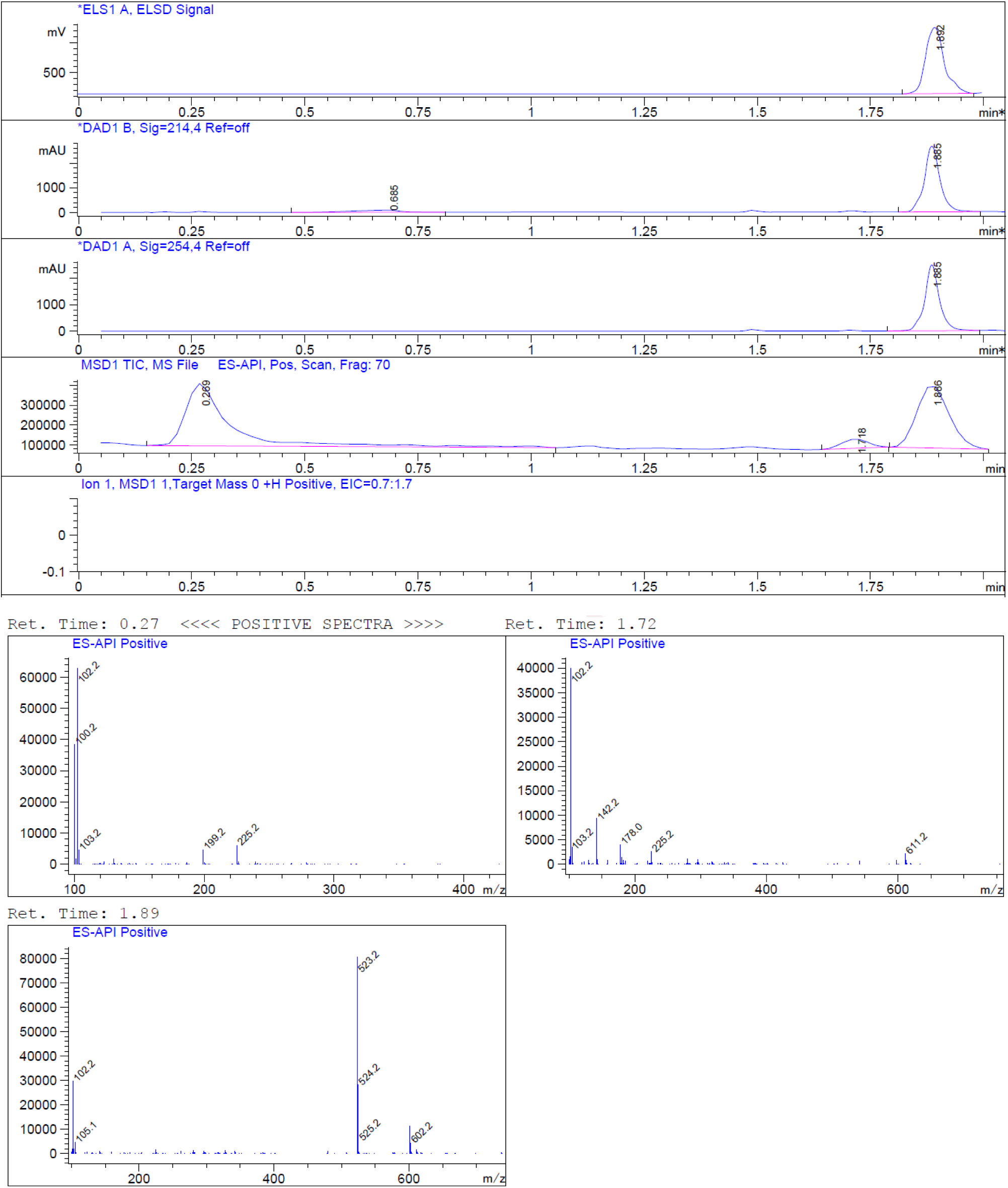

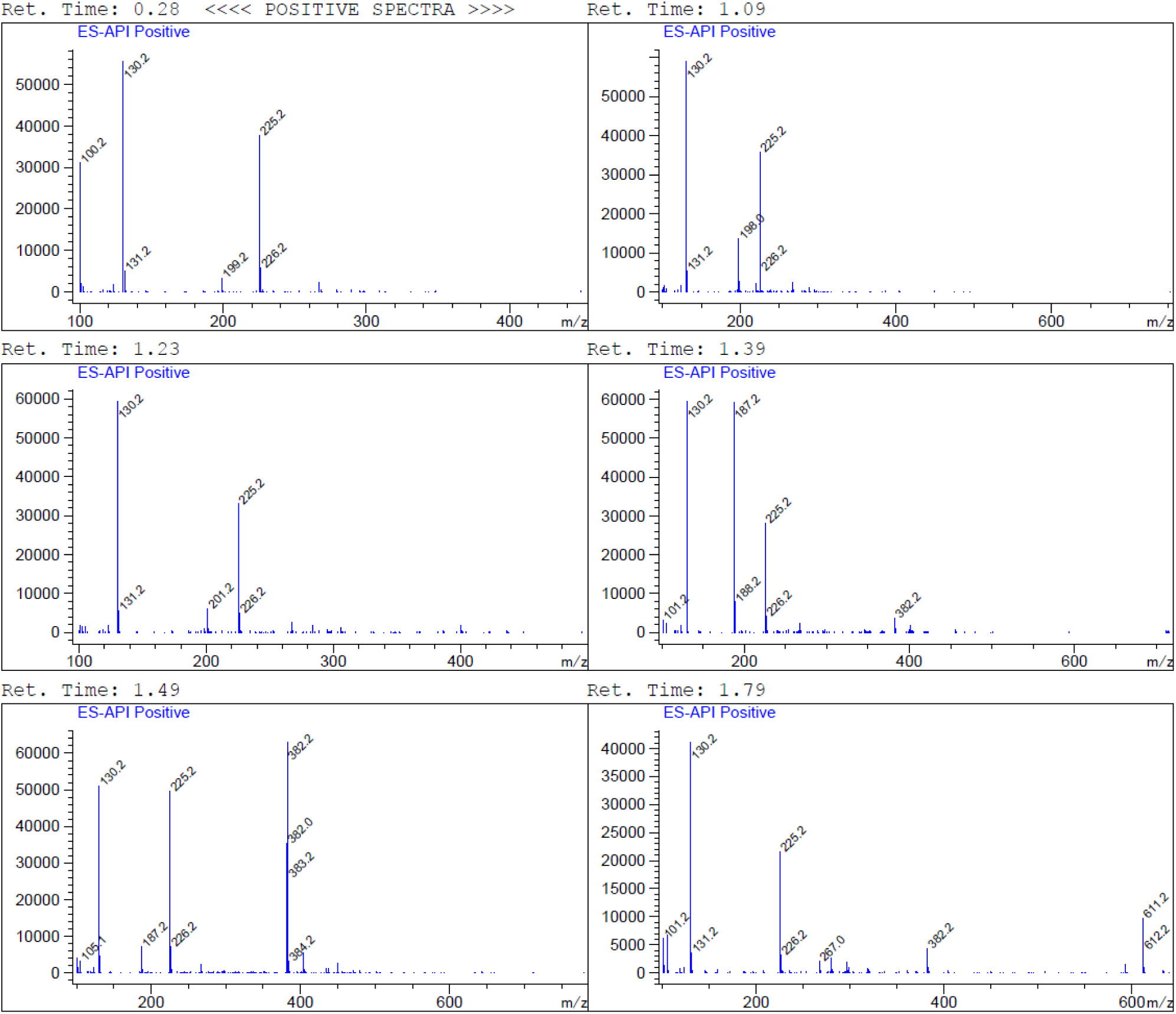

#### (6) Synthesis of compound 10

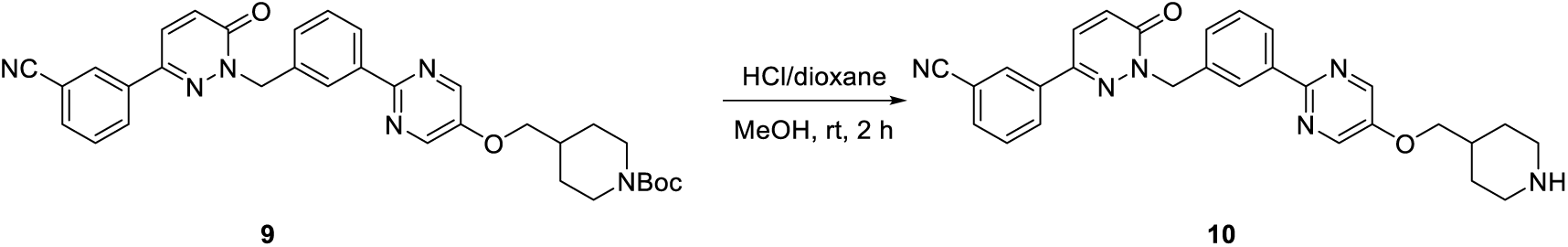

To a solution of **compound 9** (4 g, 6.91 mmol) in DCM (40 mL) was added dropwise TFA (10 mL). The resulting solution was stirred at room temperature for 2 hours. LCMS showed the reaction was complete. The mixture was concentrated in a vacuum to give the crude **compound 10** (4 g, crude), which was used for the next step without further purification.

LCMS: m/z = 479.2 [M+H] ^+^, Rt = 1.373 min, purity: 52.07% (214nm)

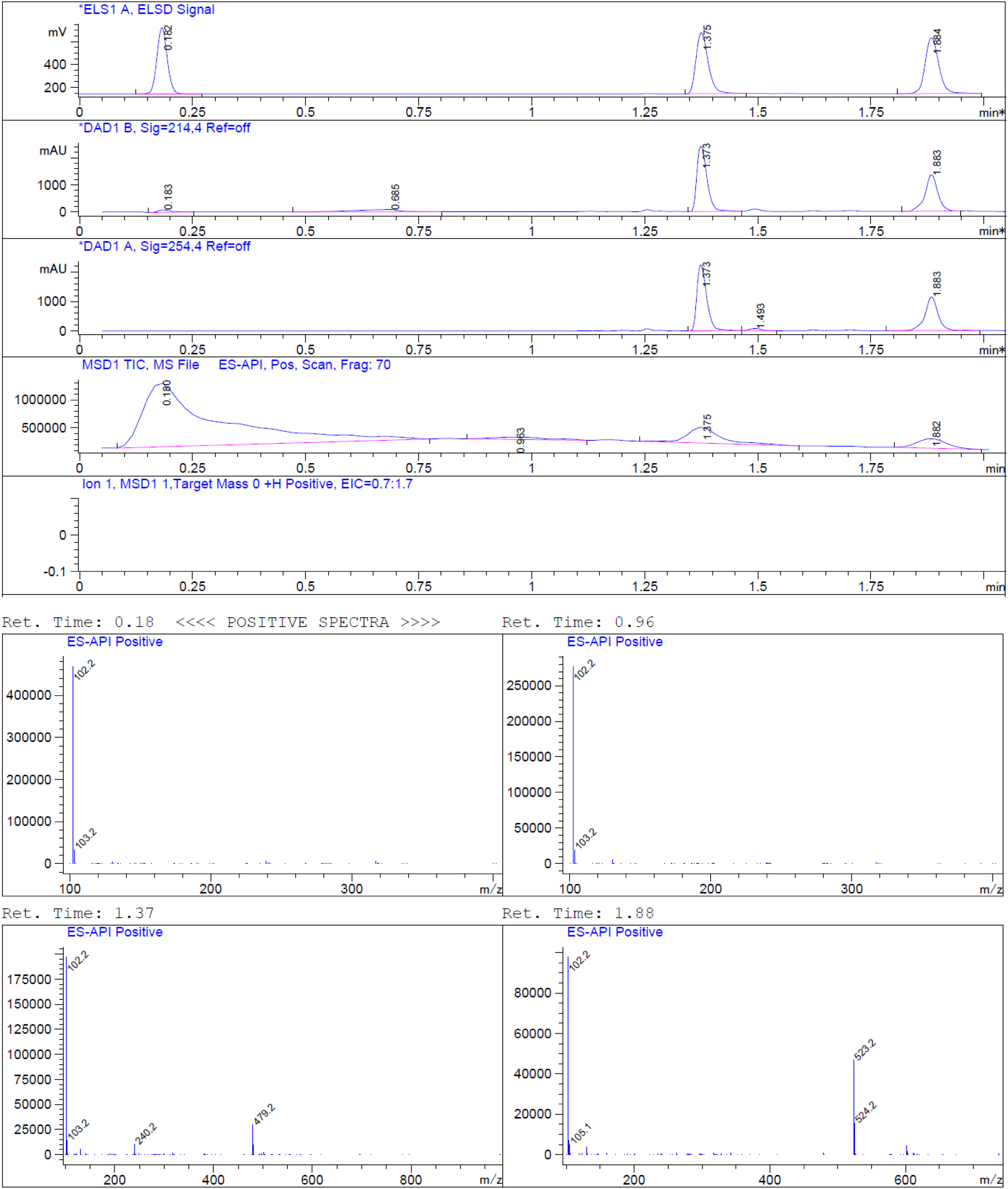

#### (7) Synthesis of compound 12

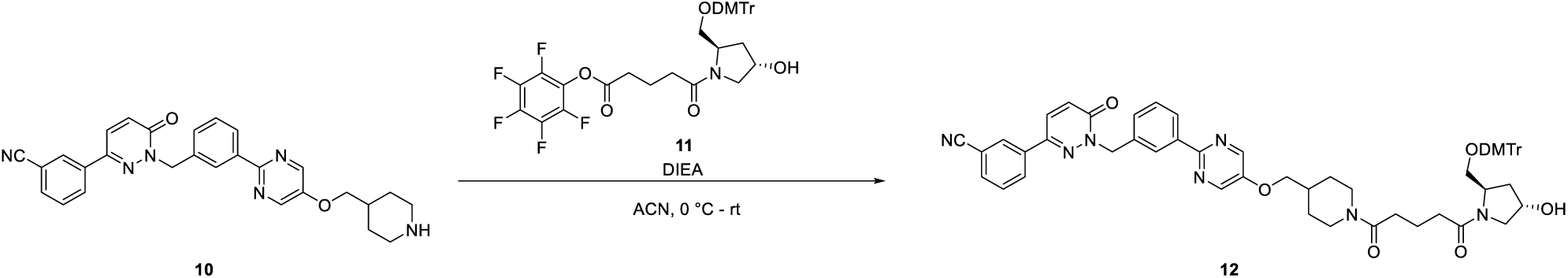

To a solution of **compound 10** (2.0 g, 4.29 mmol) and DIEA (5.5 g, 42.9 mmol) in ACN (20 mL) was added dropwise a solution of **compound 11** (3 g, 4.29 mmol) in ACN (10 mL) at 0 °C under nitrogen. The reaction was stirred at room temperature for 10 hours. LCMS showed the reaction was complete. The mixture was concentrated in a vacuum. The residue was purified by flash column chromatography (MeOH in DCM, 0% to 10%) to give **compound 12** (3 g, crude) as a white solid.

LCMS: m/z= 1016.7 [M+Na] ^+^, Rt = 2.145 min, purity: 54.68% (214 nm)

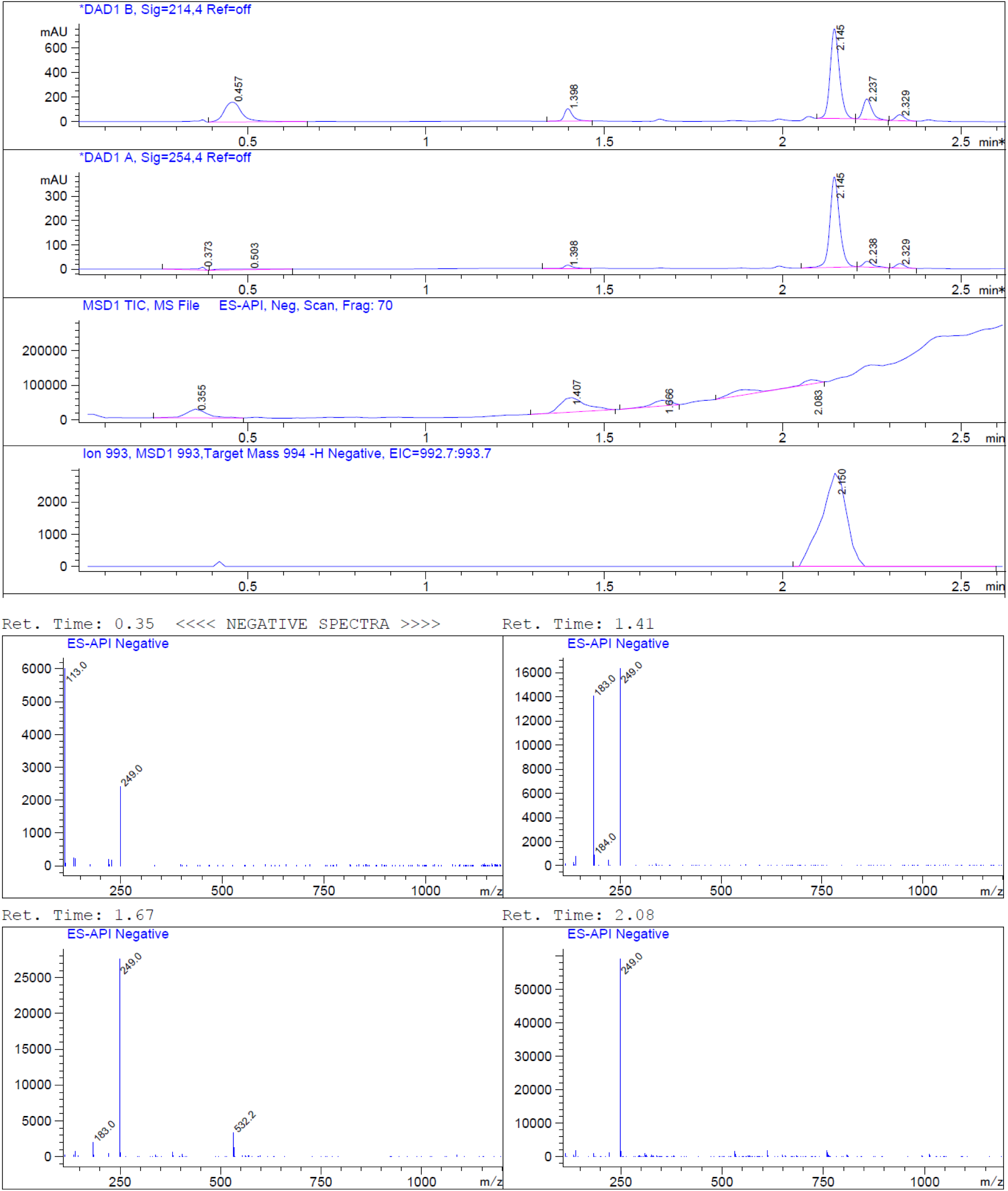

#### (8) Synthesis of compound 13

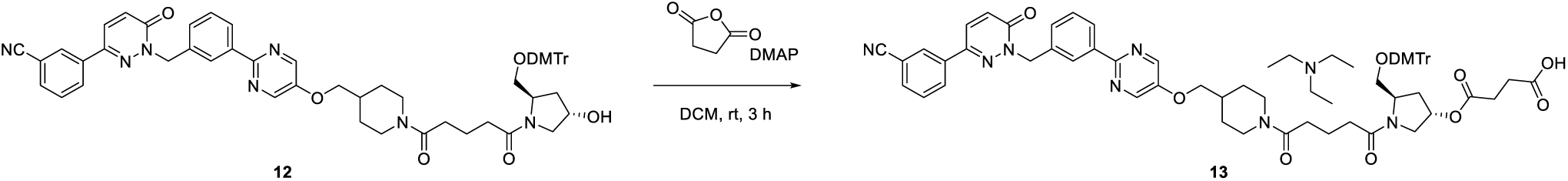

To a solution of **compound 12** (3 g, 3.02 mmol) in DCM (30 mL) were added dihydrofuran-2,5-dione (603 mg, 6.04 mmol) and DMAP (652 mg, 6.04 mmol). The reaction was stirred at room temperature for 3 hours. LCMS showed the reaction was complete. The reaction mixture was concentrated under vacuum. The residue was purified by reversed-phase column chromatography to give **compound 13** as a white solid (1.5 g, yield: 45.43%).

LCMS: m/z= 1116.9 [M+Na] ^+^, Rt = 7.86 min, purity: 100% (214 nm)

**^1^H NMR (400 MHz, DMSO-*d_6_*)** *δ* 8.67 - 8.60 (m, 2H), 8.38 (d, *J* = 1.7 Hz, 2H), 8.27 - 8.20 (m, 2H), 8.17 (d, *J* = 9.8 Hz, 1H), 7.93 (d, *J* = 7.8 Hz, 1H), 7.71 (t, *J* = 7.9 Hz, 1H), 7.52 - 7.46 (m, 2H), 7.36 - 7.26 (m, 4H), 7.25 - 7.16 (m, 6H), 6.87 (d, *J* = 8.6 Hz, 4H), 5.44 (s, 2H), 5.37 (s, 1H), 4.42 (d, *J* = 12.2 Hz, 1H), 4.22 (s, 1H), 4.04 (d, *J* = 5.7 Hz, 2H), 3.92 - 3.76 (m, 2H), 3.73 (s, 6H), 3.59-3.53 (m, 1H), 3.45 (s, 3H), 3.26 (s, 3H), 3.09 (s, 1H), 2.99-2.96 (m, 2H), 2.58-2.55 (m, 1H), 2.48 – 2.41 (m, 8H), 2.39 - 2.28 (m, 3H), 2.26 - 2.14 (m, 2H), 2.10 - 1.97 (m, 2H), 1.80 - 1.70 (m, 3H), 1.24 - 1.09 (m, 2H), 0.94 (t, J = 7.1 Hz, 7H).

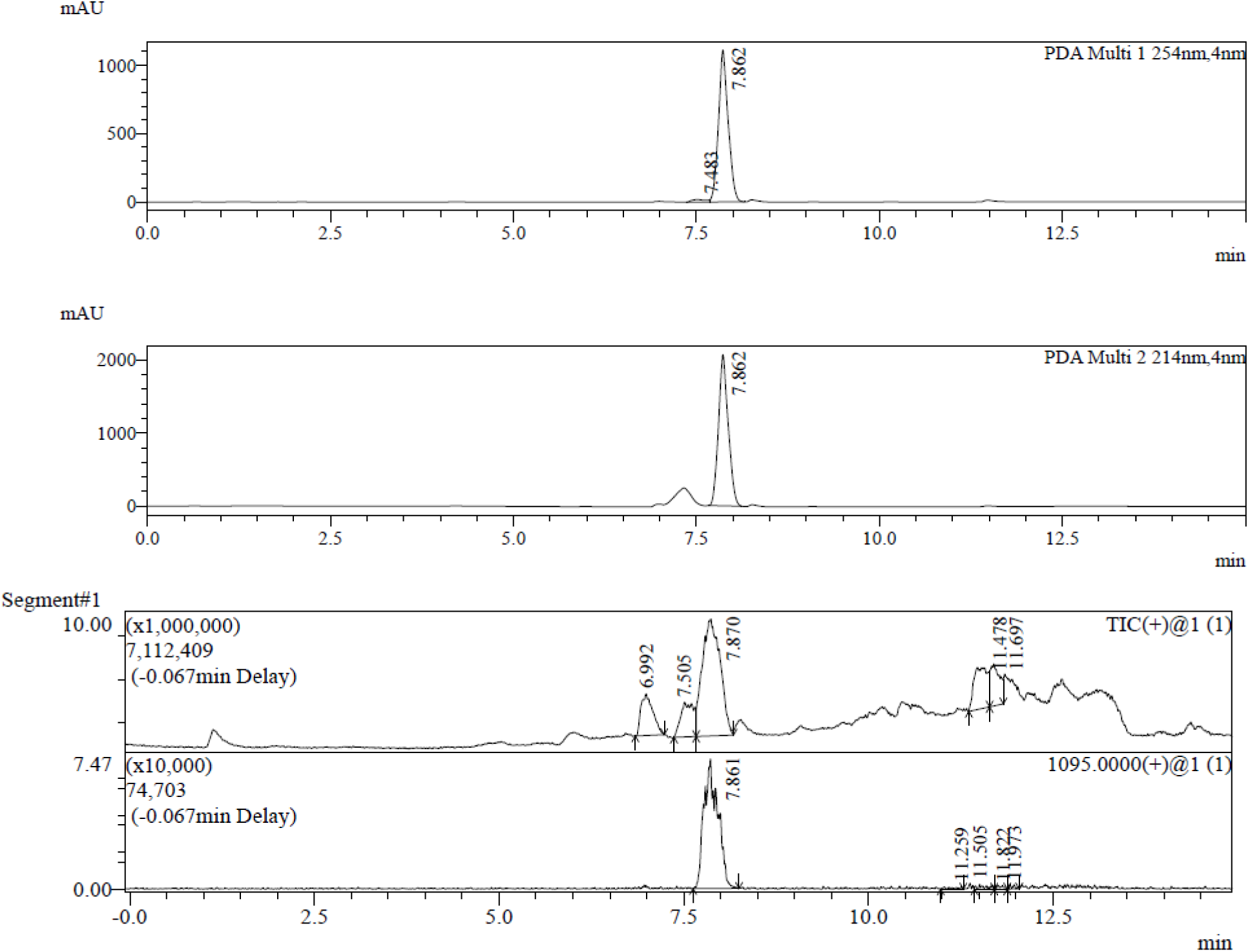

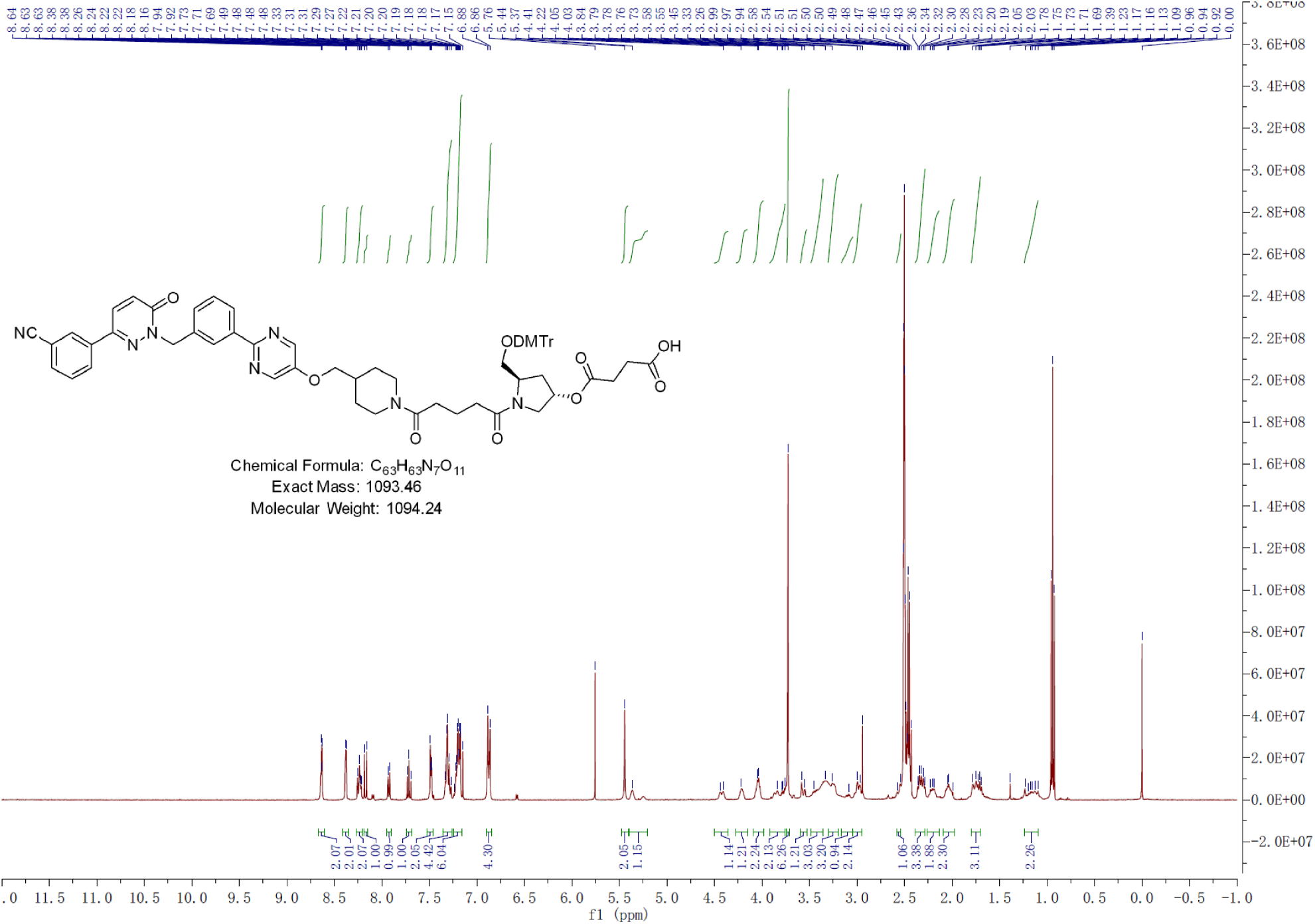

#### (9) Synthesis of SAPT8-Tep-1-CPG

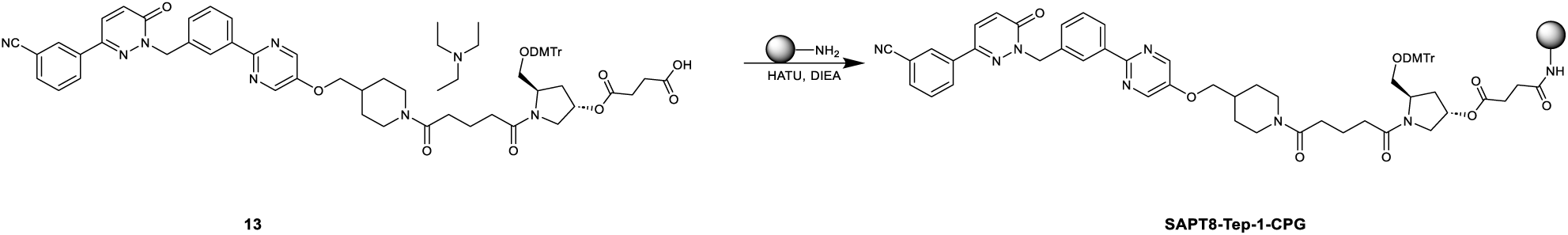

To a solution of **compound 13** (500 mg, 456,9 μmol) in ACN (30 mL) were added HATU (200 mg, 526 μmol), DIEA (200 μL), lcaa-CPG (1000 °A, 2500 mg), respectively, at room temperature and oscillated for 12 hours. After the reaction was completed, CPG was washed with ACN, CAP A (acetic anhydride: tetrahydrofuran= 1:9, v/v, 200 mL), and CAP B (n-methylimidazole: pyridine: acetonitrile= 15:10:75, v/v/v, 200 mL) were added, and the mixture was oscillated at room temperature for 1 hour. Then the mixture was filtered and washed with ACN (300 mL) 3 times. **SAPT8-Tep-1-CPG** was obtained after freeze-drying the mixture as a white powder (2500 mg).

#### (10) Synthesis of SAPT8-Tep-1

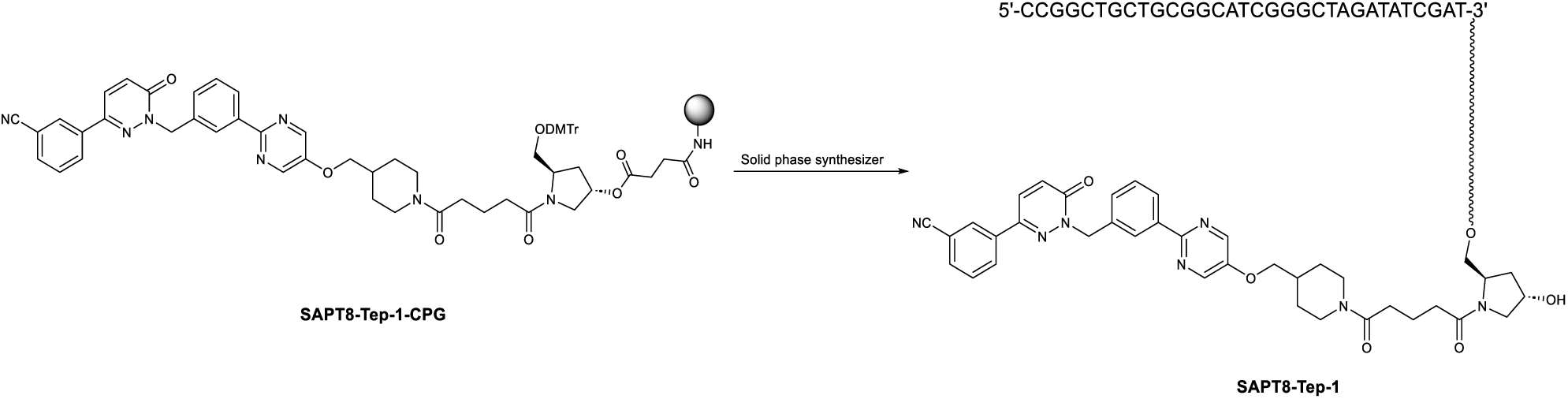

**SAPT8-Tep-1-CPG** in synthesis columns (60 mg*24) and synthesize by K-A H-8 Solid phase synthesizer. Solid phase synthesis involves four steps: Detritylation, Coupling, Capping and Oxidation. After the reaction was done, CPG of each synthesis column was added 1.5 mL aqueous ammonia and heated in an oven at 65 °C for 16 hours. Then collected the supernatant and washed with water (1 mL*3). The combined crude was purified by RP-IP-HPLC (WATERS 2489, 3767) (**Column**: XBridge BET C18 2.5μm. **RP-IP-HPLC method:** Mobile phase A: 2 % HFIP +0.1 % TEA in DD water, Mobile phase B: MeOH) to afford **SAPT8-Tep-1** as a white powder (32.62 mg, purity = 98.104%).

UPLC-MS (WATERS ACQUITY PREMIER): **SAPT8-Tep-1-UPLC**, *m/z* = 10611.21249 [M]^-^(deconvolution); *t*_R_ = 10.79 min (260 nm). Mass error <50 ppm.

HPLC (SPT-M40): **SAPT8-Tep-1-HPLC**, *t*_R_ = 12.130 min (260 nm), purity: 98.104%.

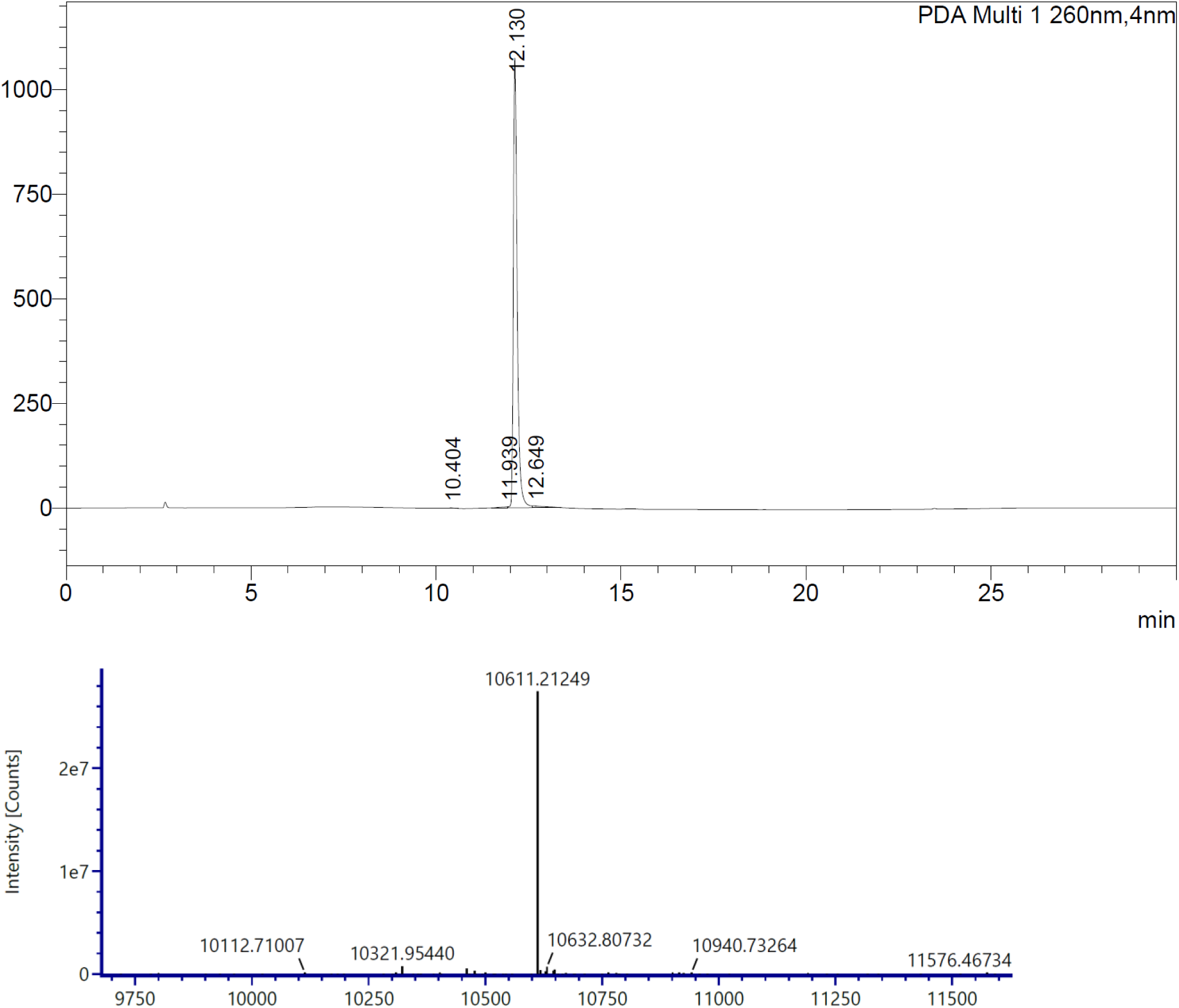

### Scheme 2. Synthetic route for SAPT8-Tep-2

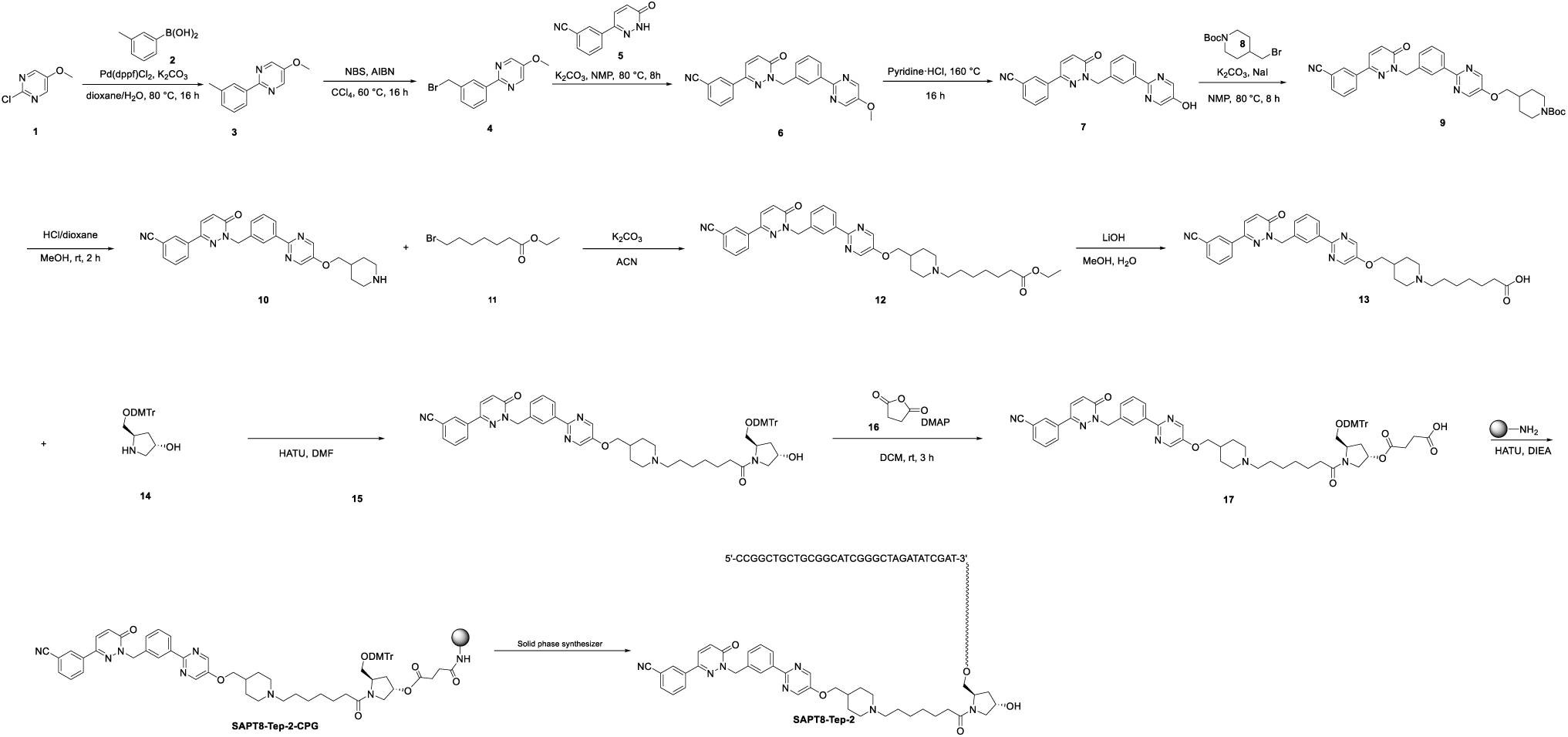

#### (1) Synthesis of compound 3

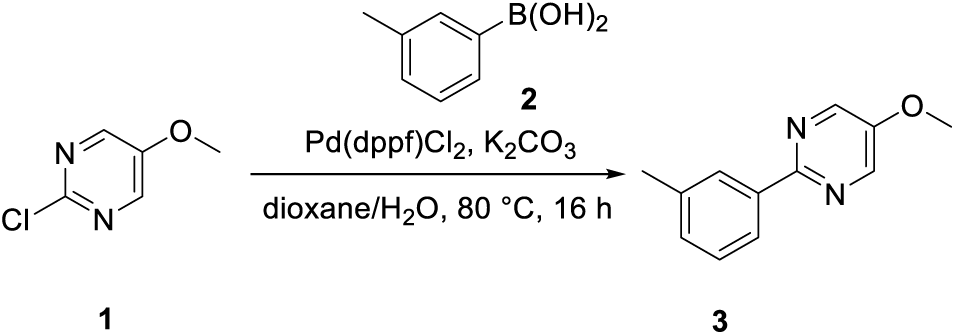

To a solution of **compound 1** (25 g, 172.94 mmol) in dioxane (450 mL) and H2O (150 mL) were added **compound 2** (23.5 g, 172.94 mmol), K2CO3 (47 g, 345.88 mmol) and Pd(dppf)Cl2 (6.2 g, 8.65 mmol). The mixture was stirred at 80 °C under nitrogen atmosphere for 16 hours. LCMS showed the reaction was complete. The mixture was diluted with water (300 mL) and extracted with EtOAc (100 mL x 3). The organic layers was combined and washed with saturated aqueous NaCl (300 mL). The organic phase was dried over anhydrous Na2SO4 and filtered. The filtrate was concentrated in vacuum. The residue was purified by flash column chromatography eluting with EtOAc in PE (0-30%) to afford **compound 3** (32 g, yield: 94%) as a brown solid. LCMS: m/z = 201.2 [M+H] ^+^, Rt = 2.13 min, purity: 96.34% (214 nm).

**^1^HNMR (400 MHz, DMSO-*d*_6_)** *δ* 8.64 - 8.62 (m, 2H), 8.20 - 8.05 (m, 2H), 7.40 - 7.36 (m, 1H), 7.29 - 7.27 (m, 1H), 3.96 (s, 3H), 2.40 (s, 3H).

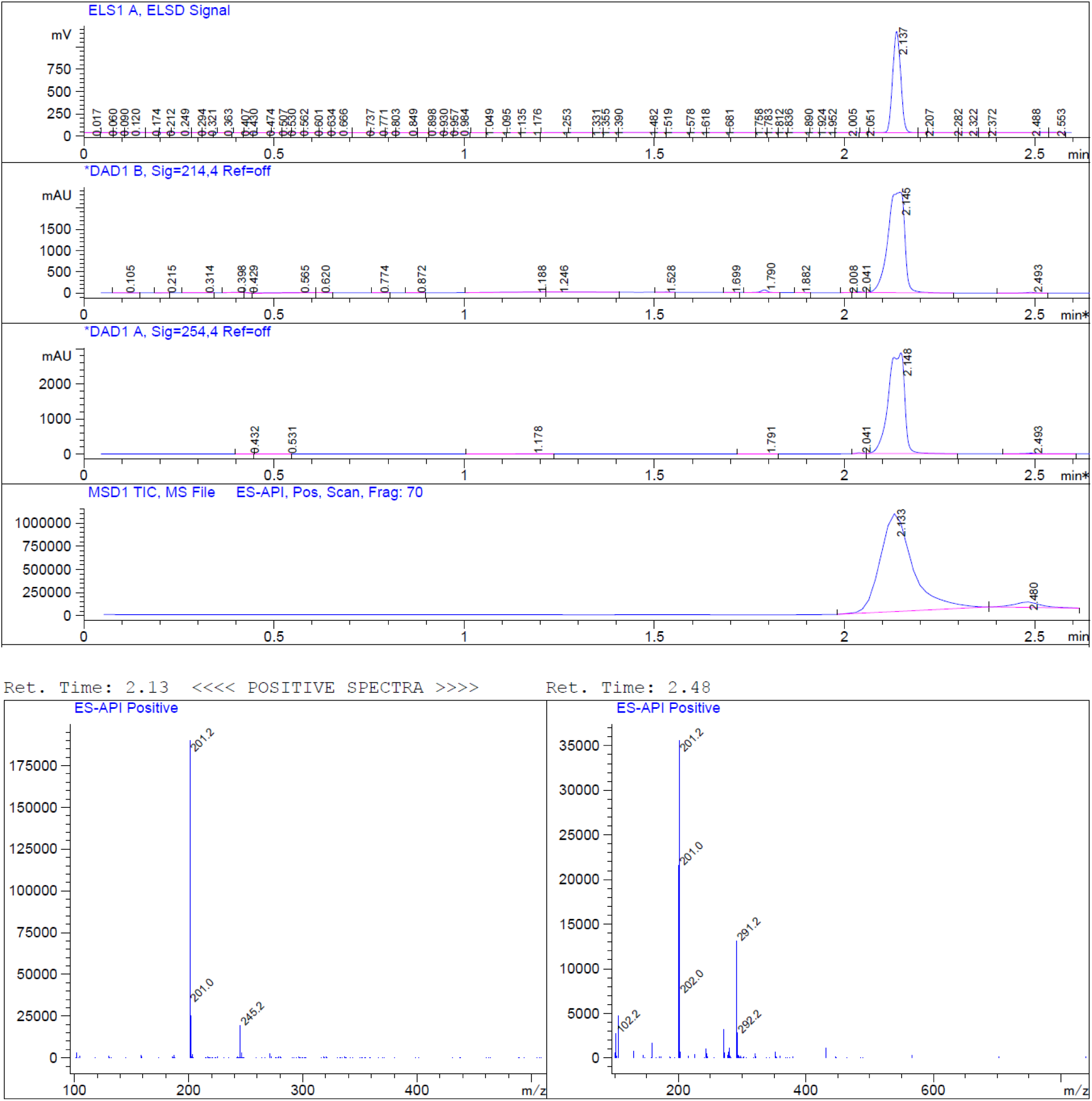

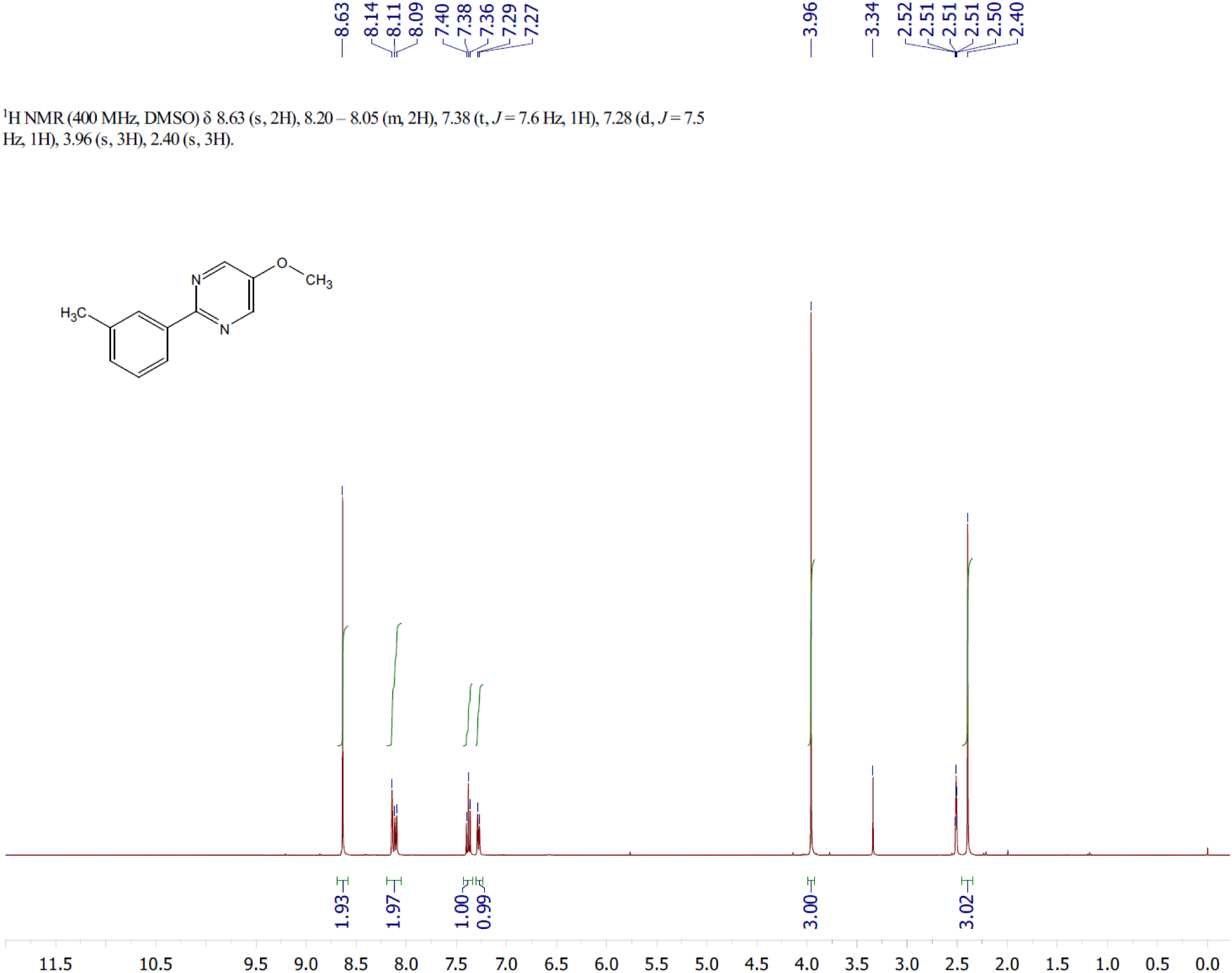

#### (2) Synthesis of compound 4

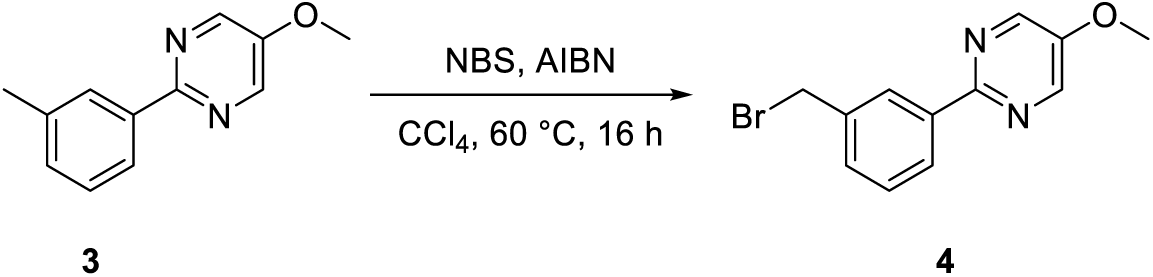

To a solution of **compound 3** (18.7 g, 93.39 mmol) and NBS (16.6 g, 93.39 mmol) in CCl4 (190 mL) were added AIBN (1.5 g, 9.34 mmol). The mixture was stirred at 60 °C under nitrogen atmosphere for 16 hours. LCMS showed the reaction was complete. The mixture was diluted with water (500 mL) and extract with EtOAc (150 mL x 3). The organic phase was washed with saturated aqueous NaCl (500 mL), dried over Na2SO4 and filtered. The filtrate was concentrated in vacuo to afford **compound 4** (crude, 30 g) as a brown solid.

LCMS: m/z = 279.0 [M+H] ^+^, Rt = 1.657min, purity: 72.41% (214 nm).

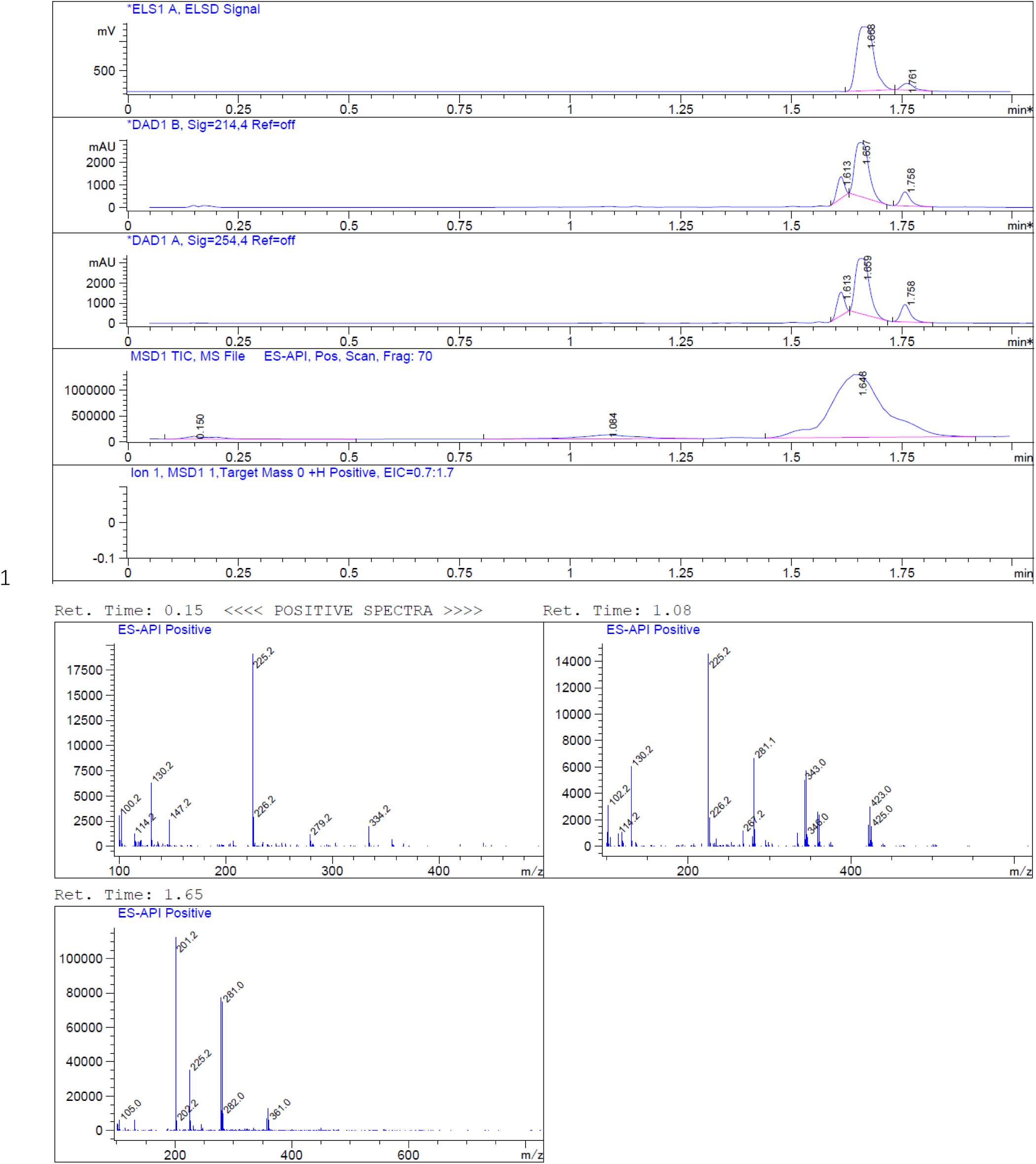

#### (3) Synthesis of compound 6

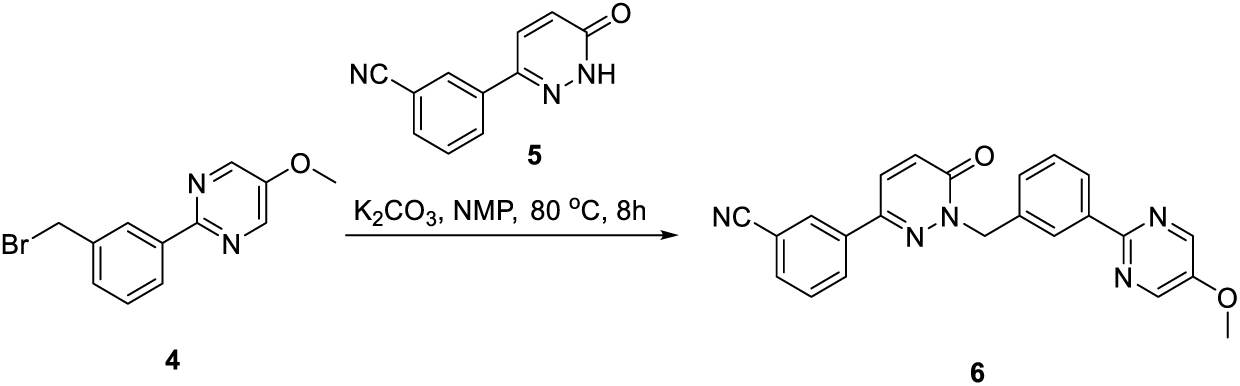

To a stirred solution of **compound 4** (30 g, 107.47 mmol) and K2CO3 (16.3 g, 118.22 mmol) in NMP (250 mL) was added **compound 5** (16.9 g, 85.98 mmol). The mixture was stirred at 80 °C for 16 hours. LCMS showed the reaction was completed. The reaction was quenched by water (1000 mL). The mixture was filtered. The wet filter cake was recrystallized from EtOAc (300 mL) to give **compound 6** as an off-white solid (26 g, crude). LCMS: m/z = 396.0 [M+H] ^+^, Rt = 1.627 min, purity: 88.03% (214 nm)

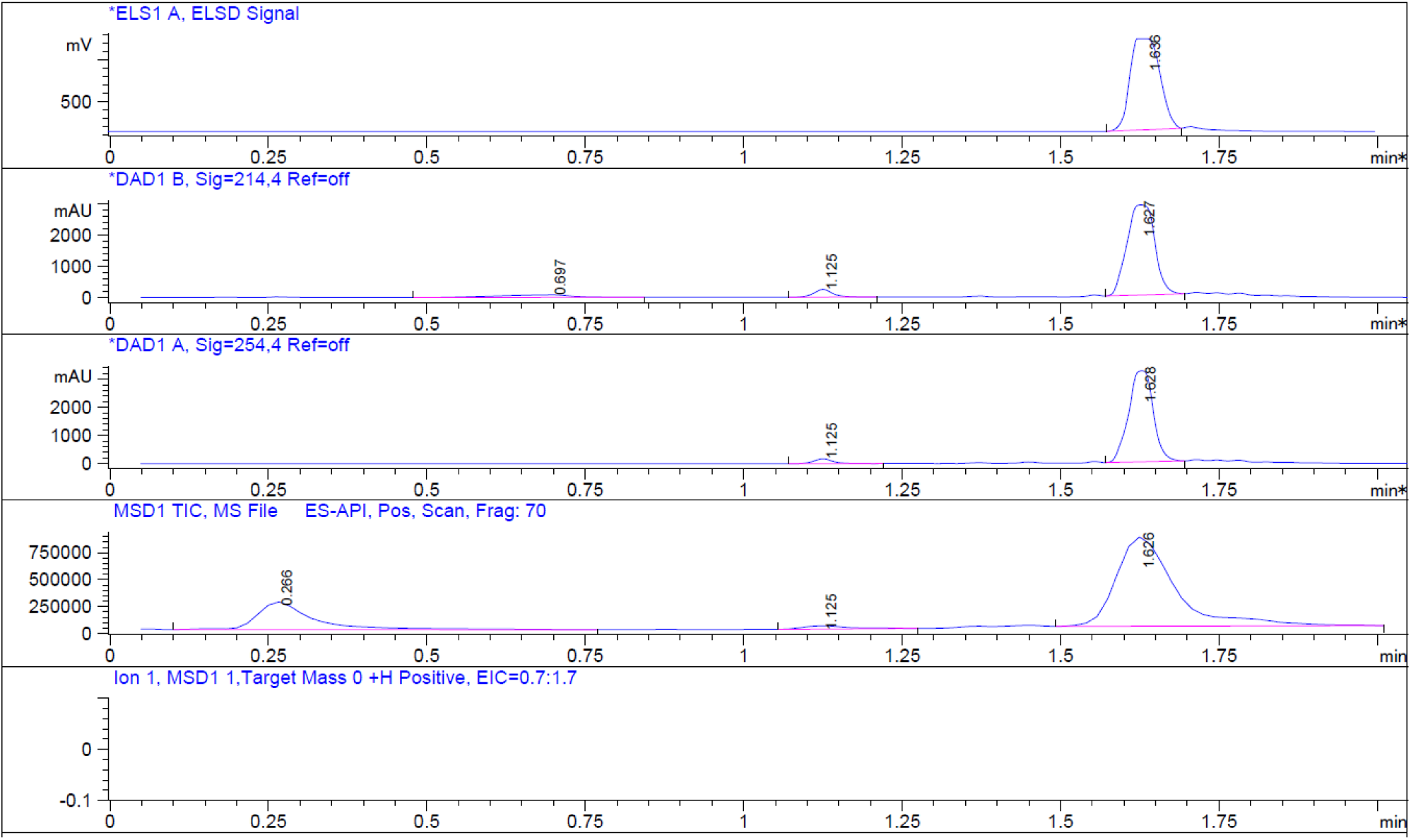

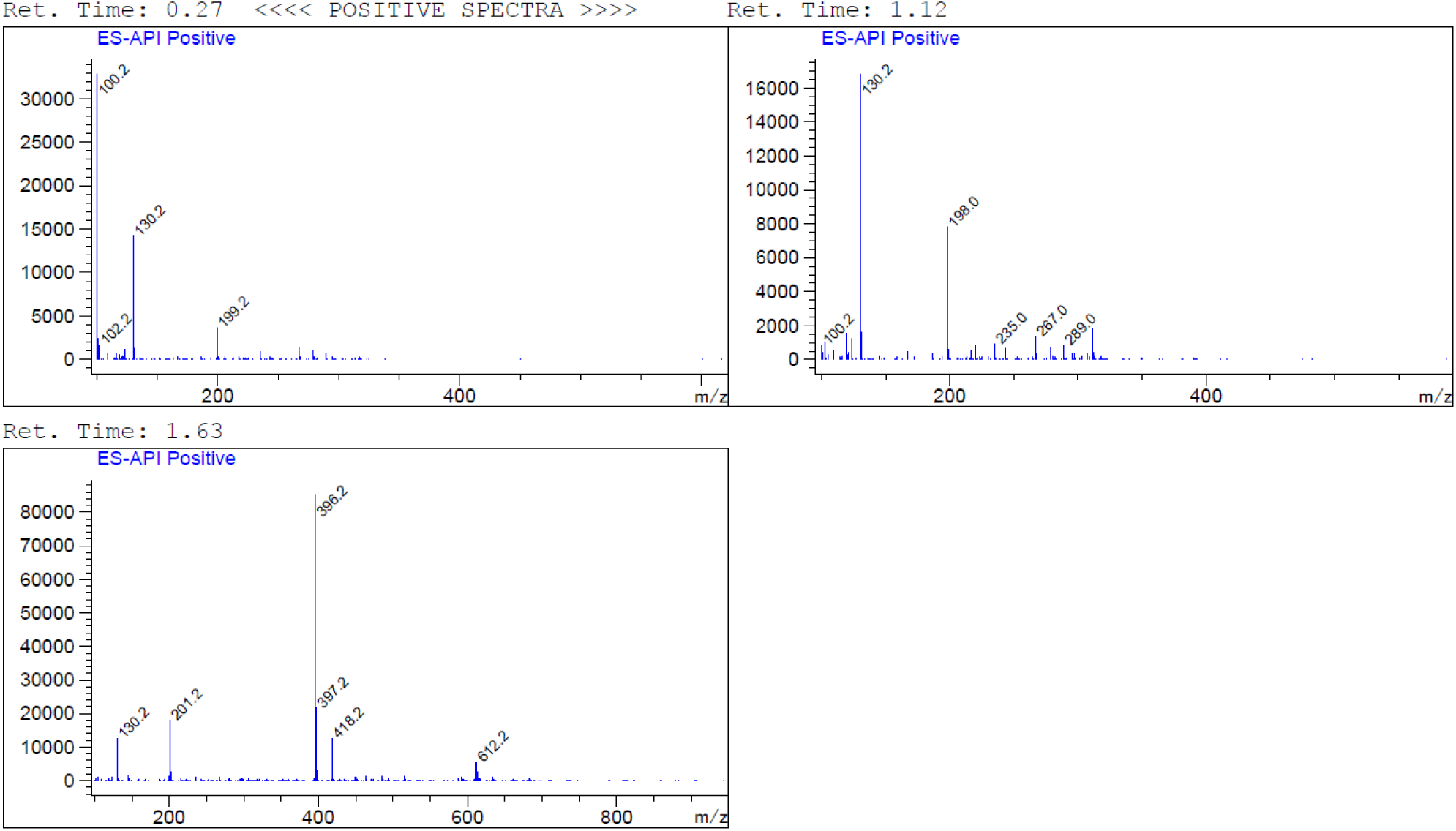

#### (4) Synthesis of compound 7

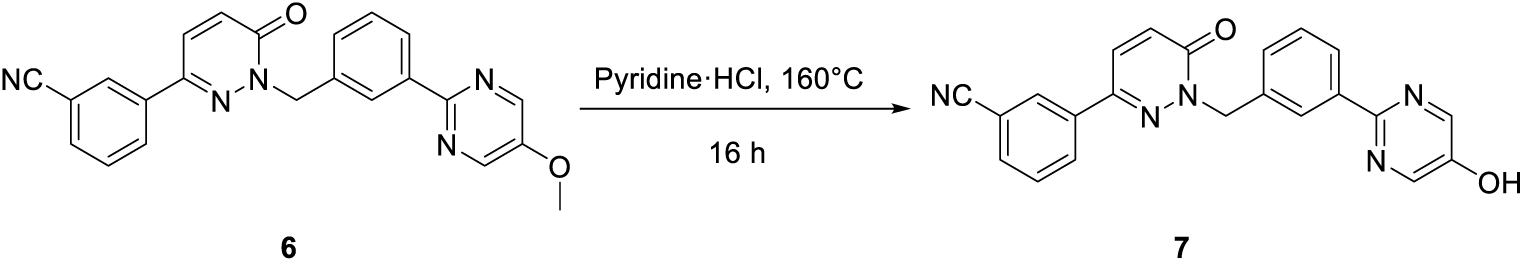

**Compound 6** (26 g, 65.25 mmol) was added pyridine ·HCl. The mixture was stirred at 160 °C for 16 hours. LCMS showed the reaction was completed. The reaction was quenched by water (200 mL). The mixture was filtered. The wet filter cake was recrystallized from EtOAc (200 mL) to give **compound 7** as an off-white solid (31.6 g, crude).

LCMS: m/z = 382.2 [M+H] ^+^, Rt = 1.486 min, purity: 63.38% (214 nm)

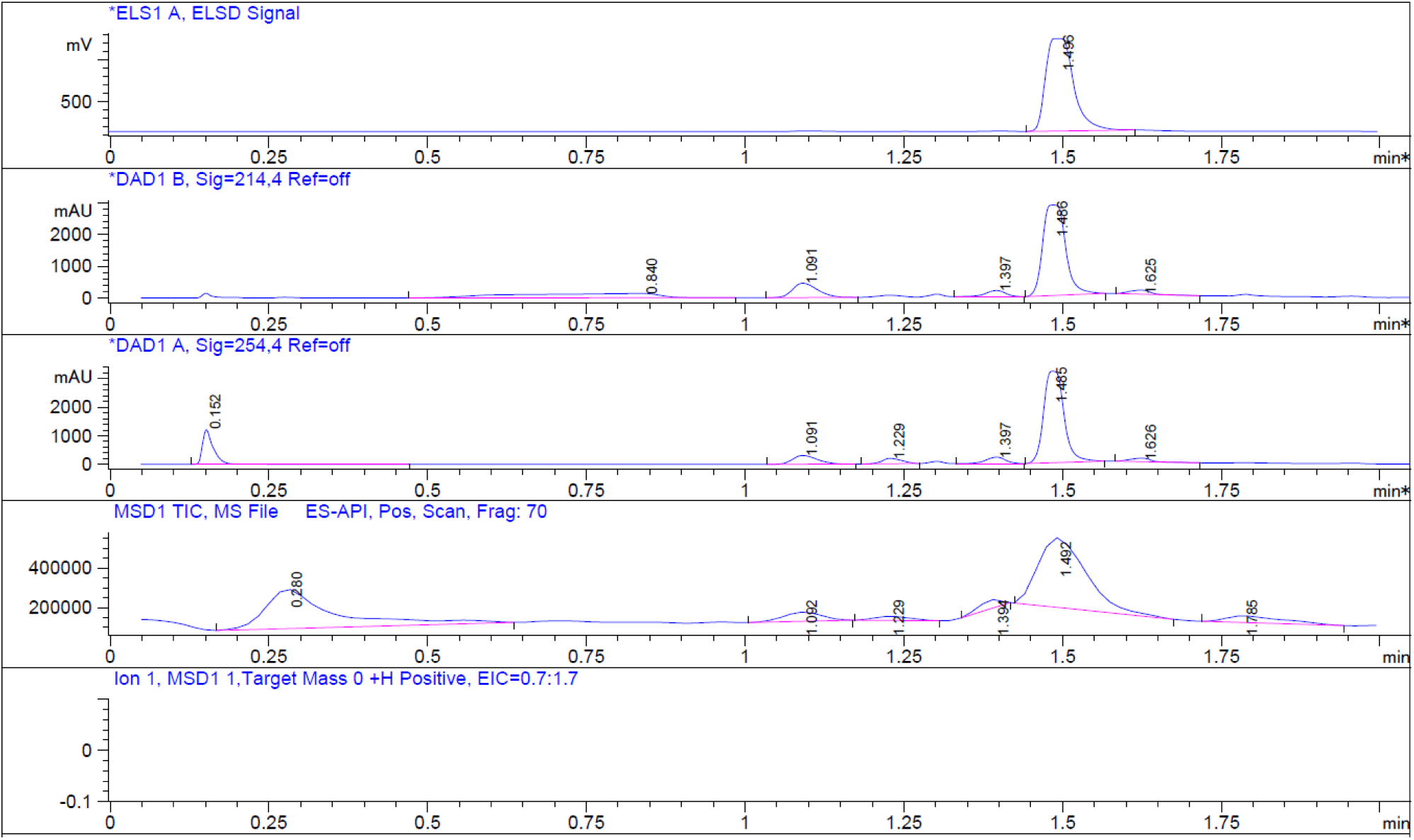

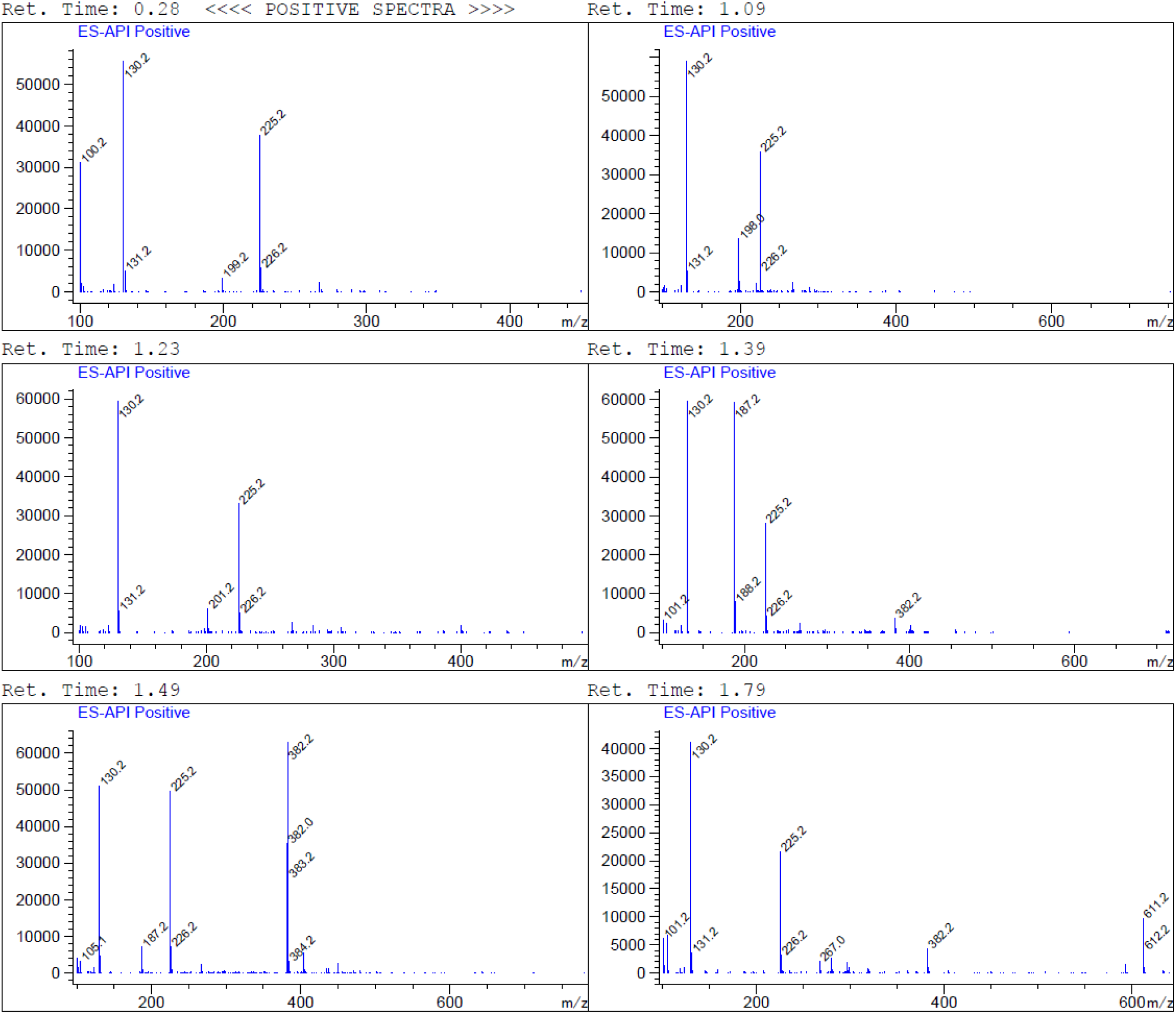

#### (5) Synthesis of compound 9

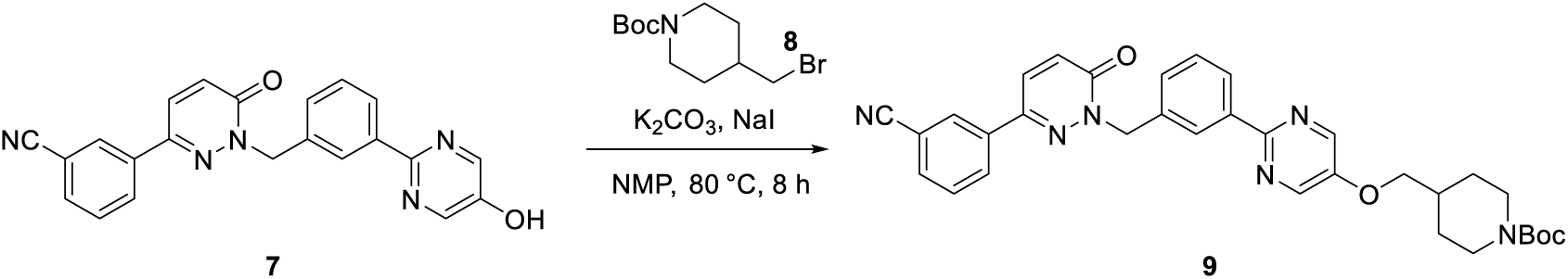

To a suspension of **compound 7** (16.6 g, 43.52 mmol) in NMP (200 mL) were added NaI (0.65 g, 4.35 mmol), K2CO3 (11.75 g, 85.04 mmol) and compound 8 (16.95 g, 60.93 mmol). The reaction was stirred at 80 °C for 8 hours. LCMS showed the reaction was complete. The reaction was quenched by water (800 mL). The mixture was filtered. The wet filter cake was recrystallized from EtOAc (200 mL) to give **compound 9** as an off-white solid (11 g, crude).

LCMS: m/z = 523.2 [M-55] ^+^, Rt = 1.885 min, purity: 91.46% (214nm)

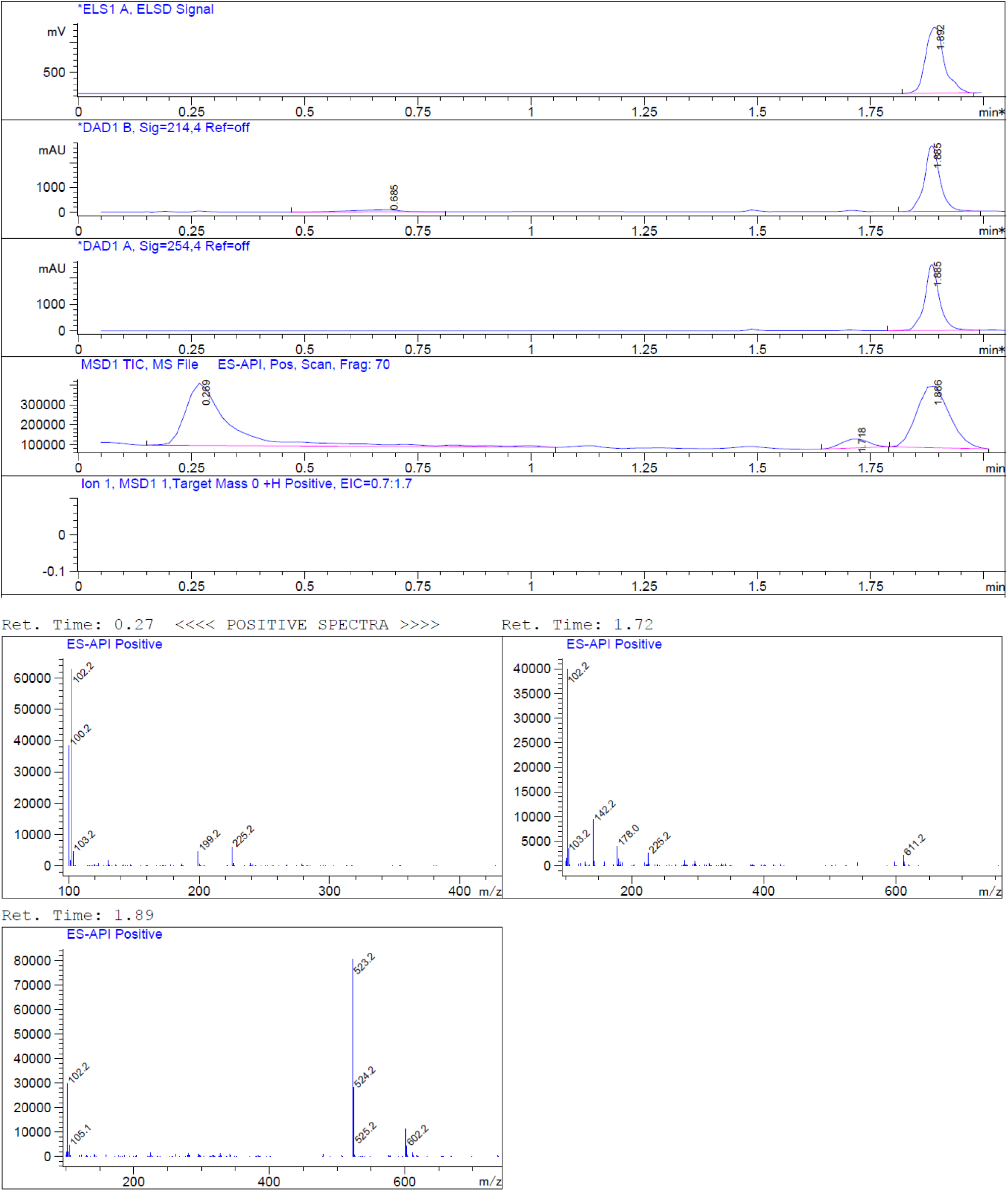

#### (6) Synthesis of compound 10

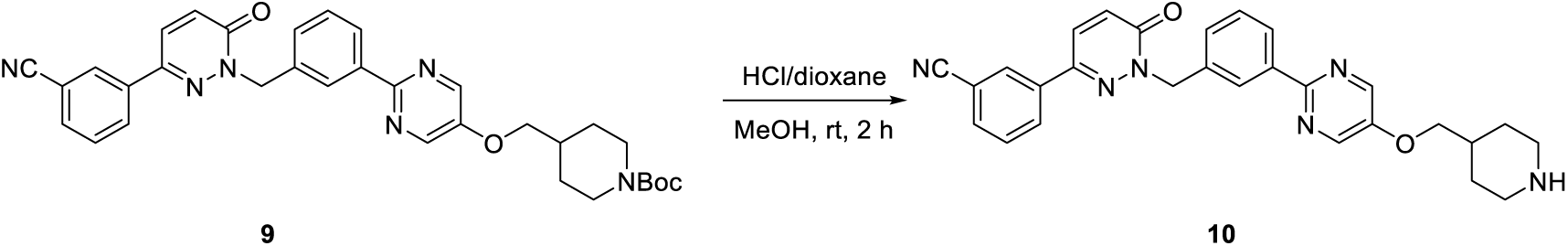

To a solution of **compound 9** (4 g, 6.91 mmol) in DCM (40 mL) was added dropwise TFA (10 mL). The resulting solution was stirred at room temperature for 2 hours. LCMS showed the reaction was complete. The mixture was concentrated in vacuum to give the crude **compound 10** (4 g, crude), which was used for the next step without further purification.

LCMS: m/z = 479.2 [M+H] ^+^, Rt = 1.373 min, purity: 52.07% (214nm)

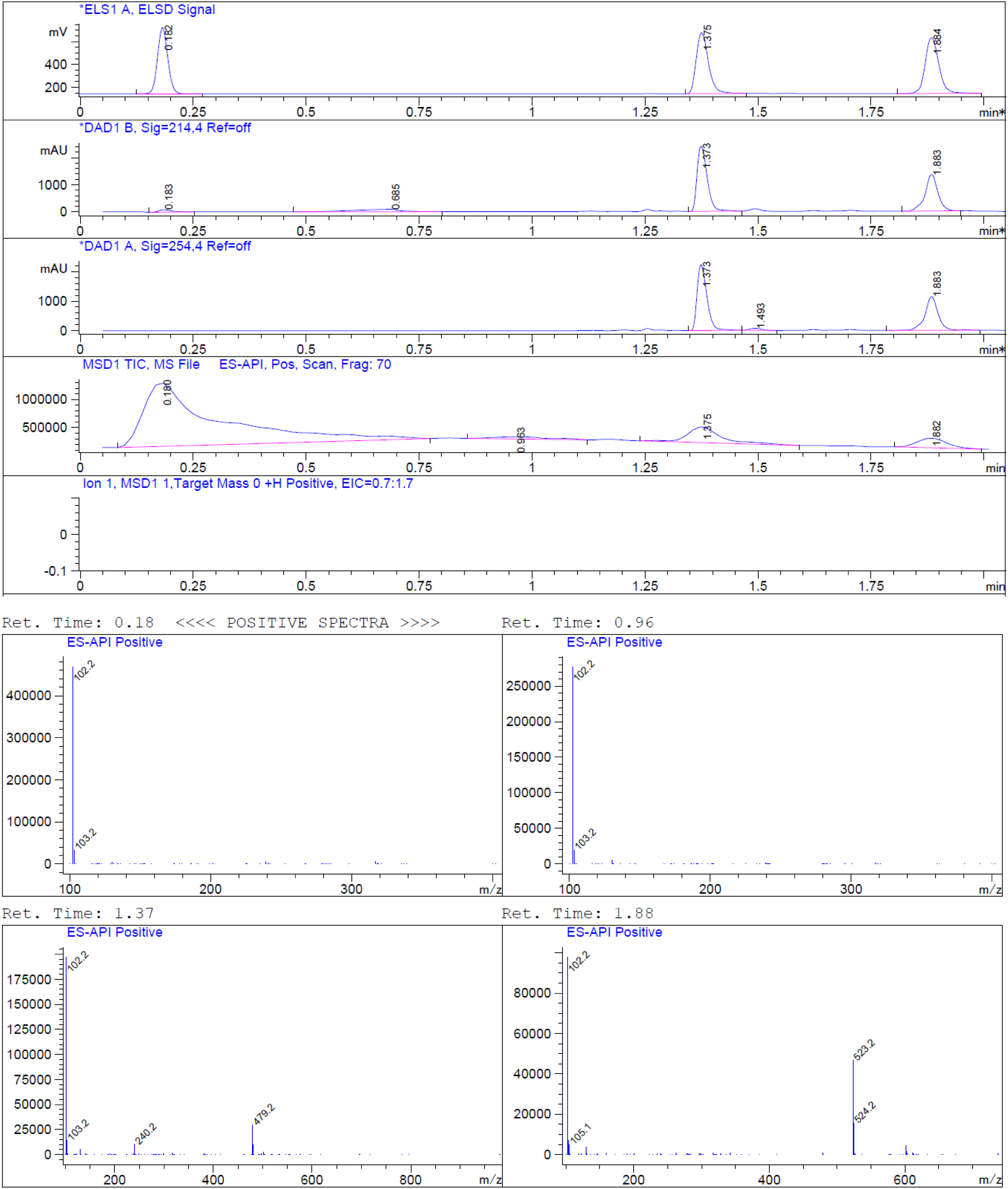

#### (7) Synthesis of compound 12

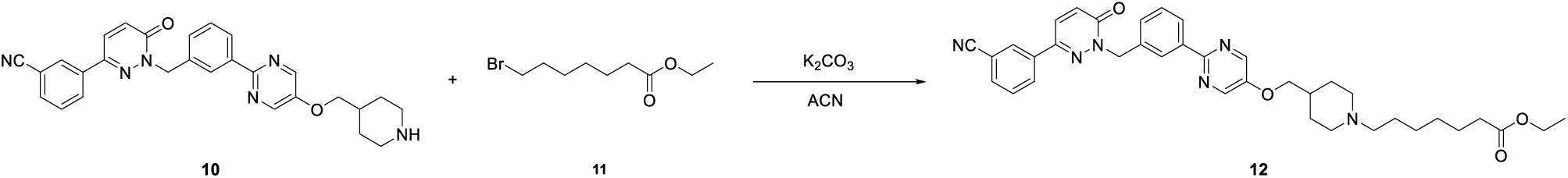

To a solution of **compound 10** (2.5 g, 5.2 mmol) and **compound 11** (1.85 g, 7.84 mmol) in ACN (25 mL) was added K2CO3 (2.16 g, 15.68 mmol) and KI (433 mg, 2.6 mmol) under Ar atmosphere at room temperature. The mixture was stirred under Ar atmosphere at 60 °C 16 hours. LCMS showed the reaction was complete. The mixture was quenched with Water (40 mL) and extracted with EtOAc (30 mL x 3). The organic layers were washed with brine (30 mL), dried over anhydrous Na2SO4 and filtered. The filtrate was concentrated invacuum. The reside was purified by flash column chromatograph (eluted with MeOH: DCM =7%) to afford **compound 12** (2.1 g, 64% yield) as a white solid.

LCMS: m/z = 635.6 [M+H] ^+^, Rt = 1.524 min, Purity: 97.47% (214 nm).

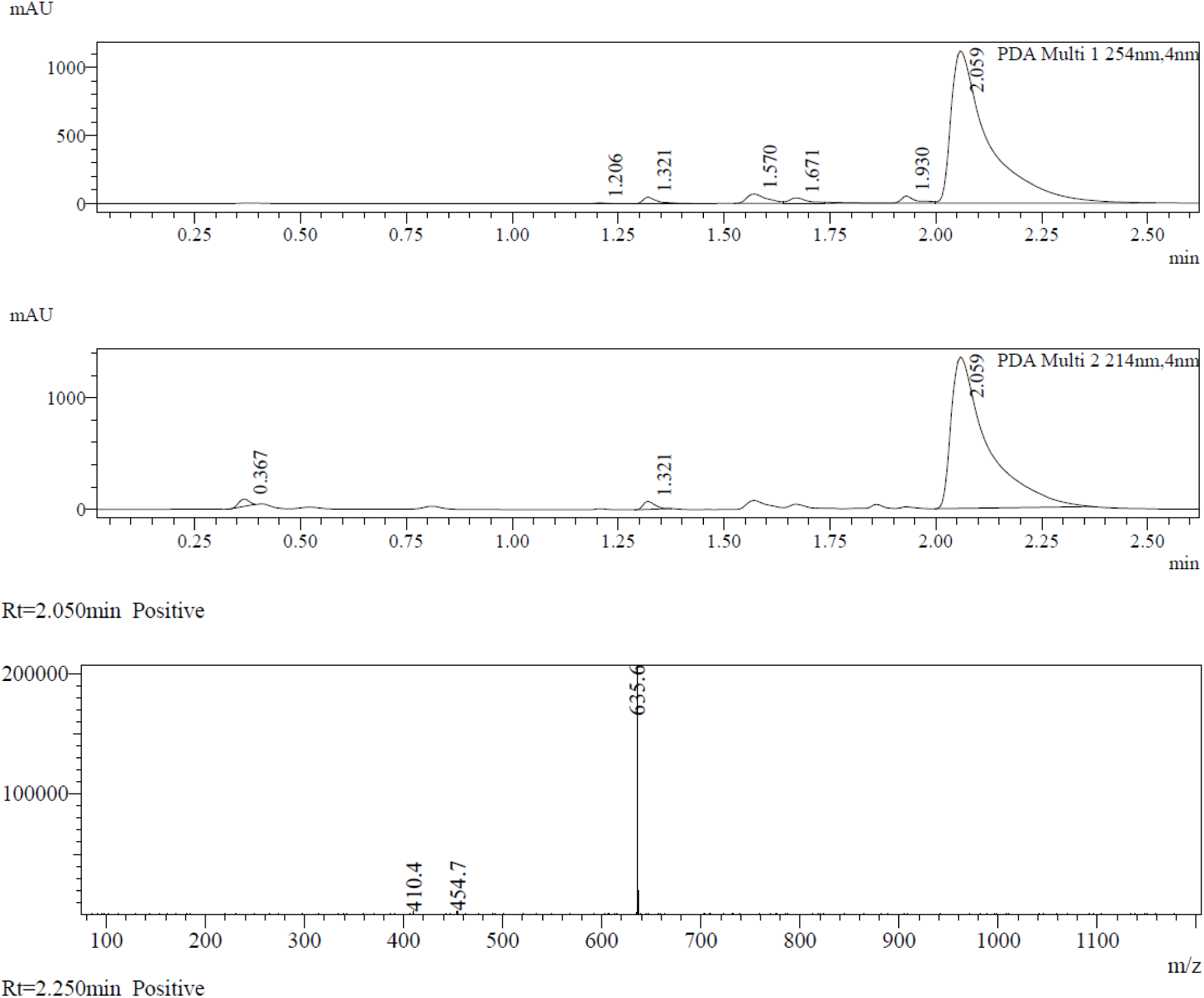

#### (8) Synthesis of compound 13

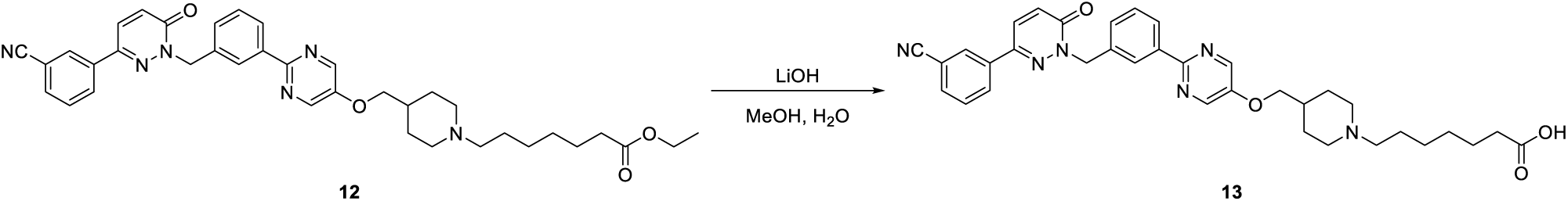

To a solution of compound 12 (1.9 g, 3.0 mmol) in MeOH (20 mL) and H2O (20 mL) was added LiOH (720 mg, 30 mmol). The mixtures were stirred at 35 °C for 2 hours. LCMS showed the reaction was complete. The mixture was concentrated in vacuum to remove of MeOH. The residue was adjusted to pH = 6 with HCl (1M). The mixture was filtered to give the compound 13 (1.5 mg(crude)) as a yellow solid.

LCMS: m/z = 607.2 [M+H] ^+^, Rt = 1.08 min, purity: 69.30% (214 nm)

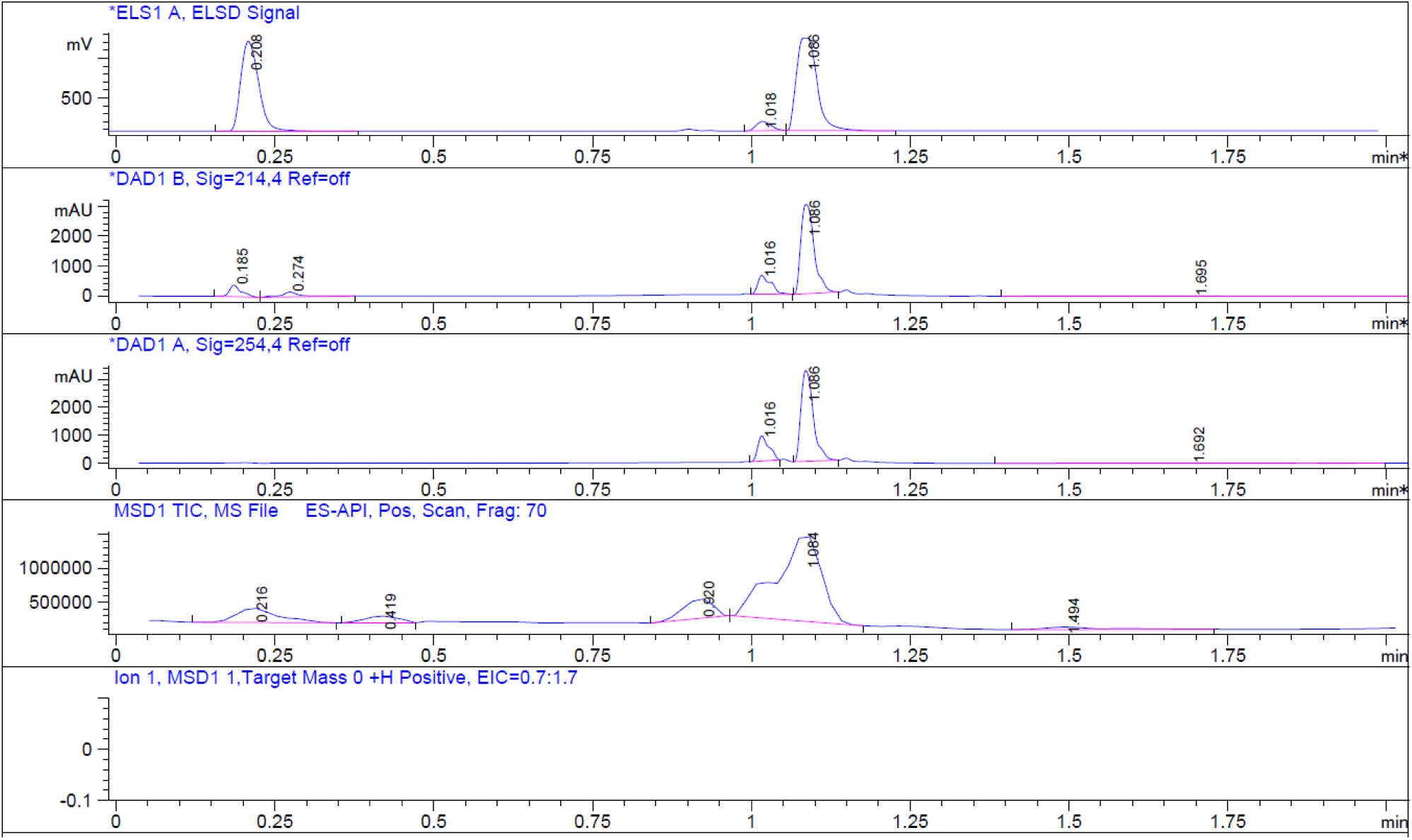

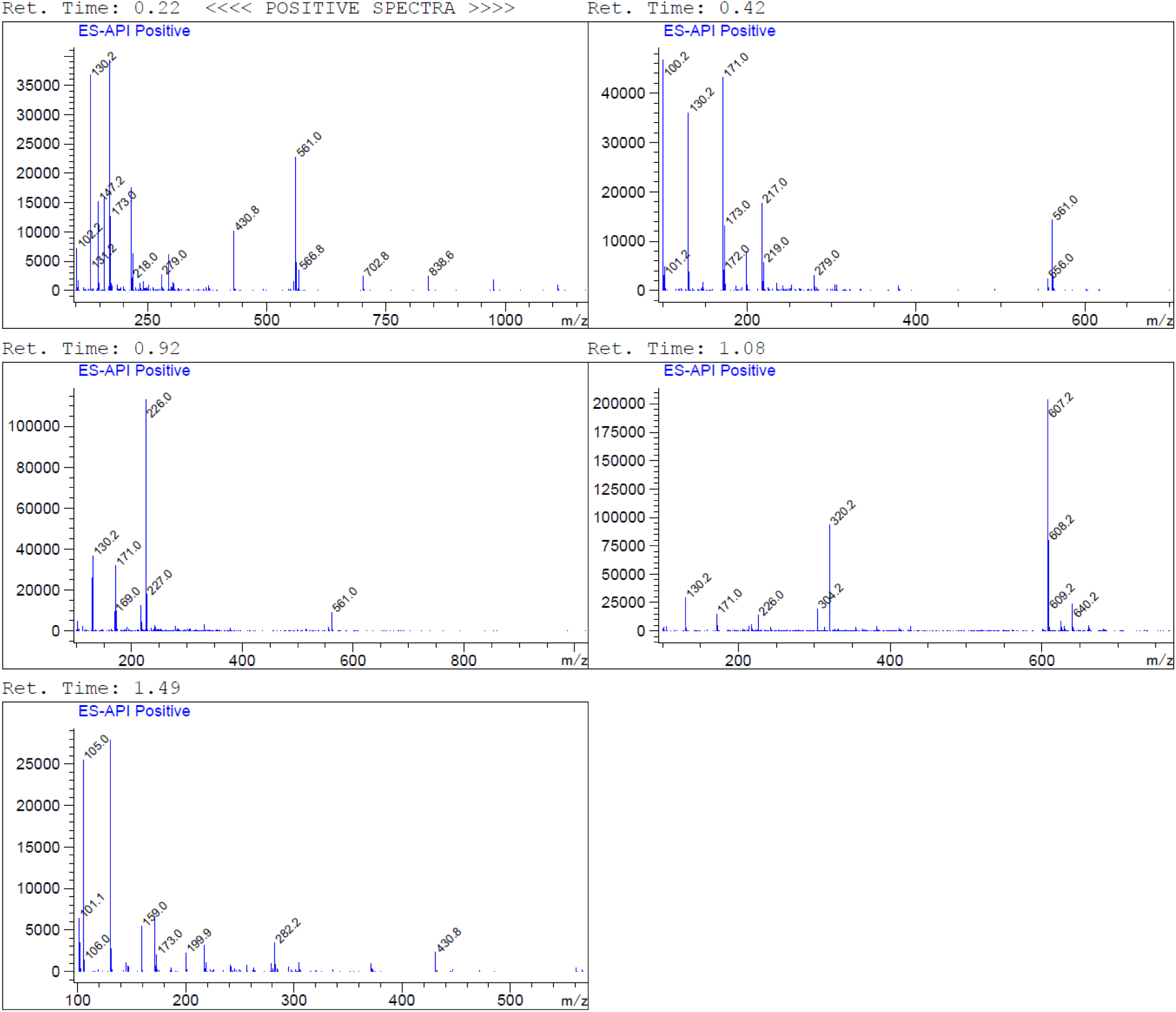

#### (9) Synthesis of compound 15

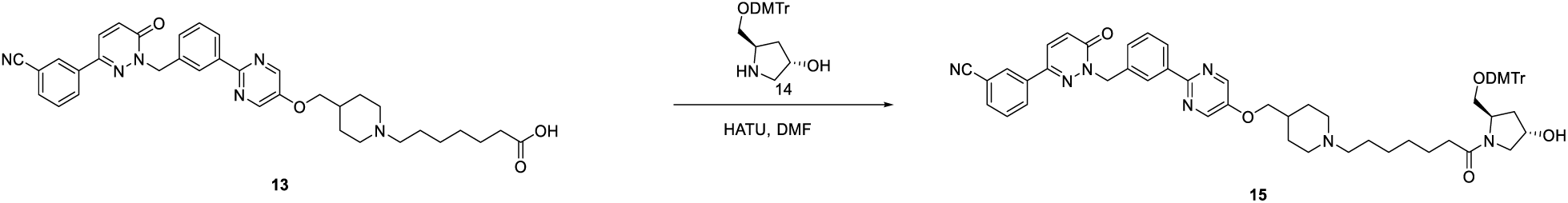

To a solution of **compound 13** (1.5 g, 2.48 mmol) and **compound 14** (1.04 g, 2.5 mmol) in DMF (20 mL) was added HATU (1.9 g, 4.96 mmol) and DIPEA (640 mg, 4.96 mmol). The mixture was stirred under Ar atmosphere at room temperature for 2 hours. LCMS showed the reaction was complete. The mixture was diluted with H2O (30 mL) and extracted with EtOAc (40 mL x 3). The organic layers were wash with brine (30mL), dried over anhydrous Na2SO4 and filtered. The filtrate was concentrated in vacuum. The residue was purified by flash column chromatograph (eluted with MeOH: DCM = 10 %) to give **compound 15** (1.7 g, 68 % yield) as a white solid

LCMS: m/z = 1008.4 [M+H] ^+^, Rt = 1.700 min, purity: 92.5% (214 nm)

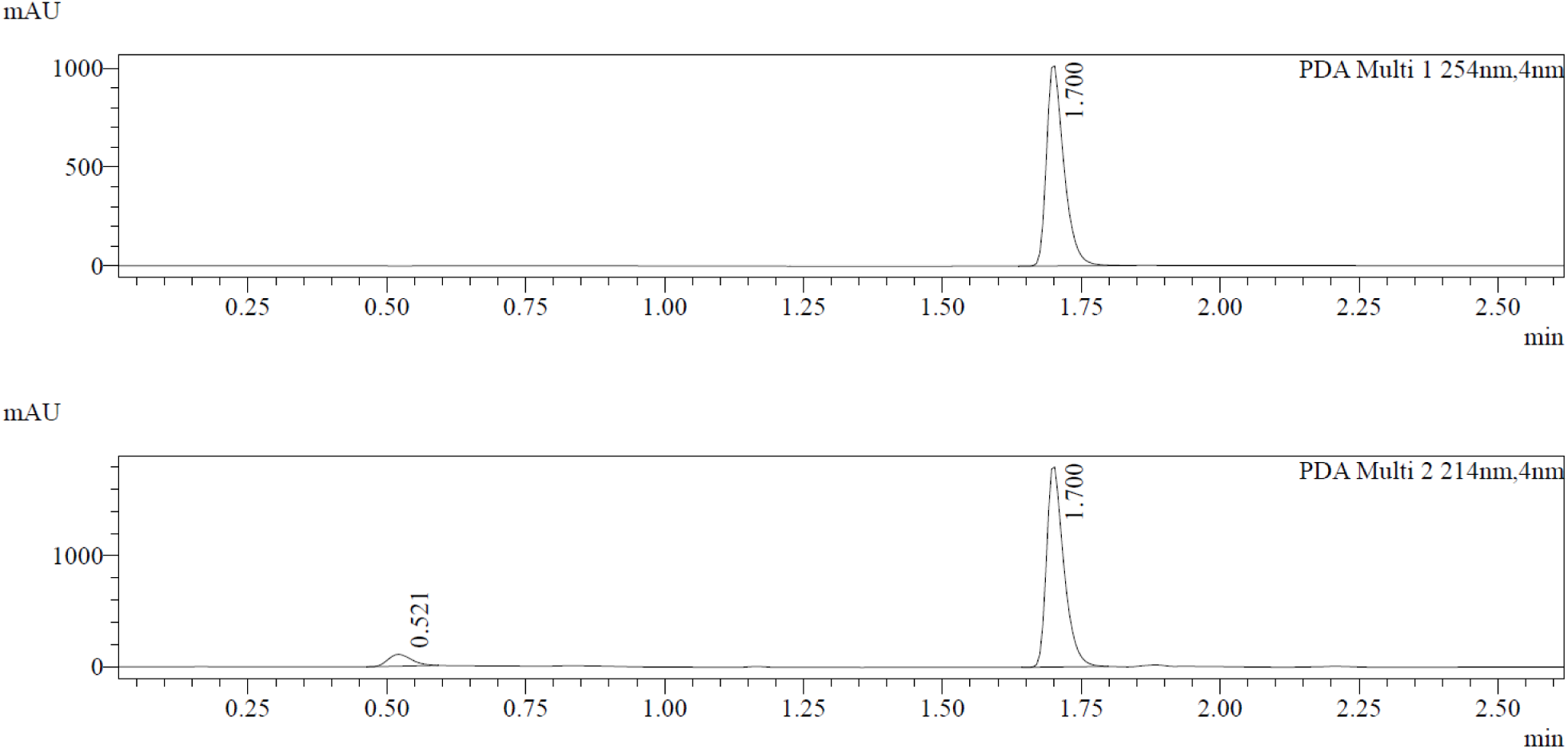

#### (10) Synthesis of compound 17

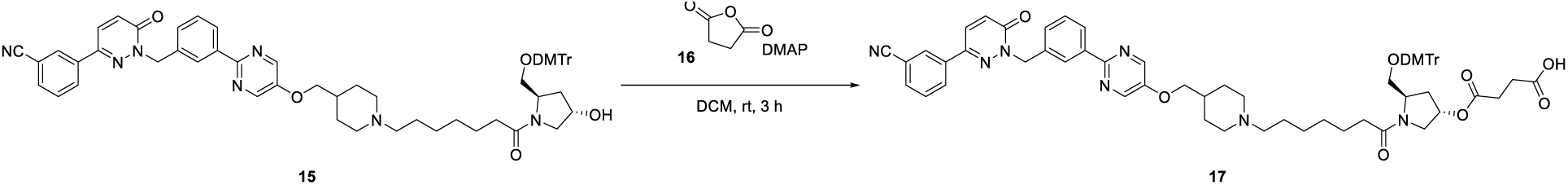

To a solution of **compound 15** (1.7 g, 1.69 mmol) in DCM (30 mL) were added **compound 16** (337.5 mg, 3.38 mmol) and DMAP (364.9 mg, 3.18 mmol) under nitrogen protected. The mixture was stirred at room temperature for 3 hours. LCMS showed the reaction was complete. The mixture was concentrated in vacuum. The residue was purified by reversed phase column chromatography to give **compound 17** (1.23 g, 65.78 % yield) as a white solid.

LCMS: m/z = 1108.4 [M+H] ^+^, Rt= 1.843 min, purity:100% (214 nm)

**^1^H NMR (400 MHz, DMSO-*d_6_*)** *δ* 8.64 (s, 2H), 8.42 - 8.31 (m, 2H), 8.28 - 8.13 (m, 3H), 7.99 - 7.88 (m, 1H), 7.77 - 7.65 (m, 1H), 7.54 - 7.43 (m, 2H), 7.37 - 7.26 (m, 4H), 7.25 - 7.11 (m, 6H), 6.96 - 6.79 (m, 4H), 5.44 (s, 2H), 5.41 - 5.18 (m, 1H), 4.27 - 4.13 (m, 1H), 4.09 - 4.00 (m, 2H), 3.80 - 3.75 (m, 1H), 3.73 (s, 6H), 3.59- 3.54 (m, 2H), 3.26 - 3.19 (m, 2H), 3.09 - 2.95 (m, 4H), 2.49 - 2.44 (m, 4H), 2.44 - 1.89 (m, 9H), 1.88 - 1.70 (m, 3H), 1.55 - 1.21 (m, 9H), 1.21 - 1.05 (m, 2H).

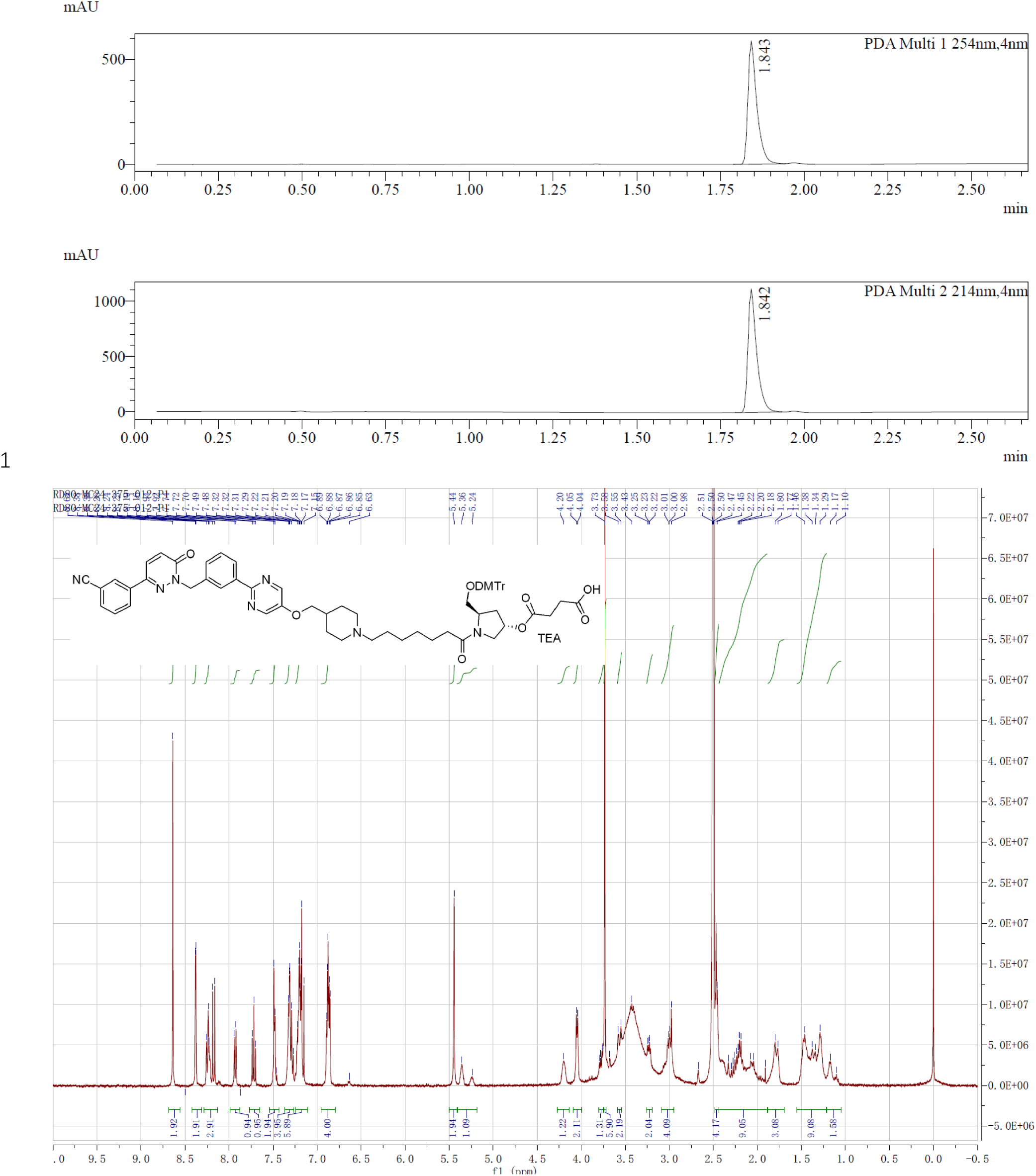

#### (11) Synthesis of SAPT8-Tep-2-CPG

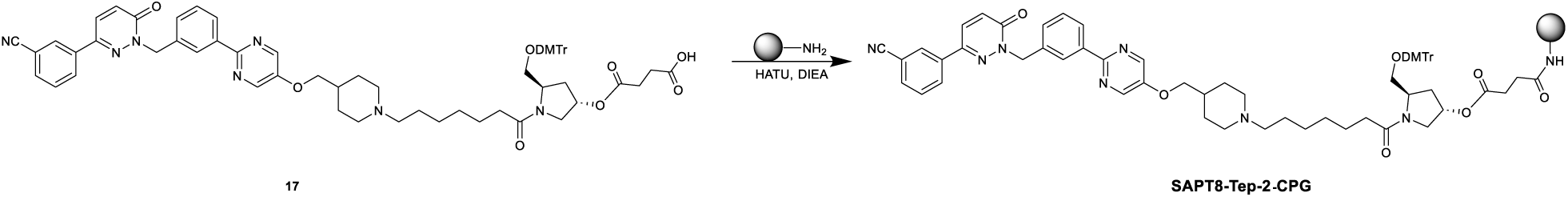

To a solution of **compound 17** (300 mg, 248.0 μmol) in ACN (18.0 mL) were added HATU (120 mg, 315.6 μmol), DIEA (120 μL), lcaa-CPG (1000 Å, 1500 mg) respectively at room temperature and oscillates for 12 hours. After the reaction was completed, CPG was washed with ACN, CAP A (acetic anhydride: tetrahydrofuran= 1:9, v/v, 12.0 mL) and CAP B (N-methylimidazole: pyridine: acetonitrile= 15:10:75, v/v/v, 12.0 mL) was added, the mixture was oscillated at room temperature for 1 hour. Then the mixture was filtered and washed with ACN (180 mL) for 3 times. **SAPT8-Tep-2-CPG** was obtained after freeze-drying of the mixture as a white powder (1500 mg).

#### (12) Synthesis of SAPT8-Tep-2

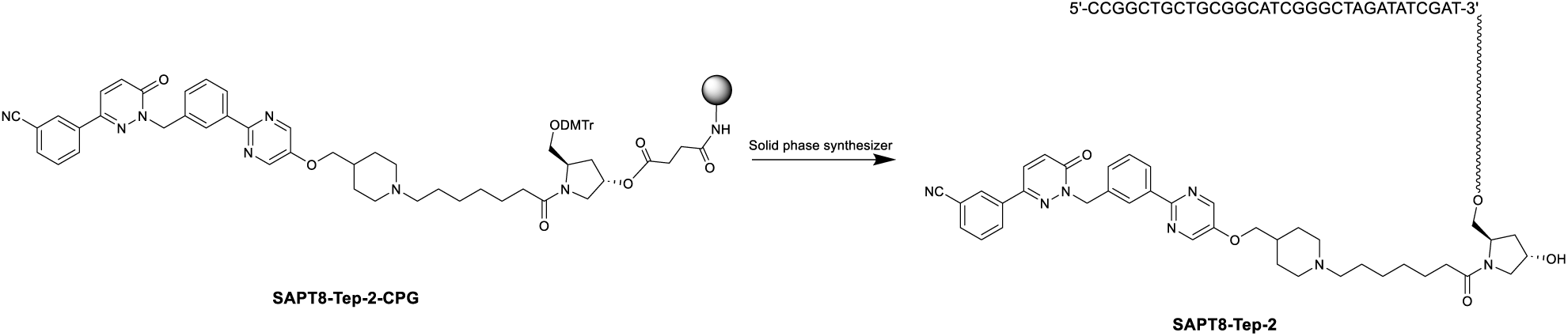

**SAPT8-Tep-2-CPG** in synthesis columns (60 mg*24) and synthesize by K-A H-8 Solid phase synthesizer. Solid phase synthesis involves four steps: Detritylation, Coupling, Capping and Oxidation. After reaction was done, CPG of each synthesis column was added 1.5 mL aqueous ammonia and heated in oven at 65 °C for 16 hours. Then collected the supernatant and washed with water (1 mL*3). The combined crude was purified by RP-IP-HPLC (WATERS 2489, 3767) (**Column**: XBridge BET C18 2.5μm. **RP-IP-HPLC method**: Mobile phase A: 2 % HFIP +0.1 % TEA in DD water, Mobile phase B: MeOH) to afford **SAPT8-Tep-2** as a white powder (39.84 mg, purity = 99.369%).

UPLC-MS (WATERS ACQUITY PREMIER): **SAPT8-Tep-2-MS**, m/z = 10624.78099 [M]^-^(deconvolution); *t*_R_ = 10.802 min (260 nm). Mass error <50 ppm.

HPLC (SPT-M40): **SAPT8-Tep-2-HPLC**, *t*_R_ = 11.139 min (260 nm), purity: 99.369%.

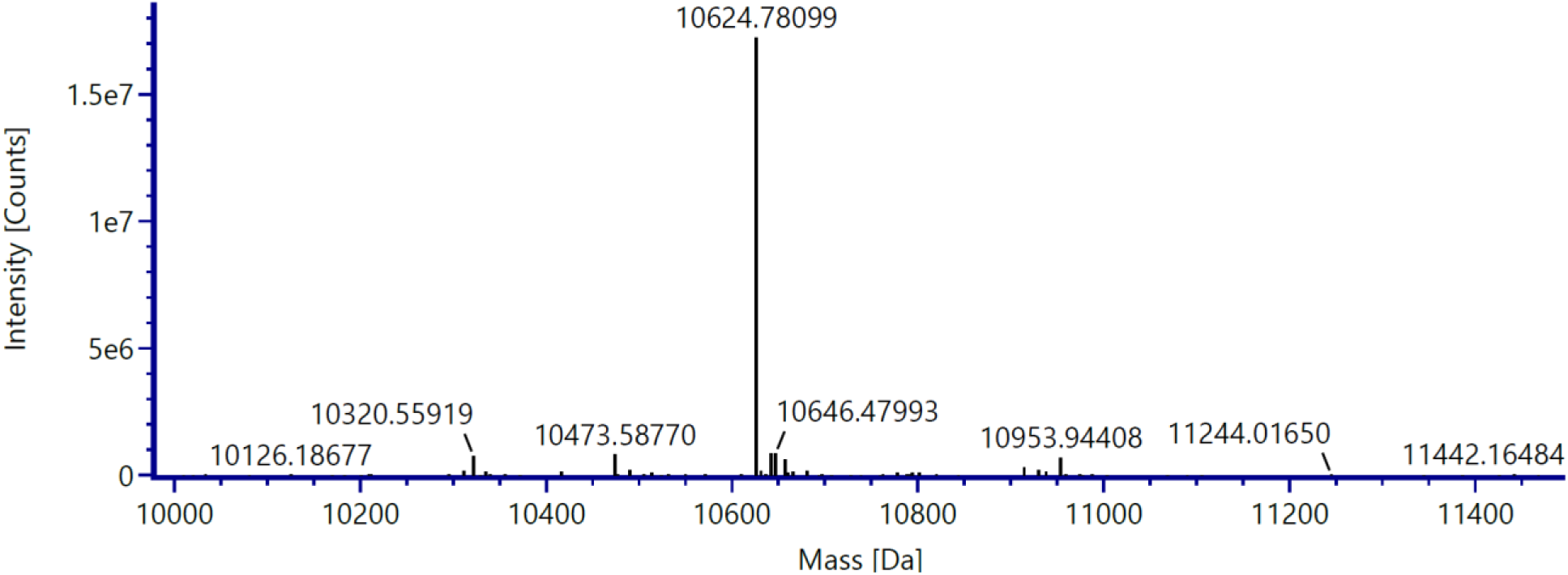

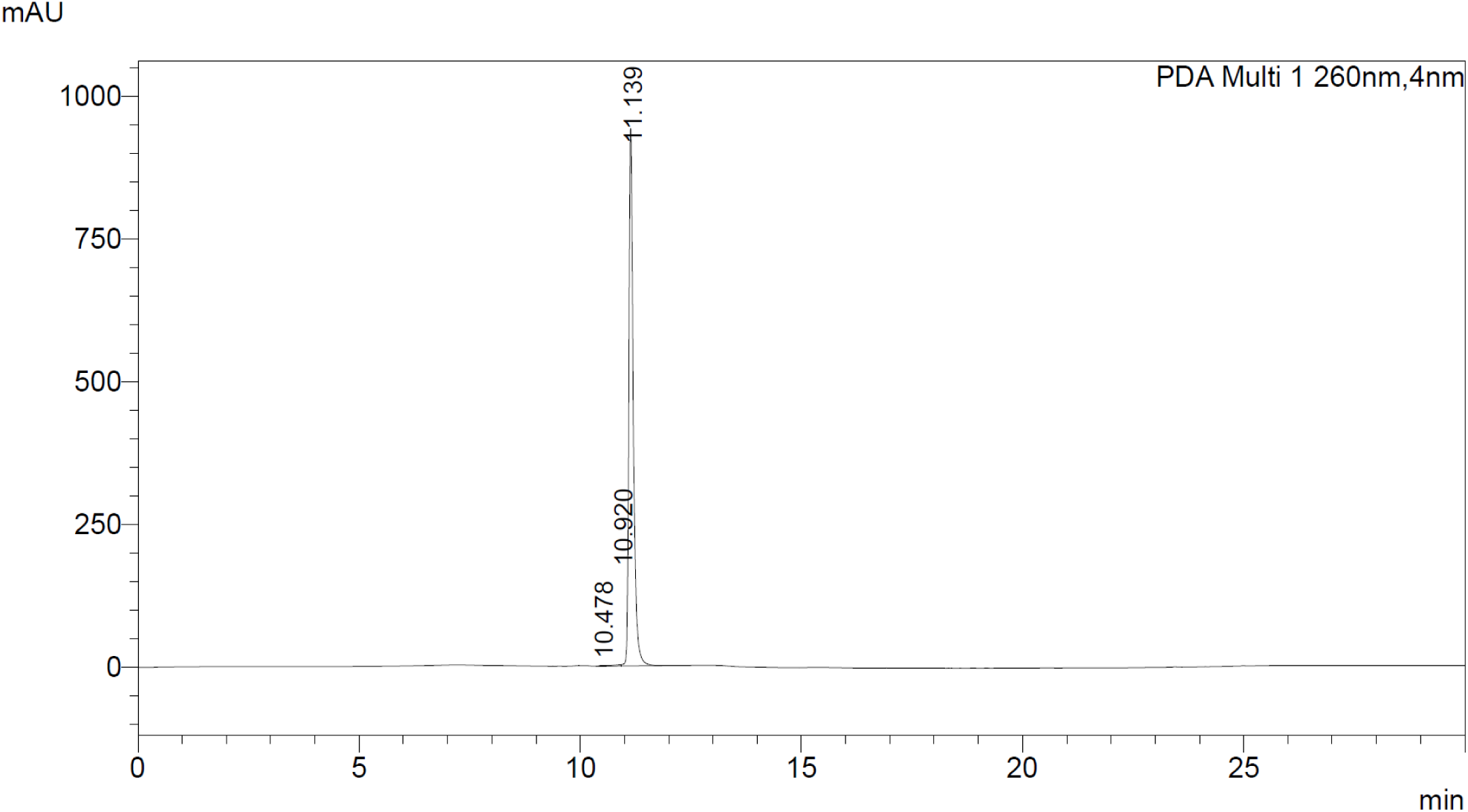

### Scheme 3. Synthetic route for SAPT8-Tep-3

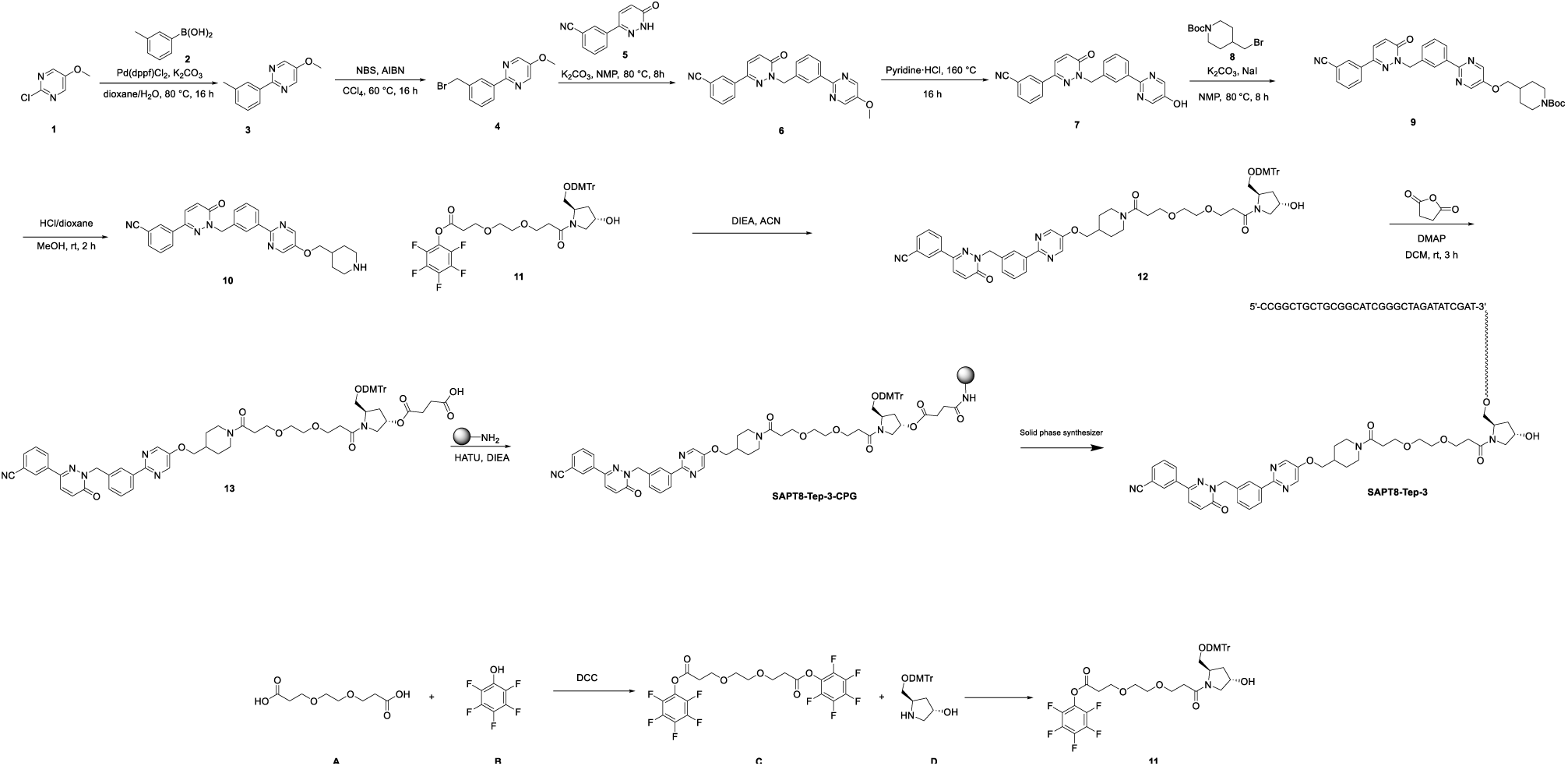

#### (1) Synthesis of compound 3

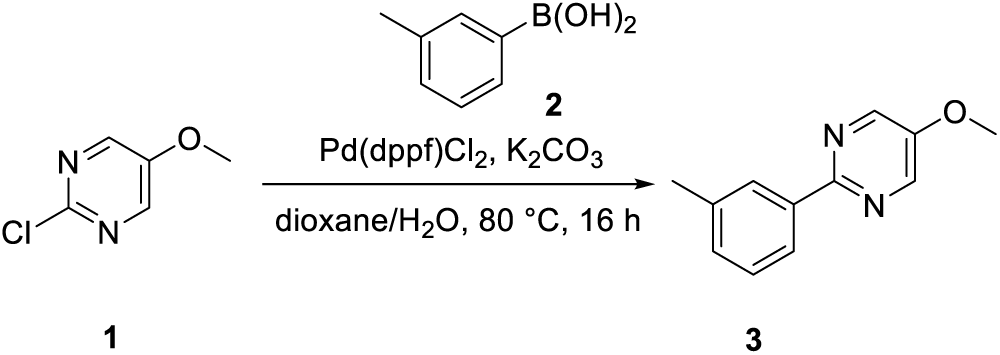

To a solution of **compound 1** (25 g, 172.94 mmol) in dioxane (450 mL) and H2O (150 mL) were added **compound 2** (23.5 g, 172.94 mmol), K2CO3 (47 g, 345.88 mmol) and Pd(dppf)Cl2 (6.2 g, 8.65 mmol). The mixture was stirred at 80 °C under nitrogen atmosphere for 16 hours. LCMS showed the reaction was complete. The mixture was diluted with water (300 mL) and extracted with EtOAc (100 mL x 3). The organic layers were combined and washed with saturated aqueous NaCl (300 mL). The organic phase was dried over anhydrous Na2SO4 and filtered. The filtrate was concentrated in vacuum. The residue was purified by flash column chromatography eluting with EtOAc in PE (0-30%) to afford **compound 3** (32 g, yield: 94%) as a brown solid.

LCMS: m/z = 201.2 [M+H] ^+^, Rt = 2.13 min, purity: 96.34% (214 nm).

**^1^HNMR (400 MHz, DMSO-*d*_6_)** *δ* 8.64 - 8.62 (m, 2H), 8.20 - 8.05 (m, 2H), 7.40 - 7.36 (m, 1H), 7.29 - 7.27 (m, 1H), 3.96 (s, 3H), 2.40 (s, 3H).

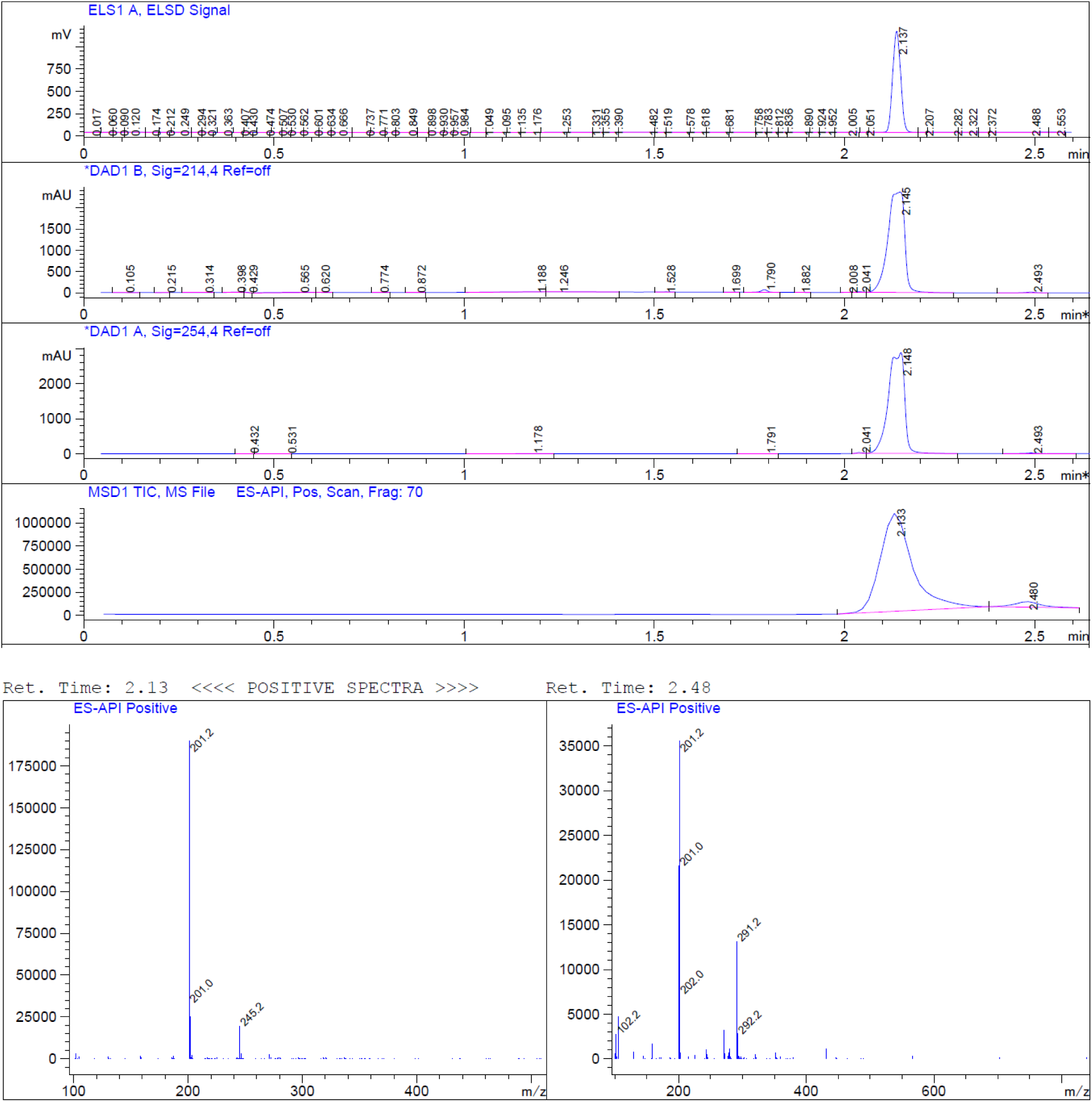

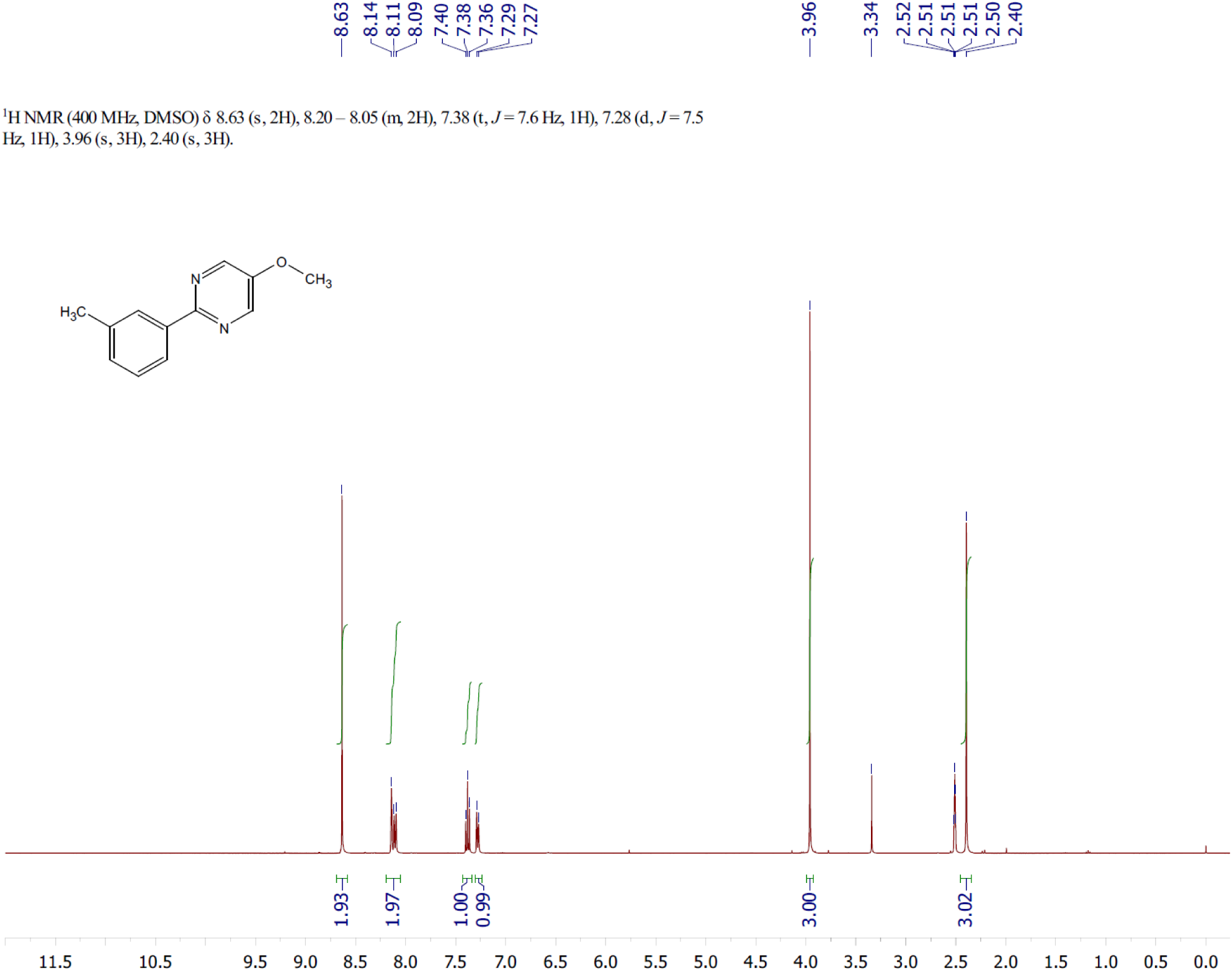

#### (2) Synthesis of compound 4

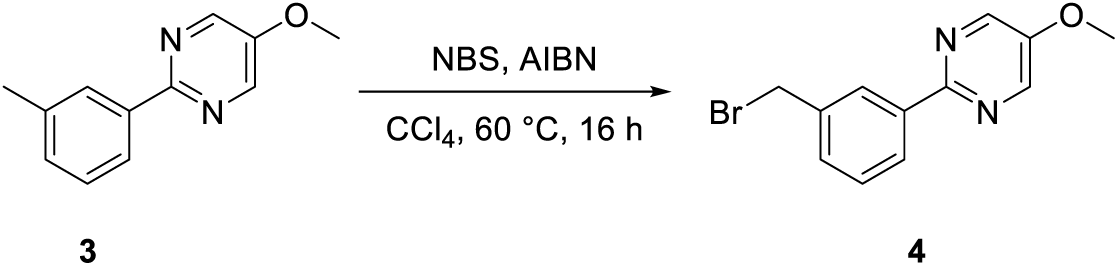

To a solution of **compound 3** (18.7 g, 93.39 mmol) and NBS (16.6 g, 93.39 mmol) in CCl4 (190 mL) were added AIBN (1.5 g, 9.34 mmol). The mixture was stirred at 60 °C under nitrogen atmosphere for 16 hours. LCMS showed the reaction was complete. The mixture was diluted with water (500 mL) and extract with EtOAc (150 mL x 3). The organic phase was washed with saturated aqueous NaCl (500 mL), dried over Na2SO4 and filtered. The filtrate was concentrated in vacuo to afford **compound 4** (crude, 30 g) as a brown solid.

LCMS: m/z = 279.0 [M+H] ^+^, Rt = 1.657min, purity: 72.41% (214 nm).

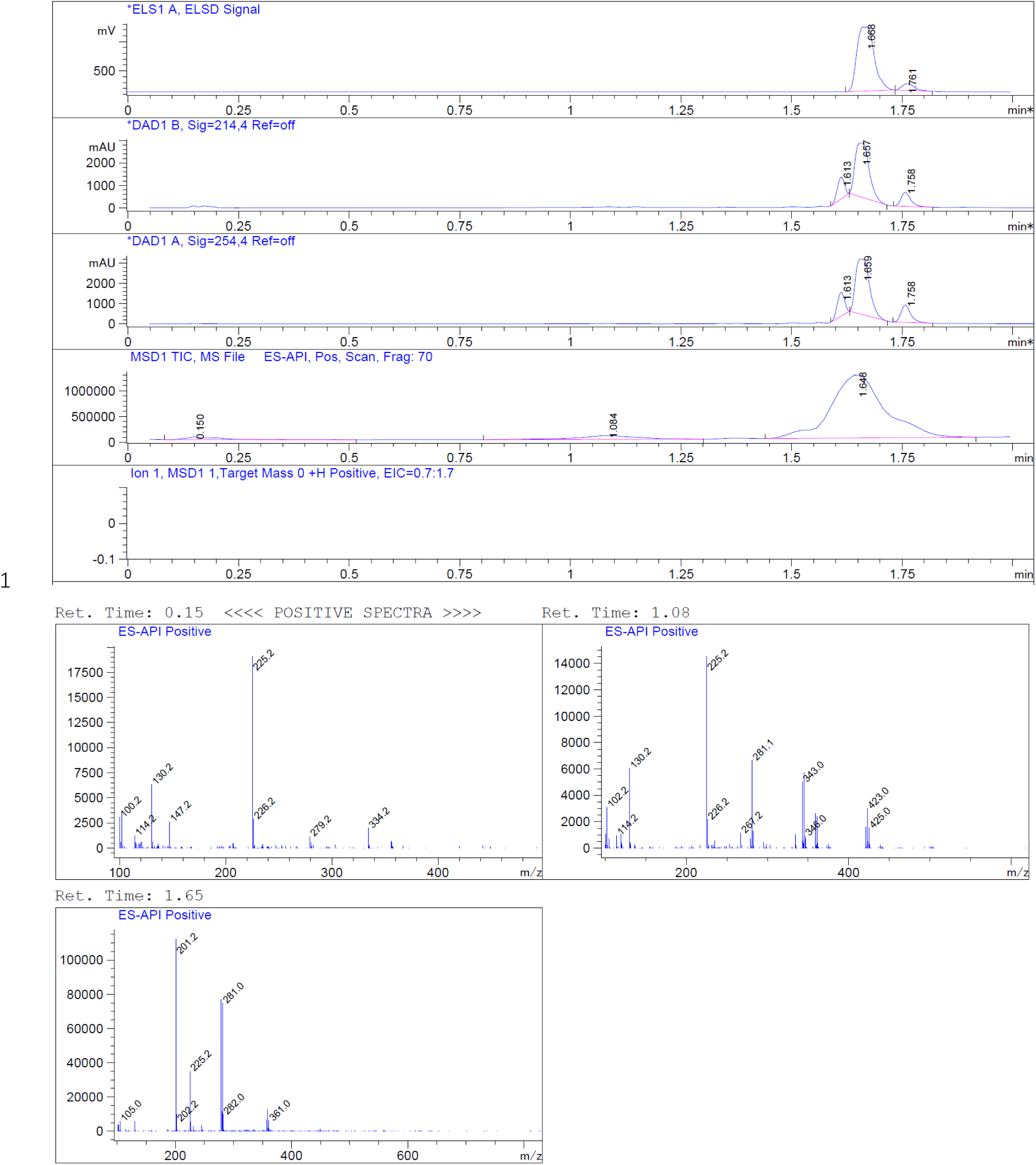

#### (3) Synthesis of compound 6

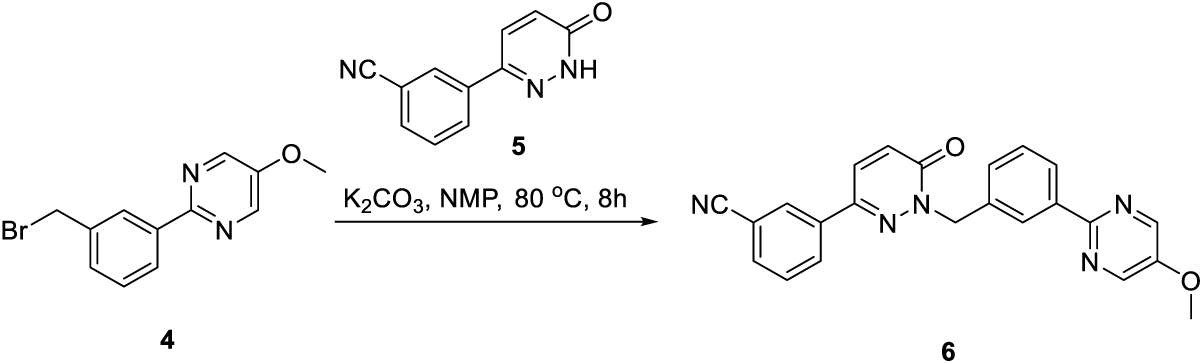

To a stirred solution of **compound 4** (30 g, 107.47 mmol) and K2CO3 (16.3 g, 118.22 mmol) in NMP (250 mL) was added compound 5 (16.9 g, 85.98 mmol). The mixture was stirred at 80 °C for 16 hours. LCMS showed the reaction was completed. The reaction was quenched by water (1000 mL). The mixture was filtered. The wet filter cake was recrystallized from EtOAc (300 mL) to give **compound 6** as an off-white solid (26 g, crude).

LCMS: m/z = 396.0 [M+H] ^+^, Rt = 1.627 min, purity: 88.03% (214 nm)

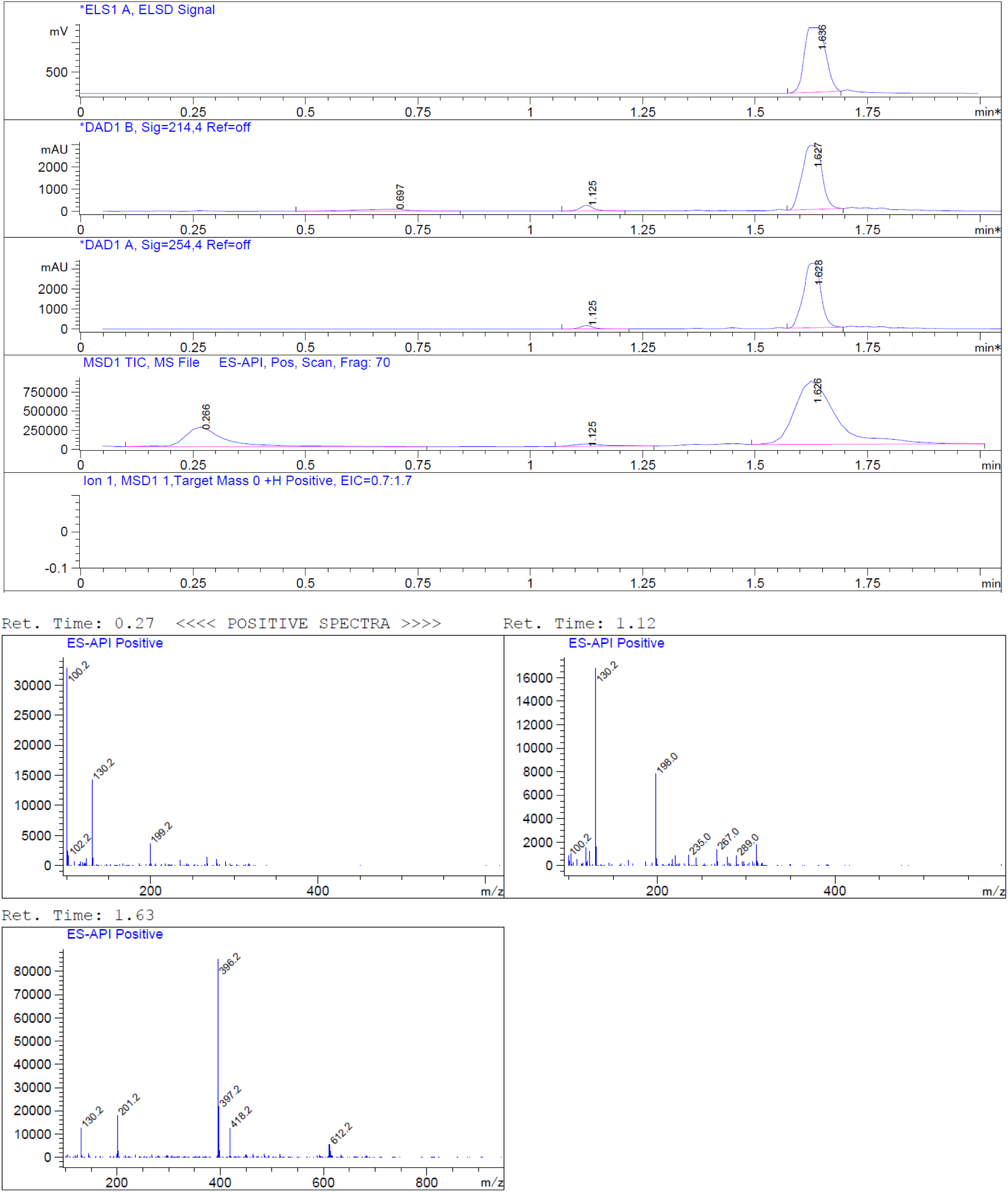

#### (4) Synthesis of compound 7

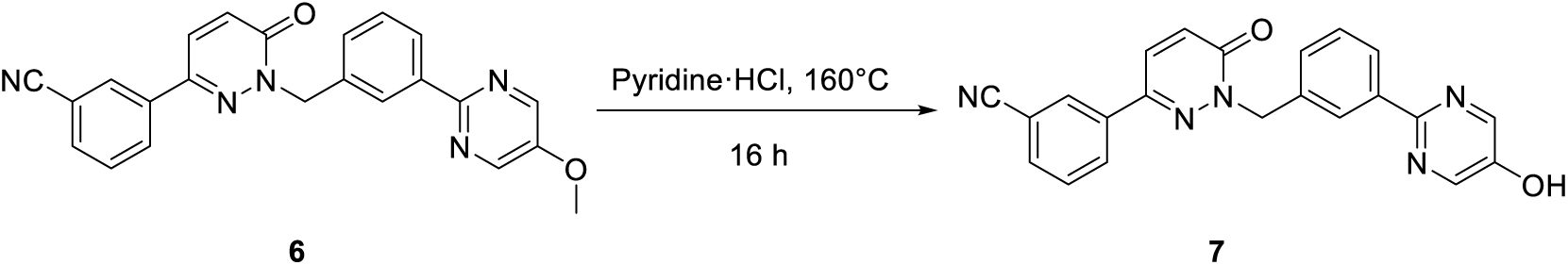

**Compound 6** (26 g, 65.25 mmol) was added pyridine ·HCl. The mixture was stirred at 160 °C for 16 hours. LCMS showed the reaction was completed. The reaction was quenched by water (200 mL). The mixture was filtered. The wet filter cake was recrystallized from EtOAc (200 mL) to give **compound 7** as an off-white solid (31.6 g, crude).

LCMS: m/z = 382.2 [M+H] ^+^, Rt = 1.486 min, purity: 63.38% (214 nm)

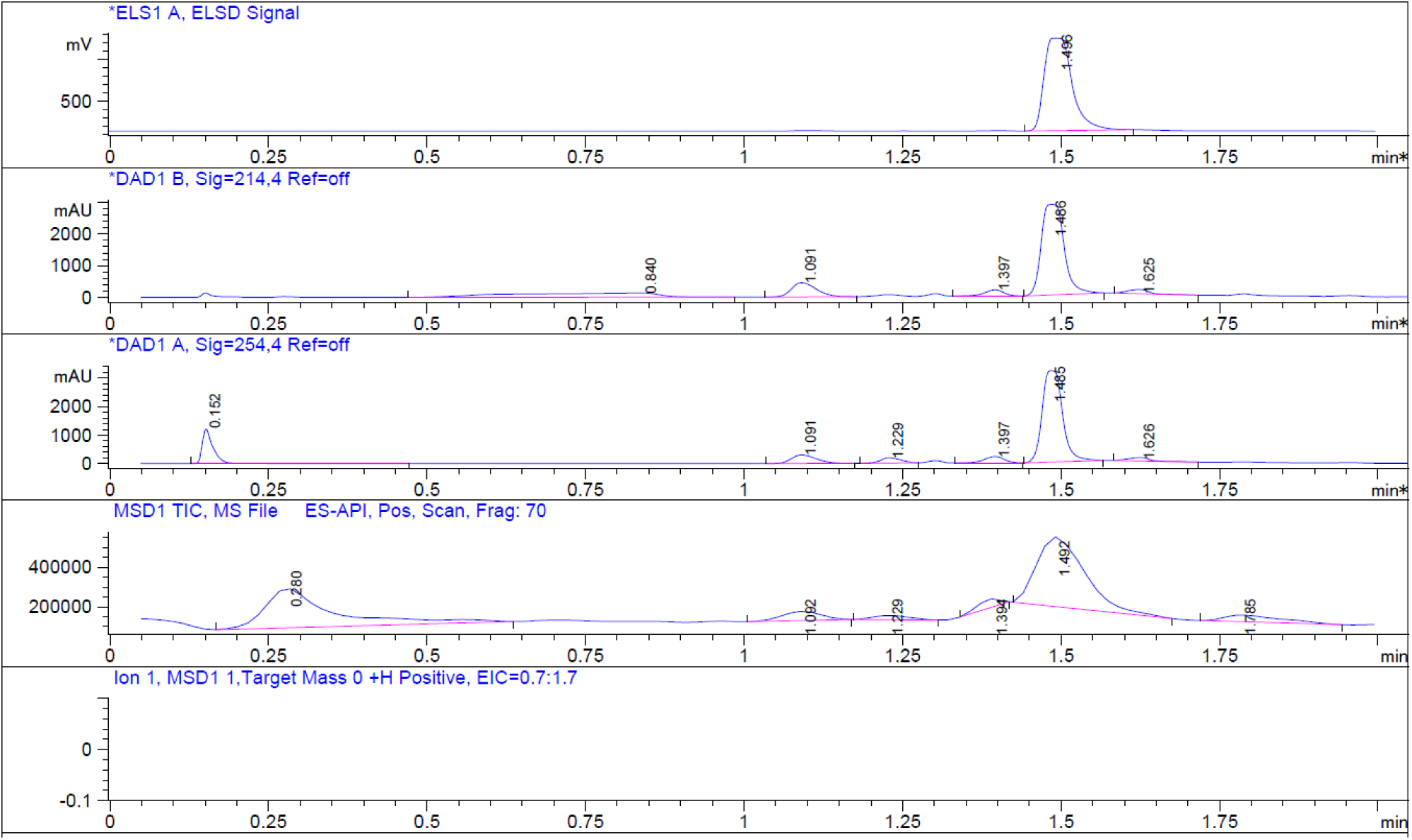

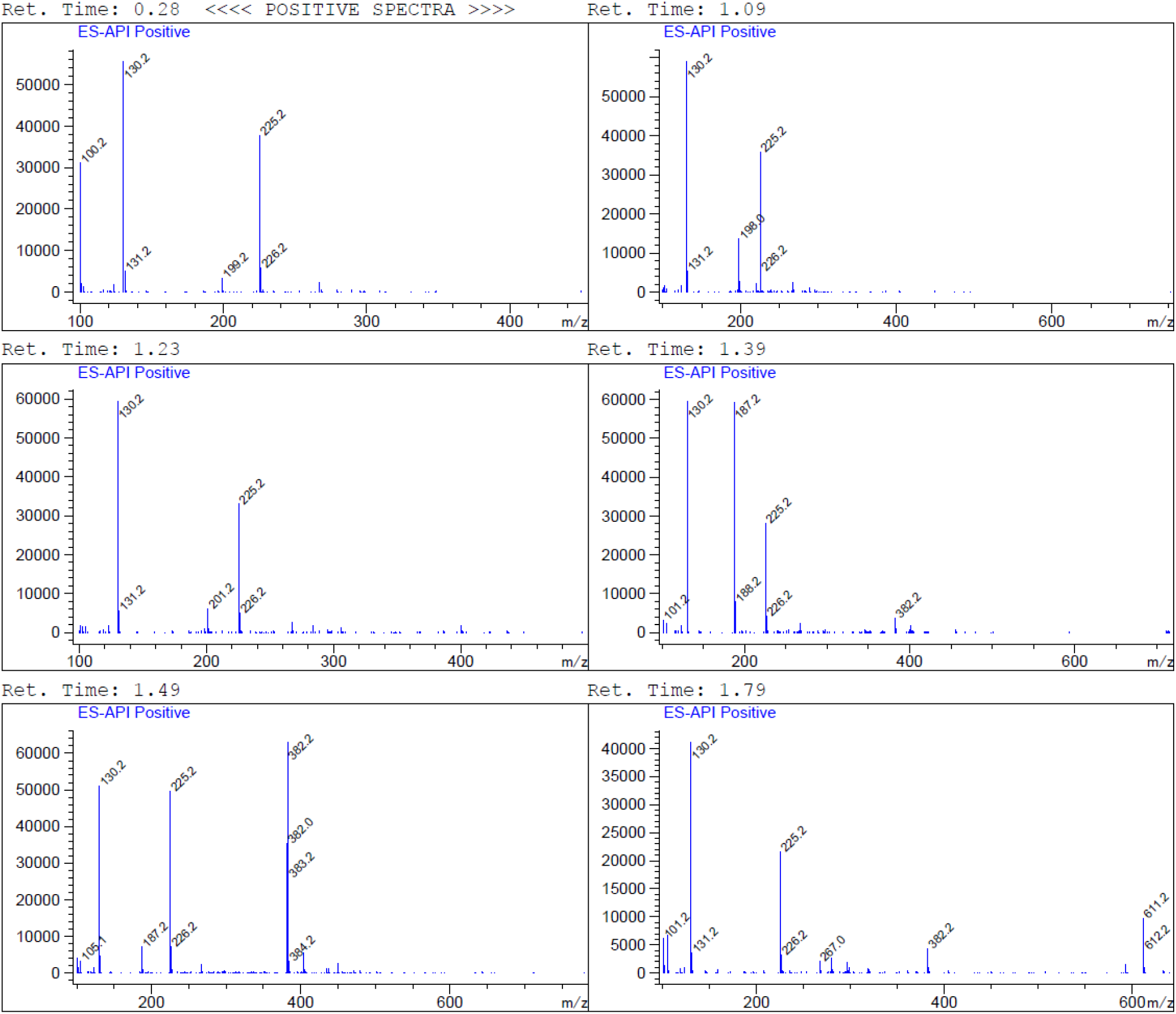

#### (5) Synthesis of compound 9

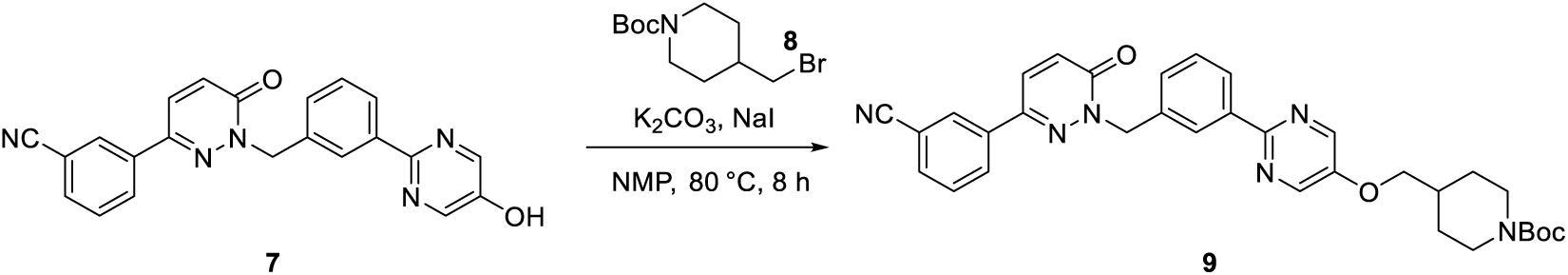

To a suspension of **compound 7** (16.6 g, 43.52 mmol) in NMP (200 mL) were added NaI (0.65 g, 4.35 mmol), K2CO3 (11.75 g, 85.04 mmol) and compound 8 (16.95 g, 60.93 mmol). The reaction was stirred at 80 °C for 8 hours. LCMS showed the reaction was complete. The reaction was quenched by water (800 mL). The mixture was filtered. The wet filter cake was recrystallized from EtOAc (200 mL) to give **compound 9** as an off-white solid (11 g, crude).

LCMS: m/z = 523.2 [M-55] ^+^, Rt = 1.885 min, purity: 91.46% (214nm)

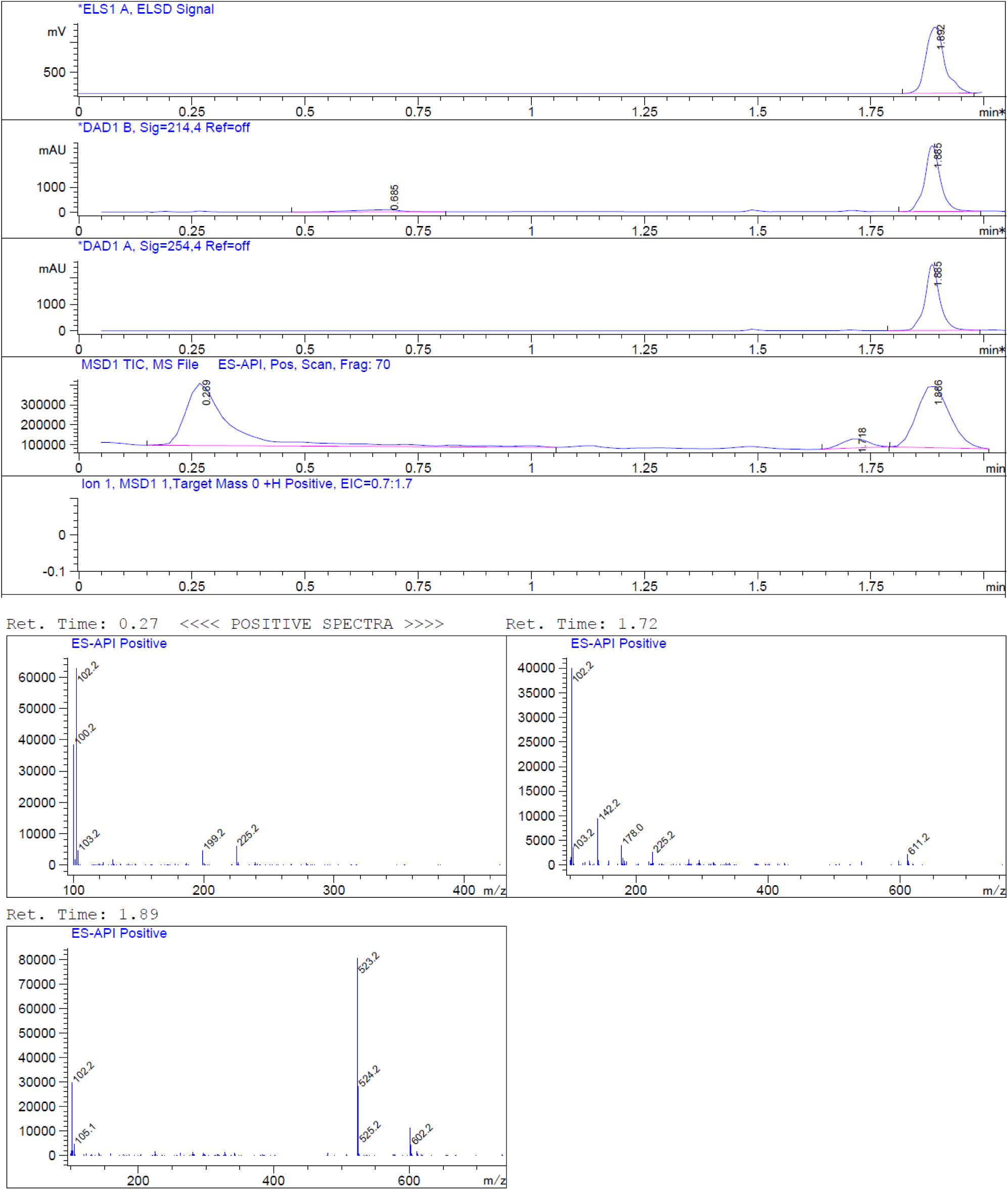

#### (6) Synthesis of compound 10

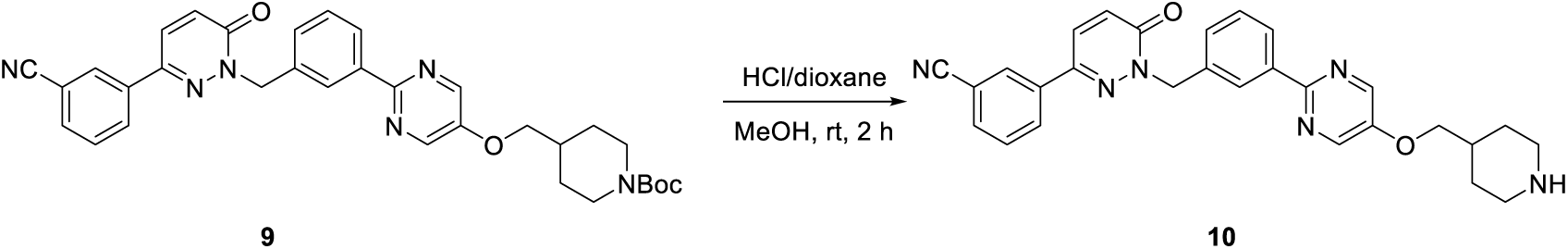

To a solution of **compound 9** (4 g, 6.91 mmol) in DCM (40 mL) was added dropwise TFA (10 mL). The resulting solution was stirred at room temperature for 2 hours. LCMS showed the reaction was complete. The mixture was concentrated in vacuum to give the crude **compound 10** (4 g, crude), which was used for the next step without further purification.

LCMS: m/z = 479.2 [M+H] ^+^, Rt = 1.373 min, purity: 52.07% (214nm)

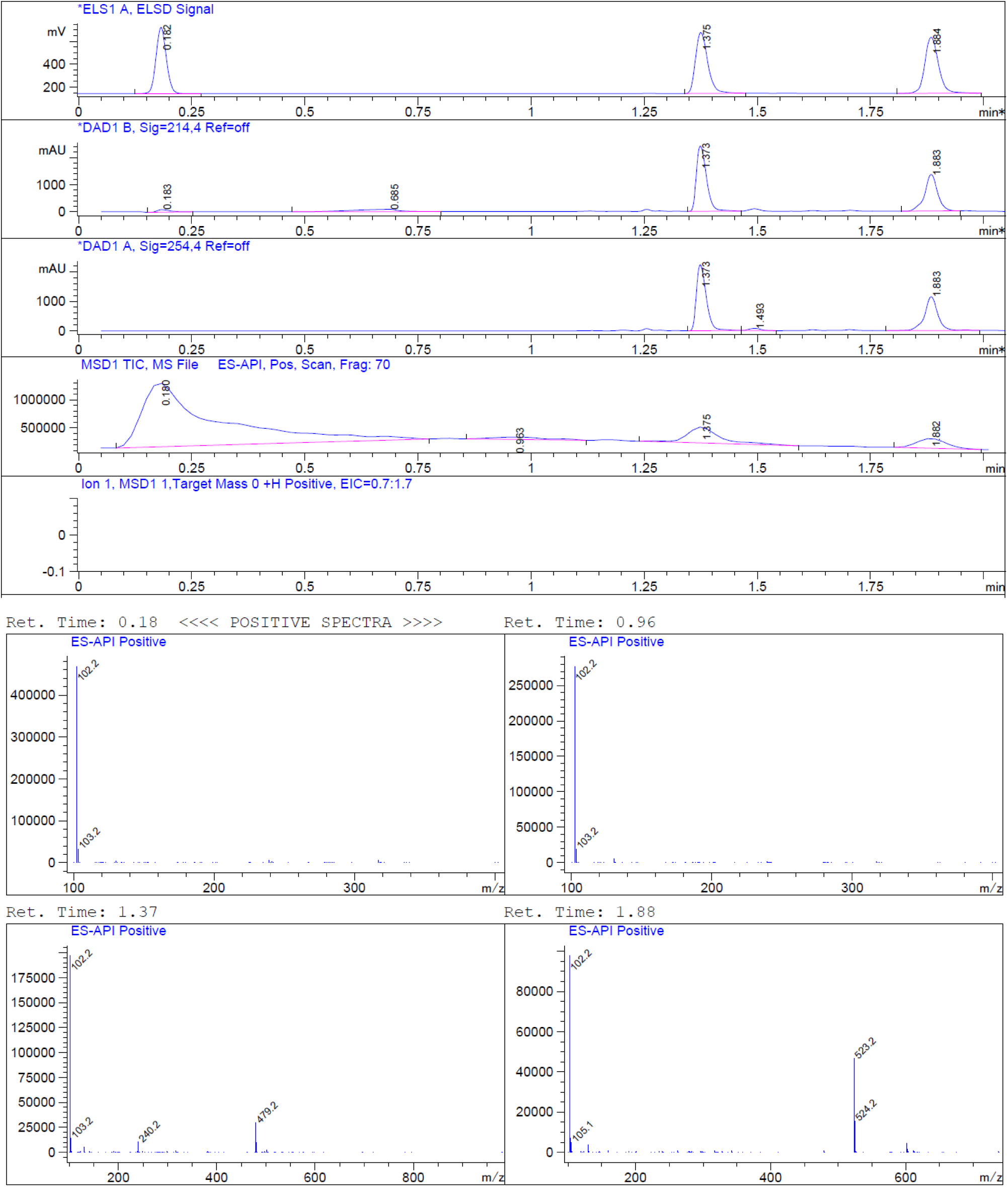

#### (7) Synthesis of compound 12

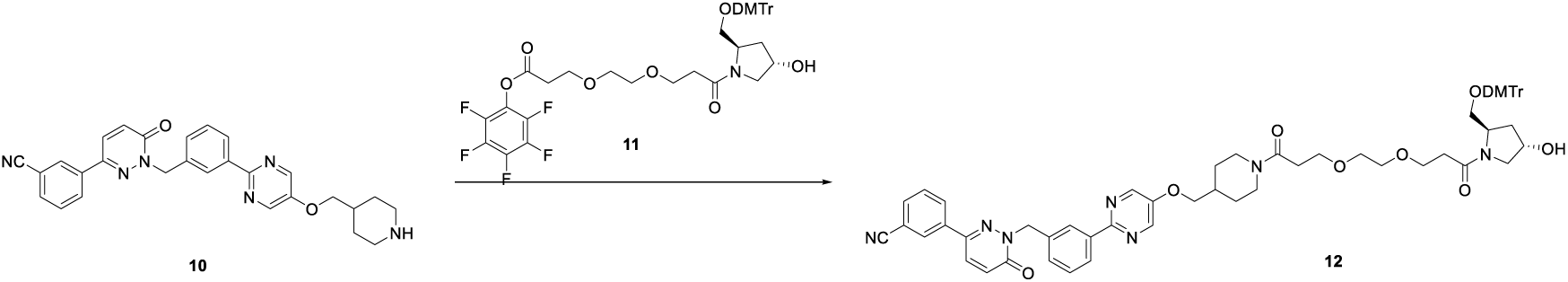

To a solution of **compound 10** (1.6 g, 2.06 mmol) in ACN (20 mL) was added DIEA (2.66 g, 20.6 mmol) under nitrogen protected and stirred for 5 min at ice-bath. **The compound 11** (0.99 g, 2.06 mmol in 10 mL ACN) was added to the mixture dropwise and stirred for about 2 hours at room temperature. LCMS showed the reaction was completed. The mixture was filtered. The filtrate was concentrated in vacuum. The reside was purified by flash column chromatograph (eluted with MeOH: DCM = 10%) to give **compound 12** (2.2 g, 99% yield) as a white solid.

LCMS: m/z = 766.4 [M-DMTr] ^+^, Rt = 1.447 min, purity:79.22% (214 nm)

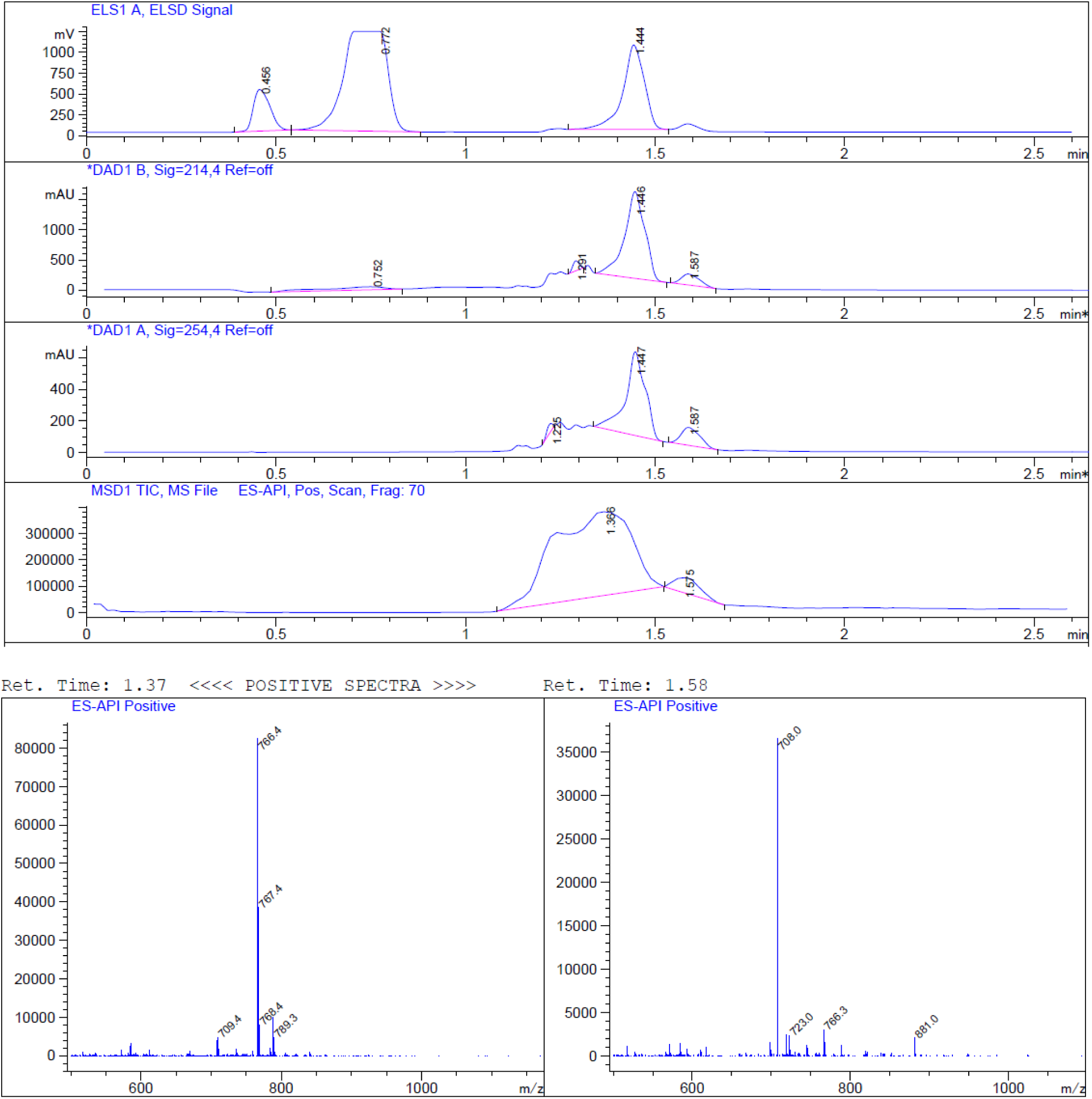

#### (8) Synthesis of compound 13

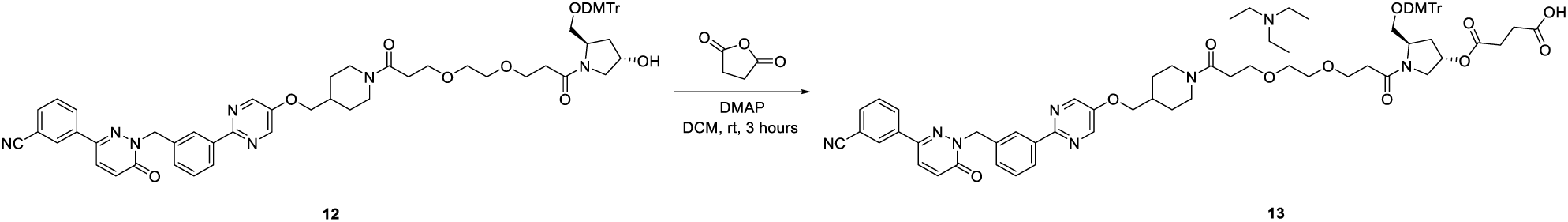

To a solution of **compound 12** (1.7 g, 1.59 mmol) in DCM (30 mL) were added succinic anhydride (317.5 mg, 3.18 mmol) and DMAP (343.3 mg, 3.18 mmol) under nitrogen protected. Then stirred for about 3 hours at room temperature. LCMS showed the reaction was complete. The mixture was concentrated in vacuum. The residue was purified by reversed phase column chromatography to give **compound 13** (910 mg, 49.00 % yield) as a white solid LCMS: m/z = 1168.5 [M+Na] ^+^, Rt = 1.689 min, purity: 100% (214 nm)

**^1^H NMR (400 MHz, DMSO-*d_6_*)** *δ* 8.63 (s, 2H), 8.42 - 8.34 (m, 2H), 8.28 - 8.14 (m, 3H), 7.97 - 7.89 (m, 1H), 7.74 - 7.67 (m, 1H), 7.51 - 7.44 (m, 2H), 7.30 - 7.28 (m, 4H), 7.24 - 7.13 (m, 6H), 6.98 - 6.74 (m, 4H), 5.44 (s, 2H), 5.37 - 5.20 (m, 1H), 4.48 - 4.35 (m, 1H), 4.29 - 4.12 (m, 1H), 4.10 - 4.00 (m, 2H), 3.94 - 3.77 (m, 2H), 3.72 (s, 6H), 3.66 - 3.54 (m, 5H), 3.51 - 3.42 (m, 5H), 3.22 - 3.17 (m, 1H), 3.13 - 2.91 (m, 3H), 2.59 - 2.54 (m, 2H), 2.49 - 2.41 (m, 10H), 2.27 - 2.17 (m, 1H), 2.12 - 1.98 (m, 2H), 1.86 -1.70 (m, 2H), 1.30 - 1.05 (m, 2H), 0.98 - 0.90 (m, 6H).

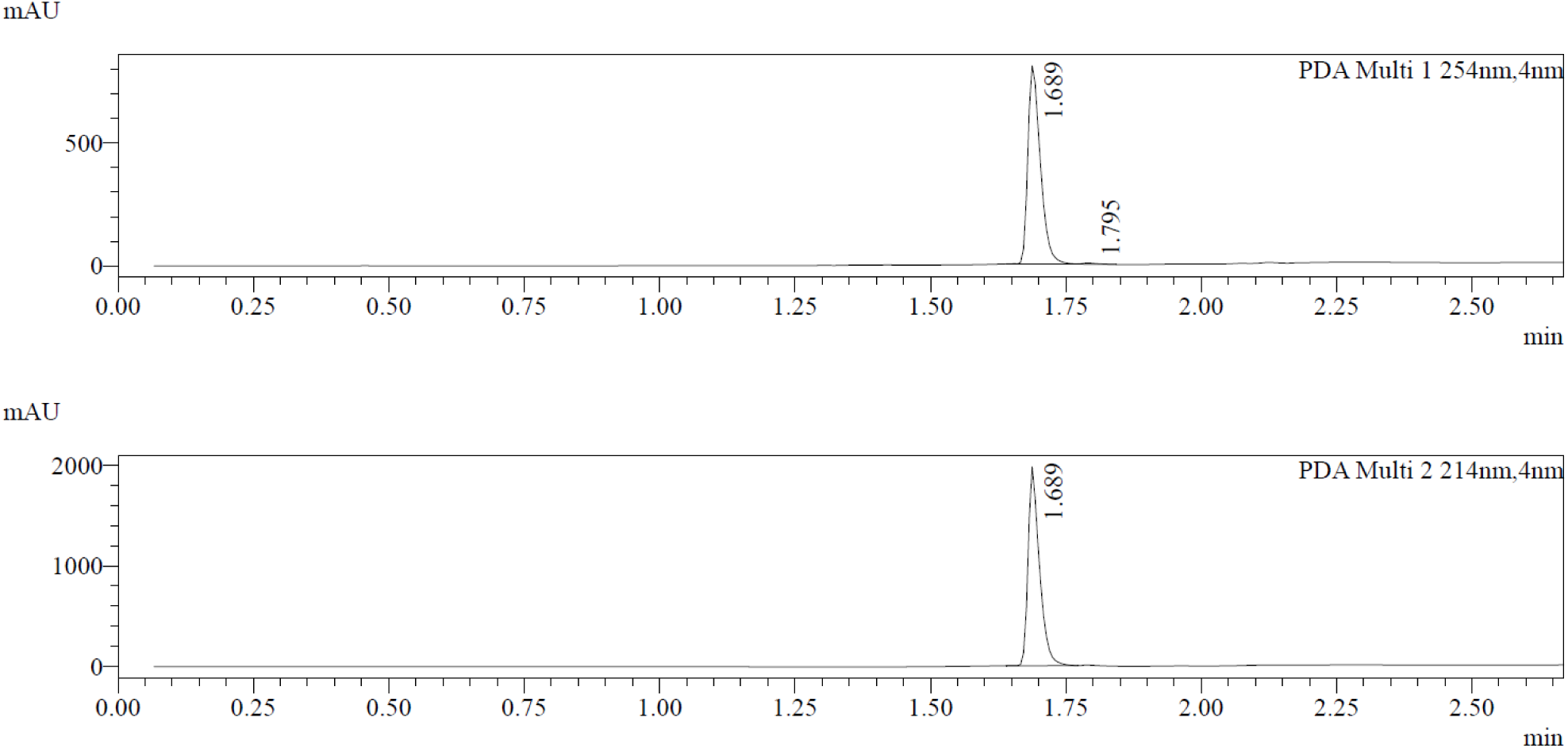

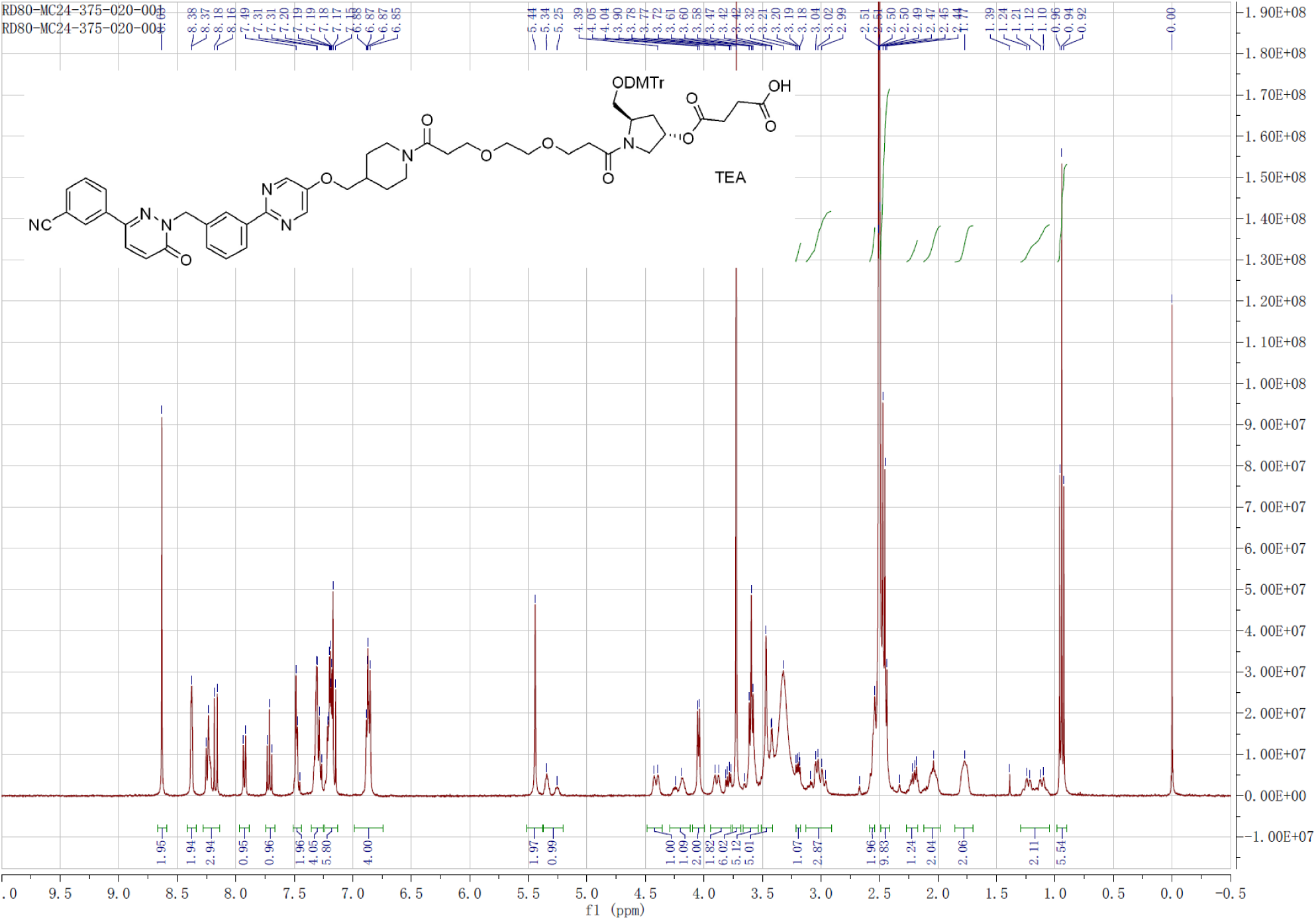

#### (9) Synthesis of compound C

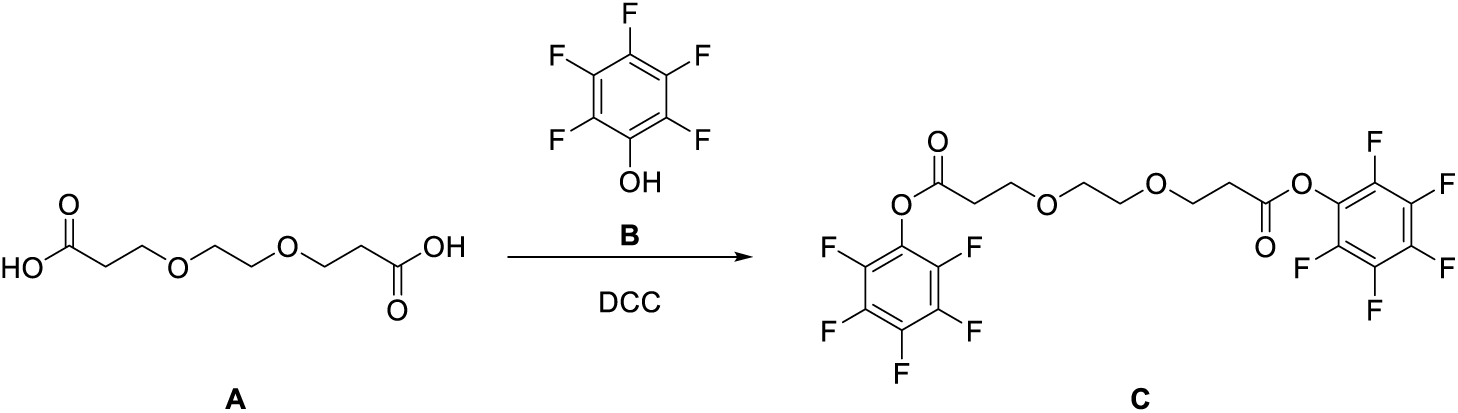

To a solution of **compound A** (4.5 g, 21.8 mmol) and **compound B** (10 g, 54.5 mmol) in DCM (45 mL) was added DCC (13.3 g, 65.4 mmol) at 0 °C. The mixture was stirred under Ar atmosphere at room temperature at 16 hours. LCMS showed the reaction was completed. The mixture was filtered. The filtrate was concentrated in vacuum. The residue was purified by flash column chromatograph (eluted with EtOAc: PE = 25%) to give **compound C** (7.5 g, 64% yield) as a white solid.

LCMS: NO MS, Rt= 1.987 min, purity: 83.591% (214 nm).

**1H NMR (400 MHz, CDCl3)** *δ* 3.90 - 3.87 (m, 4H), 3.69 - 3.67 (m, 4H), 2.96 - 2.93 (m, 4H).

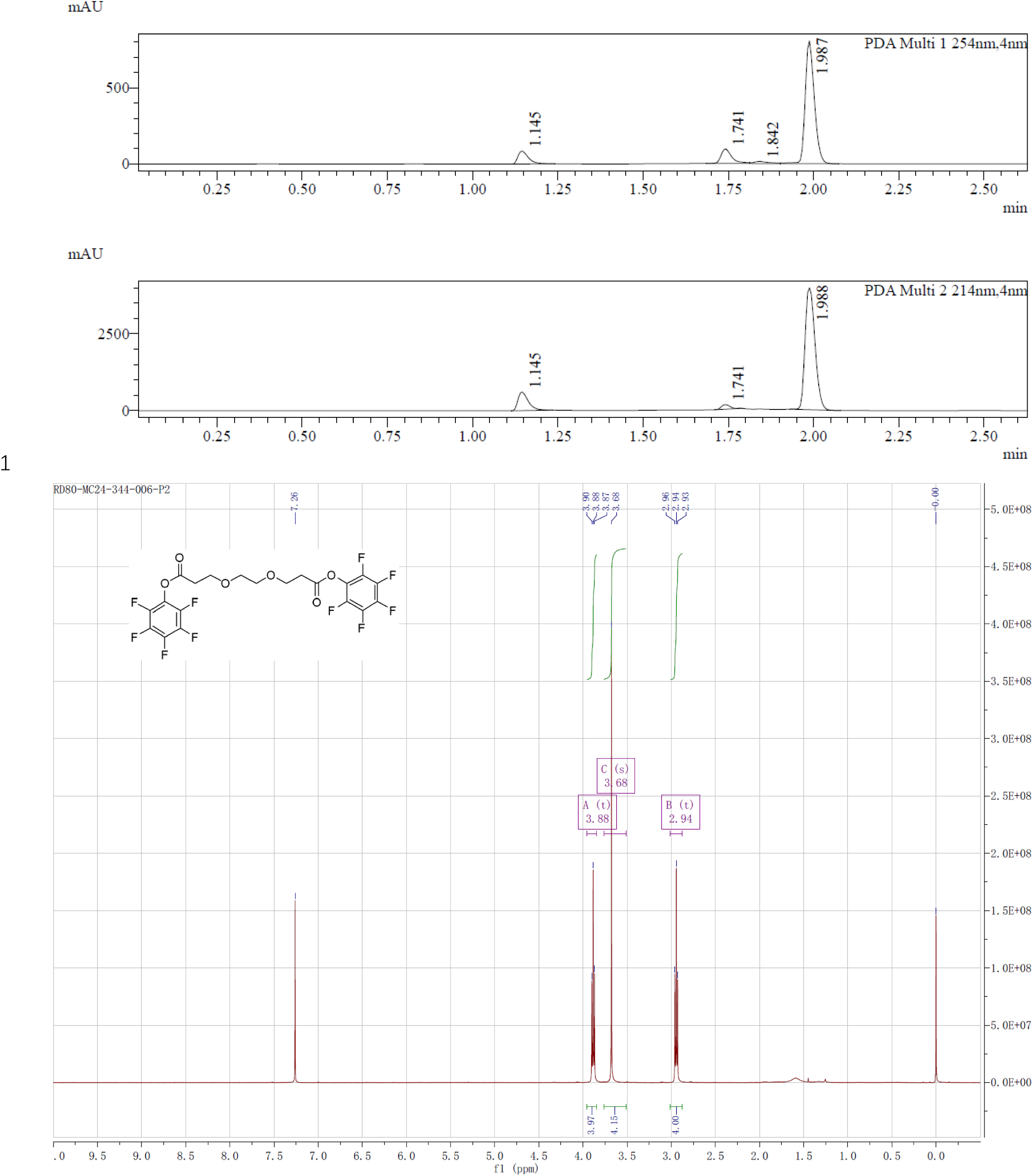

#### (10) Synthesis of compound 11

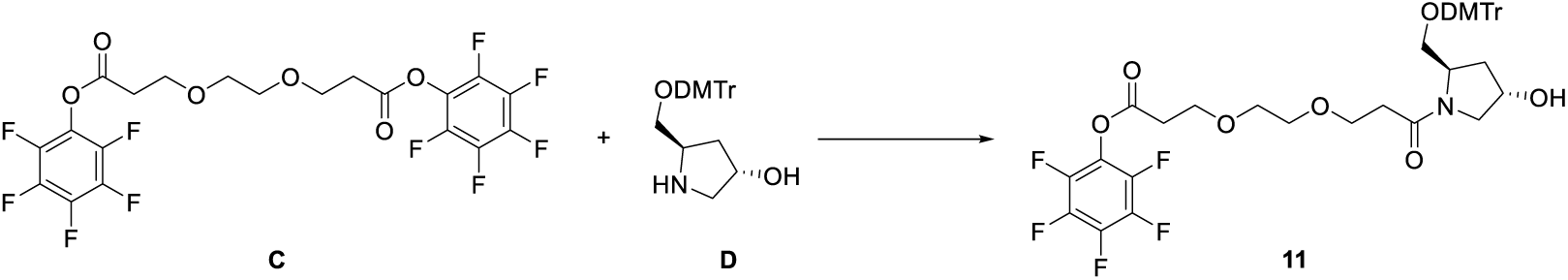

To a solution of **compound C** (6.3 g, 11.71 mmol) in ACN (20 mL) was added DIEA (15.11 g, 117.1 mmol) under N2. The **compound D** (2.46 g, 5.855 mmol in 10 mL ACN) was added dropwise to the mixture. The reaction was stirred at room temperature for 2 hours. LCMS showed the reaction was completed. The organic layers were concentrated in vacuum. The residue was purified by flash column chromatograph (eluted with EA: PE=30 %) to give **compound 11** (1.6 g, 33.34% yield) as a white solid.

LCMS: m/z = 796.2 [M+H] ^+^, Rt = 1.52min, purity: 60.68% (214 nm)

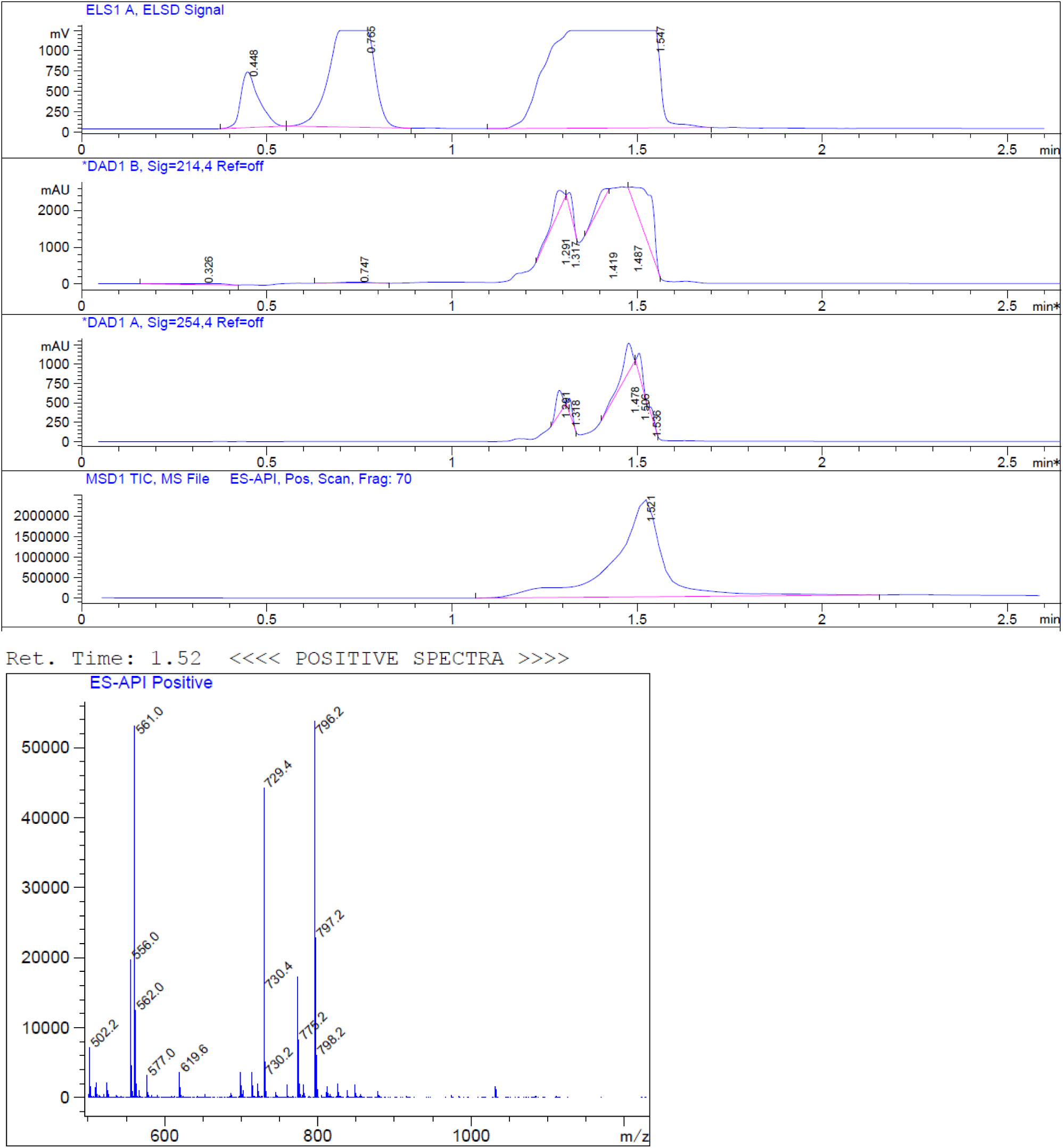

#### (11) Synthesis of SAPT8-Tep-3-CPG

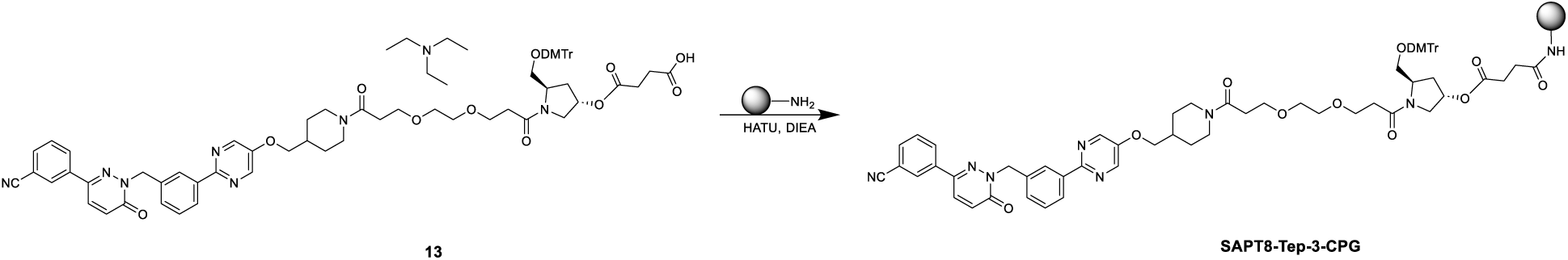

To a solution of **compound 13** (300 mg, 236.3 μmol) in ACN (18.0 mL) were added HATU (120 mg, 315.6 μmol), DIEA (120 μL), lcaa-CPG (1000 Å, 1500 mg) respectively at room temperature and oscillates for 12 hours. After the reaction was completed, CPG was washed with ACN, CAP A (acetic anhydride: tetrahydrofuran= 1:9, v/v, 12.0 mL) and CAP B (N-methylimidazole: pyridine: acetonitrile= 15:10:75, v/v/v, 12.0 mL) was added, the mixture was oscillated at room temperature for 1 hour. Then the mixture was filtered and washed with ACN (180 mL) for 3 times. **SAPT8-Tep-3-CPG** was obtained after freeze-drying of the mixture as a white powder (1500 mg).

#### (12) Synthesis of SAPT8-Tep-3

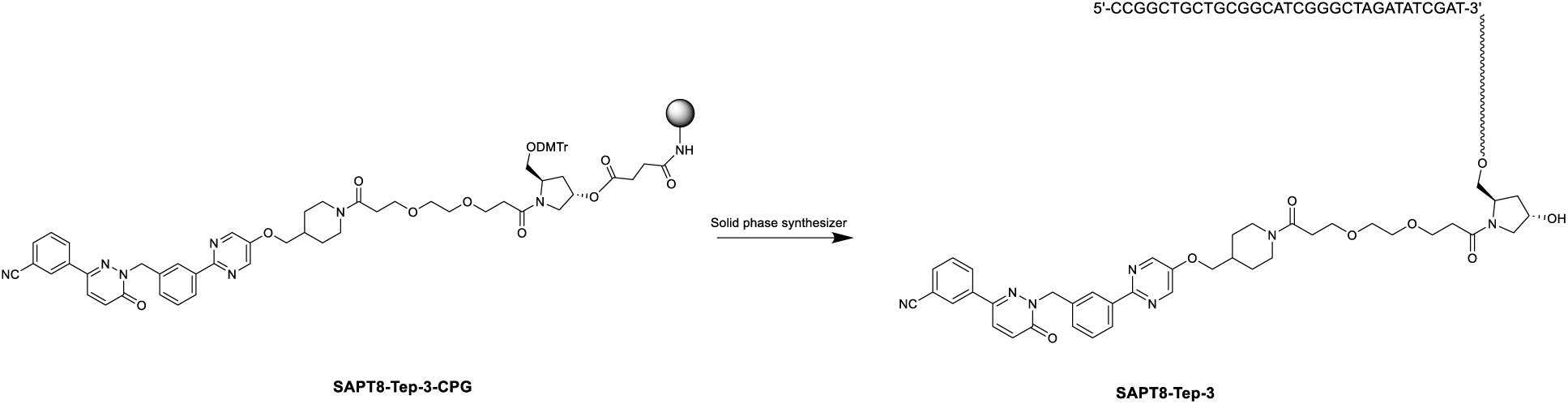

**SAPT8-Tep-3-CPG** in synthesis columns (60 mg*24) and synthesize by K-A H-8 Solid phase synthesizer. Solid phase synthesis involves four steps: Detritylation, Coupling, Capping and Oxidation. After reaction was done, CPG of each synthesis column was added 1.5 mL aqueous ammonia and heated in oven at 65 °C for 16 hours. Then collected the supernatant and washed with water (1 mL*3). The combined crude was purified by RP-IP-HPLC (WATERS 2489, 3767) (**Column**: XBridge BET C18 2.5μm. **RP-IP-HPLC method**: Mobile phase A: 2 % HFIP +0.1 % TEA in DD water, Mobile phase B: MeOH) to afford **SAPT8-Tep-3** as a white powder (43.65 mg, purity = 99.123%).

UPLC-MS (WATERS ACQUITY PREMIER): **SAPT8-Tep-3-UPLC**, m/z = 10685.27060 [M]-(deconvolution); tR = 8.079 min (260 nm). Mass error <50 ppm.

HPLC (SPT-M40): **SAPT8-Tep-3-HPLC**, *t*_R_ = 12.473 min (260 nm), purity: 99.123%.

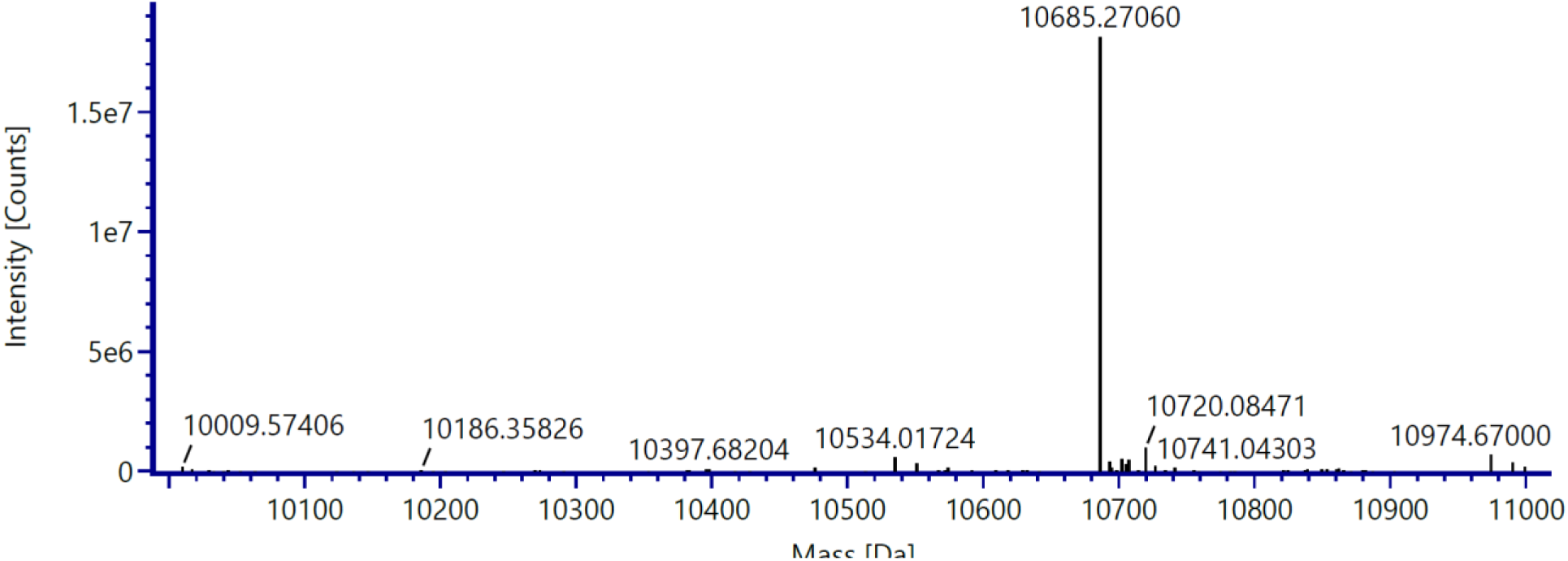

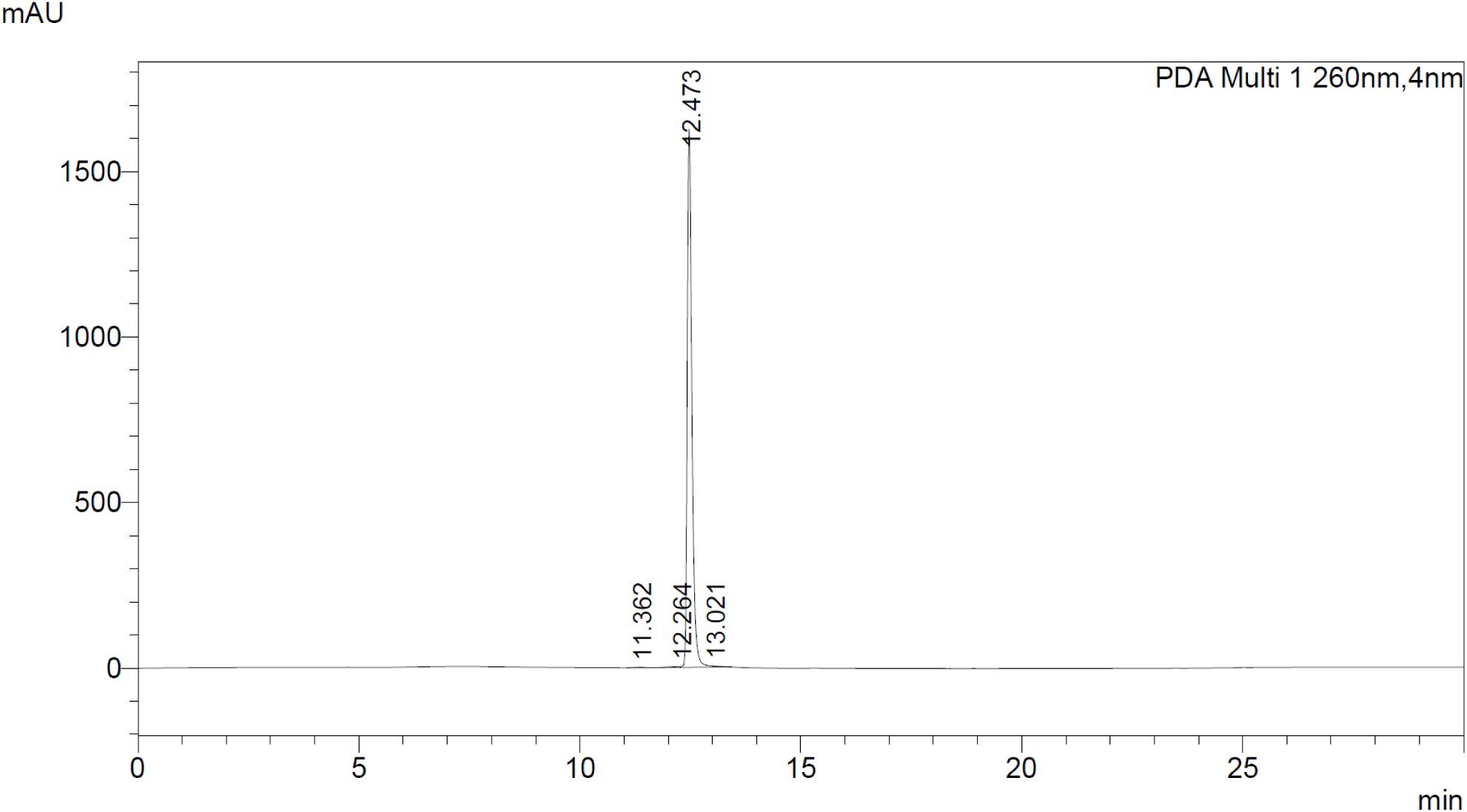

